# Novel Computational Method to Define RNA PSRs Explains Influenza A Virus Nucleotide Conservation

**DOI:** 10.1101/494336

**Authors:** Andrey Chursov, Nathan Fridlyand, Albert A. Sufianov, Oleg I. Kiselev, Irina Baranovskaya, Andrey Vasin, Jonathan W. Yewdell, Alexander Shneider

## Abstract

**ABSTRACT:** RNA molecules often fold into evolutionarily selected functional structures. Yet, the literature offers neither a satisfactory definition for “structured RNA regions”, nor a computational method to accurately identify such regions. Here, we define structured RNA regions based on the premise that both stems and loops in functional RNA structures should be conserved among RNA molecules sharing high sequence homology. In addition, we present a computational approach to identify RNA regions possessing evolutionarily conserved secondary structures, RNA ISRAEU (RNA Identification of Structured Regions As Evolutionary Unchanged). Applying this method to H1N1 influenza mRNAs revealed previously unknown structured RNA regions that are potentially essential for viral replication and/or propagation. Evolutionary conservation of RNA structural elements may explain, in part, why mutations in some nucleotide positions within influenza mRNAs occur significantly more often than in others. We found that mutations occurring in conserved nucleotide positions may be more disruptive for structured RNA regions than single nucleotide polymorphisms in positions that are more prone to changes. Finally, we predicted computationally a previously unknown stem-loop structure and demonstrated that oligonucleotides complementing the stem (but not the loop or unrelated sequences) reduce viral replication *in vitro.* These results contribute to understanding influenza A virus evolution and can be applied to rational design of attenuated vaccines and/or drug designs based on disrupting conserved RNA structural elements.

*AUTHOR SUMMARY:* RNA structures play key biological roles. However, the literature offers neither a satisfactory definition for “structured RNA regions” nor the computational methodology to identify such regions. We define structured RNA regions based on the premise that functionally relevant RNA structures should be evolutionarily conserved, and devise a computational method to identify RNA regions possessing evolutionarily conserved secondary structural elements. Applying this method to influenza virus mRNAs of pandemic and seasonal H1N1 influenza A virus generated Predicted Structured Regions (PSRs), which were previously unknown. This explains the previously mysterious sequence conservation among evolving influenza strains. Also, we have experimentally supported existence of a computationally predicted stem-loop structure predicted computationally. Our approach may be useful in designing live attenuated influenza vaccines and/or anti-viral drugs based on disrupting necessary conserved RNA structures.

## INTRODUCTION

The biological functions of RNA secondary structures and their evolutionary impact is a topic of great interest and importance (1-11). Since 1990s, conceptually novel computational approaches have been appearing to analyze RNA shapes. Multiple copies of the same RNA molecule fold into different coexisting conformations constituting an ensemble of RNA structures. The same nucleotide within the RNA may be coupled via W-C bonds in some conformations (being a part of a stem) while remaining uncoupled in others (i.e. belonging to a loop). If one analyzes the entire ensemble, they can attribute to each nucleotide within an RNA sequence its base pairing probability. This probability value reflects i) the percentage of RNA structures which have this particular nucleotide W-C bonded (although in different structures it may be bonded to a different coupling partners), and ii) the likelihood of each RNA structure within the ensemble of all possible conformations that is based on the structure’s free energy. Hence, if an RNA molecule contains X nucleotides, a series of X numbers ranging from 0 to 1 can be estimated such that each number reflects the probability of a nucleotide in this position to be coupled.

Viruses represent an excellent system for studying RNA structural biology based on several factors, including the high abundance of viral RNAs, the high mutation rate of many viruses, the ease of selecting conditional mutants, and the large number of closely related strains available for sequence conservation analysis.

Mutations disrupting influenza A virus RNA secondary structures dramatically reduce the levels of gene expression (12). We previously reported that influenza A virus, a negative strand RNA virus with a segmented genome, possesses clusters of nucleotides that significantly change their base pairing probabilities with temperature elevation (13). This suggests that local structures dispersed between non-structured RNA regions (nonPSRs) are evolutionarily selected.

Previous attempts to define “structured RNA regions” were aimed either at finding regions possessing the most stable secondary structure predicted by minimum free energy (MFE) of based paired sequences (14-16), or predicting a consensus secondary structure based on a given multiple-sequence alignment, which can be inferred either by means of energy-directed folding or using a phylogenetic stochastic context-free grammar model (17-21). The latter approach works well on short non-coding RNAs, which usually have one predicted stable structure, but its accuracy drops significantly with the increasing RNA length (22). In addition, mRNAs are typically less structured than non-coding RNAs, since structures interfere with translation by ribosomes (23). At the same time, the former approach misinterprets “PSRs” as stems possessing abnormally many W-C coupled nucleotides; i.e. it would view an RNA structure possessing evolutionarily conserved loops (where the nucleotides are not paired) as an unstructured element. The following example demonstrates how misleading interchangeable use of the words “structured” and “paired” is: a cloverleaf-like secondary structure may serve an indispensable biological function and be conserved in every strain of some organism, despite the fact that it may have fewer paired nucleotides than a simple stem (see Figure 1). Also, such approach may introduce systemic bias typically identifying mRNAs as less structured than non-coding RNAs, since excessive abundance of W-C pairing may interfere with translation by ribosomes (23).

**Figure 1:**
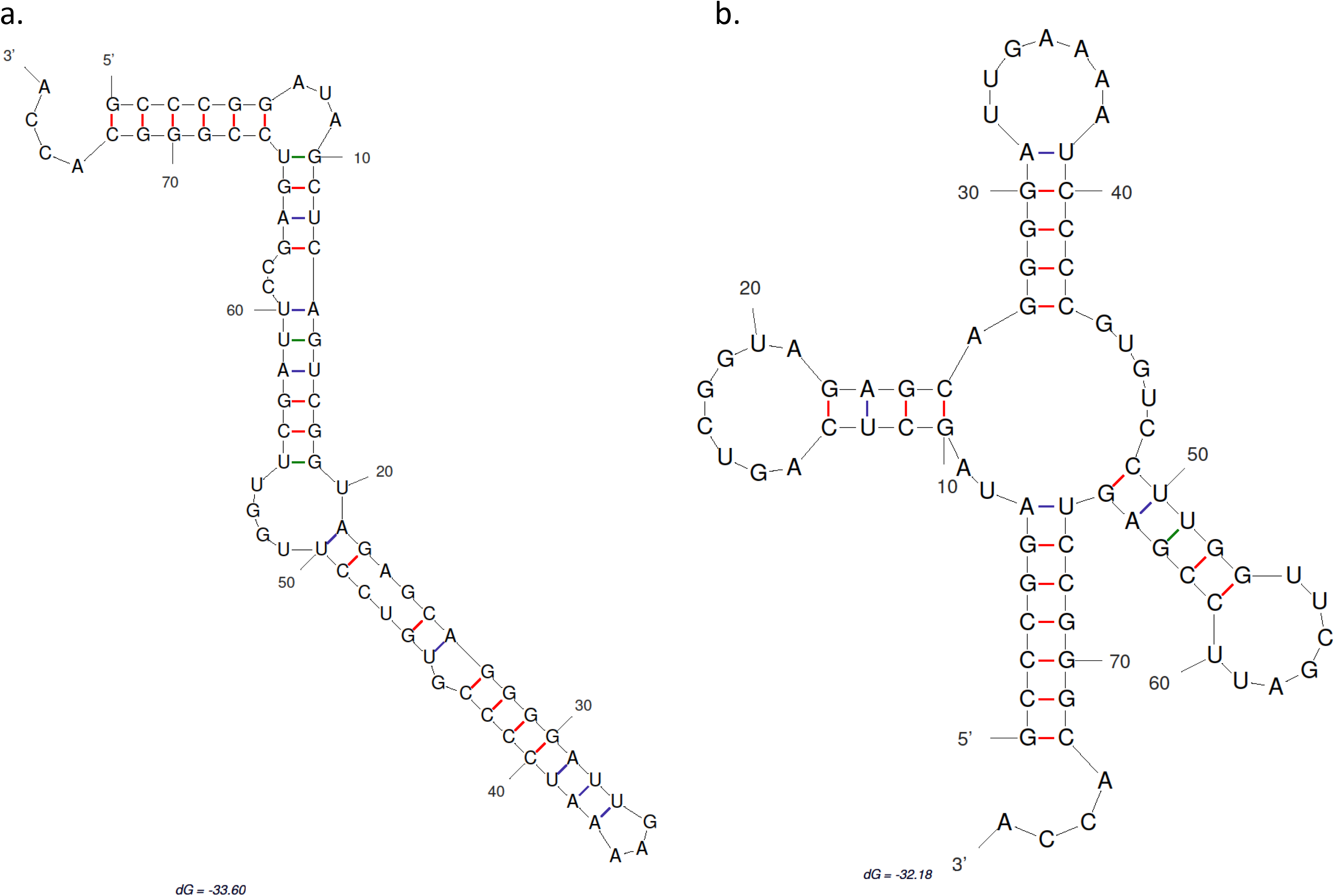
Computationally predicted by mfold 3.6 (71) (a) optimal and (b) one of many suboptimal secondary structures of tRNA. Mfold was used with the default energy parameters including the folding temperature fixed at 37°C. Despite the fact that the left structure contains more base pairs, the right structure is functional and evolutionarily conserved.

In addition, any particular RNA sequence beyond certain length clearly can fold into stable alternative states with energies being somehow different from the global minimum (24-26); and several alternative RNA conformations coexist at equilibrium (27). Some of these RNA structures may be present in multiple RNAs, especially homologous ones, while others would exist for the particular sequence only. One can assume that evolutionary conservation of the RNA shape may be indicative of its biological function. Current approaches relied upon analysis of a single sequence cannot differentiate an evolutionarily conserved structural element from an RNA shape that is energetically favorable only in a particular strain. Thus, future progress in the field requires a new methodology. Addressing these problems and proposing a computational methodology free of these shortcuts is the main aim of this paper.

## Results

### A Novel Quantitative Definition of Structured RNA Regions

Here, we present a quantitative definition of a structured RNA region (PSR) that is equally useful to predicting both, stems and loops as structured regions based on their evolutionary conservation, and a new computational method for identifying those regions. The method is robust to random RNA shapes present in a particular sequence but not selected and conserved evolutionarily. We call this method RNA ISRAEU (RNA Identification of Structured Regions As Evolutionary Unchanged).

As the first step in this method, we created a non-redundant dataset of sequences for each RNA of interest constituted of highly homologous RNAs of the same length, and built multiple sequence alignments. For the second step, the probability of every nucleotide to be paired was calculated for each RNA sequence in the dataset. Third, we substituted every nucleotide in the multiple alignment, with the nucleotide’s pairing probability, thus aligning pairing probabilities by nucleotide position. We took probabilities to be paired for all the 1st nucleotides in each RNA sequence and grouped them together; for all 2^nd^ nucleotides; for all N^th^ nucleotides. Thus, if we have X RNA sequences each constituted of Y nucleotides, we create Y sets of numbers ranging from 0 to 1; each set contains X numbers. Standard deviation was computed for each set of probability values corresponding to every position within the RNA. We proposed that such standard deviations be used as a measure of structural conservation in a specific position. If the standard deviation for a particular position within the RNA dataset was small, the probability of a nucleotide to be in a double-stranded conformation did not vary substantially across the entire dataset of aligned mRNAs. We call such positions “structure-conserved”. In contrast, if the standard deviation was high, the probability of a nucleotide to be paired changed vastly from strain to strain, a position was called “structure-variable”. The mean probability at a particular position did not matter to the position classification.

We called regions within RNA sequences formed by consecutive structure-conserved positions “Predicted Structured Regions” (PSRs), while regions predominantly formed by structure-variable positions were called “non-structured” (nonPSR). Apparently, such definition is stem-loop agnostic. A stretch of nucleotide positions, which demonstrate high probability of being paired across the entire dataset of RNAs, may form a functionally important and evolutionarily conserved stem. Similarly, a batch of nucleotide positions with low base pairing probabilities, which repeats itself across all RNAs in the dataset may form a functionally important and evolutionarily conserved loop. Still, further analysis is necessary to confirm both the stems and the loops. In all cases, within a PSR, the probability of each nucleotide to be in a double-stranded conformation does not vary significantly across the entire dataset of aligned RNAs and these positions are structure-conserved positions.

### Identification of Structured RNA Regions in H1N1 Influenza Virus mRNAs

We applied RNA ISRAEU to predict evolutionarily conserved RNA structures of influenza A virus (IAV) (28, 29).

We selected sequences encoded by the complete genomes of 107 pre-pandemic (1999 to 2009) and 173 pandemic (post-2009) human H1N1 strains (Supplementary Table 2). In 2009, a swine IAV strain was introduced into man and rapidly replaced the circulating strains. All mRNAs selected for a given gene were of the same length. For each of the 10 major viral mRNAs, we predicted structured regions and calculated sequence variation. Profiles for non-pandemic and pandemic NS2 mRNA are depicted in Figure 2 (profiles for other mRNAs are presented in Supplementary Figures 2-38).

**Figure 2:**
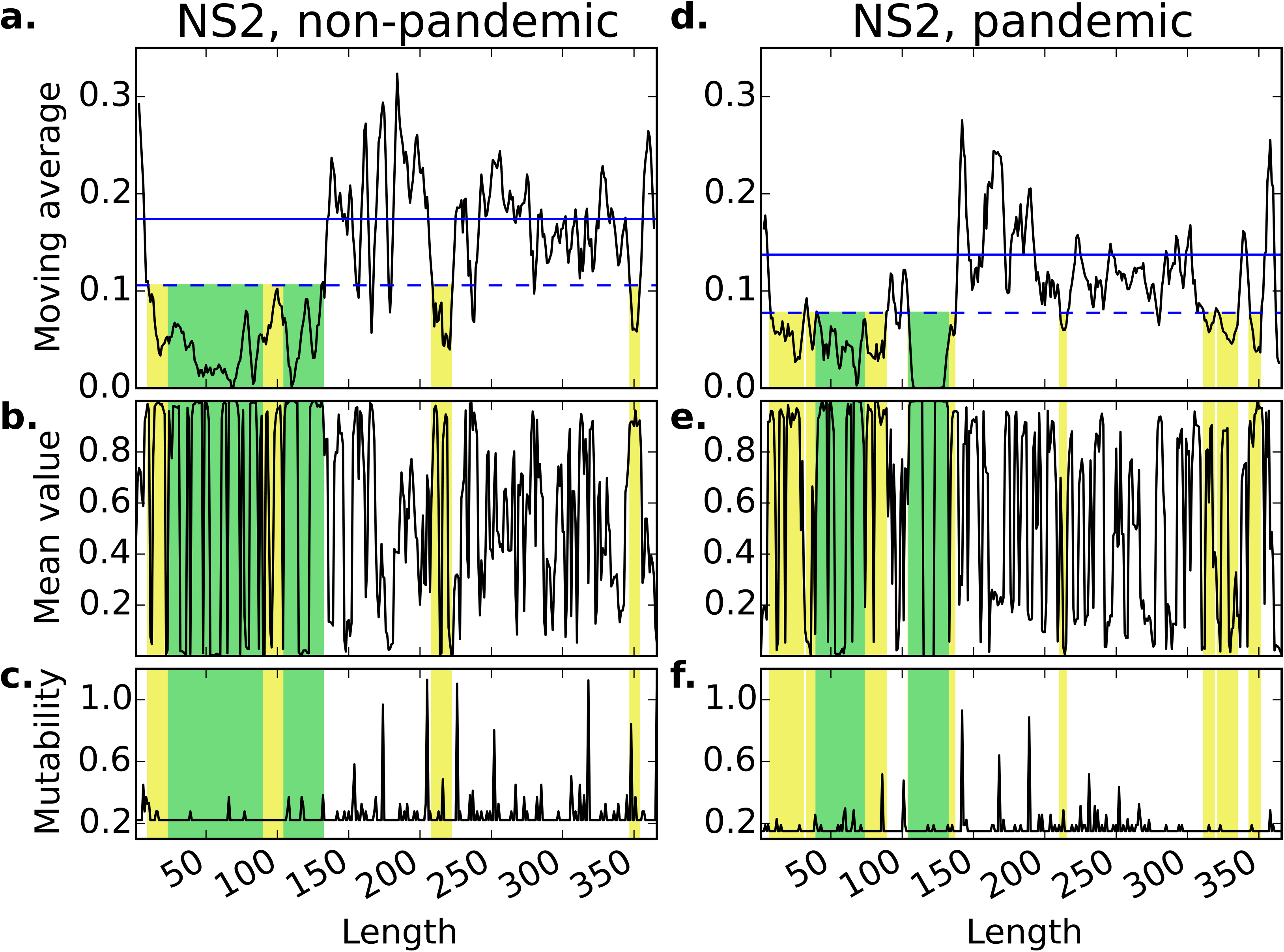
Structure variability and mutability profiles for non-pandemic ((a), (b), and (c)) and pandemic ((d), (e), and (f)) NS2 influenza mRNAs. Plots (a) and (d) demonstrate structure conservation profiles; namely, they show the moving average that was calculated by applying a sliding window approach to smooth individual fluctuations of standard deviations of nucleotide base pairing probabilities. The blue solid line demonstrates the mean level of all moving average values, and the blue dashed line demonstrates the level equal to the mean of all moving average values decreased by the standard deviation of all moving average values. In this case, the mean and the standard deviation were computed based on all moving average values from all mRNAs of a particular type (pandemic or non-pandemic) of influenza strains. According to our definition, when the moving average goes below the blue dashed line, it is a structured RNA region. Such regions are colored with either yellow or green across the plots. Plots (b) and (e) demonstrate profiles of the mean values of probabilities of nucleotide positions to be in a double-stranded conformation. If this value is close to 1, it means that in most strains in the dataset the correspondent nucleotide has a very high probability to be paired; and, if this value is close to 0, the correspondent nucleotide is very likely to be unpaired in most strains in the dataset. Plots (c) and (f) demonstrate mutability profiles for NS2 mRNAs. Mutability of every nucleotide position is computed as a value of Shannon entropy which is calculated based on frequency of every ribonucleotide in a particular position. Areas within RNA colored with yellow or green demonstrate identified structured RNA regions. Meanwhile, areas colored with green show regions in which particular secondary structure was determined.

142 and 134 PSRs were identified in non-pandemic and pandemic influenza mRNAs respectively. The length of PSRs varies from 5 to 121 nucleotides for non-pandemic and from 5 to 103 nucleotides for pandemic strains (Table 1). The number of PSRs varies from 2 for NS2 to 31 for NP (Table 1). Only a fraction of PSRs overlap between pandemic and non-pandemic strains. In some mRNAs (namely PB1, PB2, PA, HA and NA), the percentage of such non-overlapping regions is higher than 70%. The location of each PSR are presented in Supplementary Table 3.

**Table 1:** Numbers of identified structured RNA regions within influenza mRNAs.

| Gene name | Non-pandemic |  |  |  |  |  | Pandemic |  |  |  |  |  | Number of *in silico* SNPs |
| --- | --- | --- | --- | --- | --- | --- | --- | --- | --- | --- | --- | --- | --- |
|  | Total number | Length |  |  | Do not overlap with PSRs in pandemic strains |  | Total number | Length |  |  | Do not overlap with PSRs in non-pandemic strains |  |  |
|  |  | Min | Max | Median | Number | Percentage of total, % |  | Min | Max | Median | Number | Percentage of total, % |  |
| PB2 | 23 | 5 | 31 | 7 | 19 | 82.6 | 18 | 5 | 34 | 8.5 | 13 | 72.2 | 7 |
| PB1 | 15 | 5 | 32 | 7 | 13 | 86.7 | 15 | 5 | 103 | 9 | 13 | 86.7 | 7 |
| PA | 20 | 5 | 54 | 8 | 17 | 85.0 | 13 | 5 | 27 | 9 | 10 | 76.9 | 7 |
| HA | 11 | 5 | 10 | 9 | 9 | 81.8 | 21 | 5 | 24 | 8 | 19 | 90.5 | 5 |
| NP | 28 | 5 | 24 | 9 | 15 | 53.6 | 31 | 5 | 67 | 7 | 17 | 54.8 | 5 |
| NA | 7 | 5 | 18 | 10 | 6 | 85.7 | 6 | 7 | 24 | 10.5 | 5 | 83.3 | 5 |
| M1 | 22 | 5 | 57 | 10 | 10 | 45.5 | 15 | 5 | 30 | 8 | 2 | 13.3 | 2 |
| M2 | 5 | 7 | 34 | 14 | 2 | 40.0 | 5 | 5 | 43 | 9 | 3 | 60.0 | 1 |
| NS1 | 8 | 5 | 46 | 22.5 | 6 | 75.0 | 2 | 5 | 26 | 15.5 | 0 | 00.0 | 2 |
| NS2 | 3 | 7 | 121 | 14 | 0 | 00.0 | 8 | 5 | 48 | 11 | 2 | 25.0 | 1 |
| Total | 142 | 5 | 121 | 9 | 97 | 68.3 | 134 | 5 | 103 | 8 | 84 | 62.7 | 42 |

In comparing PSR profiles with the profiles of mean pairing probabilities, we found two evolutionarily conserved structural elements. One is located between positions 105 and 132 in non-pandemic NS2 mRNA (Figure 2), which contains a previously unknown predicted conserved hairpin (Figure 3). Nucleotides 105 to 114 and 123 to 132 have a strong predicted tendency to be paired while intervening nucleotides 115 to 124 have a strong tendency to be unpaired. By comparing Figures 2(b) and 2(e), one can predict that this new hairpin structure also exists in pandemic NS2 influenza mRNAs. The second novel PSR identified in non-pandemic NS2 mRNA is between positions 24 and 89 (Figures 2 and 3). In this case pandemic mRNAs contain only the PSR created by nucleotides 40 to 73.

**Figure 3:**
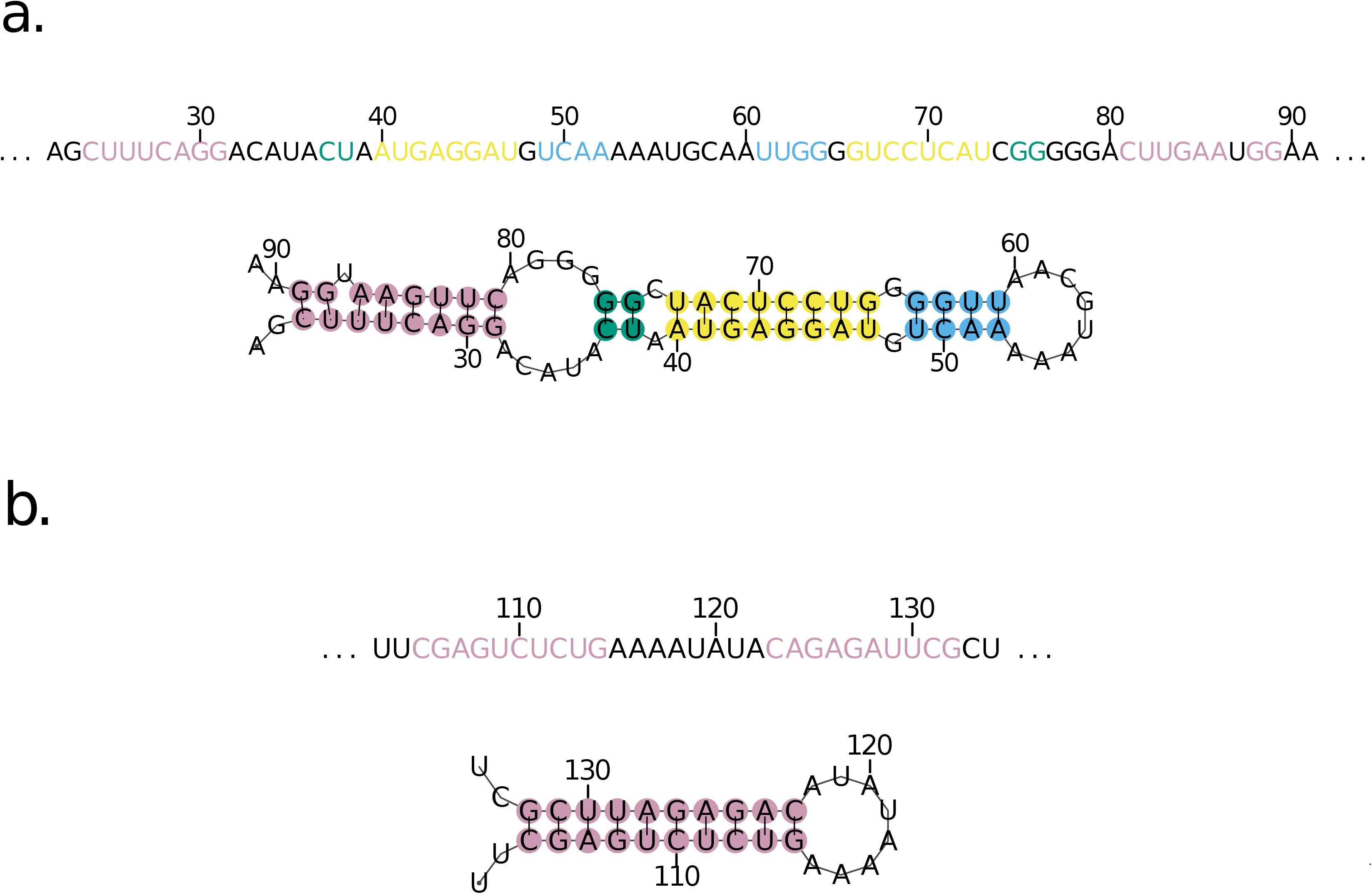
Secondary structural elements identified in the NS2 mRNA of H1N1 influenza A virus. These structural elements are evolutionarily conserved among analyzed strains. Hairpin at plot (b) was identified in both non-pandemic and pandemic H1N1 influenza. Structure shown at plot (a) exists in non-pandemic influenza virus, while pandemic mRNAs contain only part of that structure covered by nucleotide positions 40 to 73.

### Oligonucleotides complementing the stem of the newly Predicted Structured Regions interfere with *in vitro* viral replication

To test the computationally predicted RNA structured regions, we have designed oligonucleotides complementing the stem and the loop, as well as two controls of the same length. The first control did not complement any sequence within the viral or human genome while the other control bound a non-structured region adjacent to the PSR. The MDCK cell monolayer was either transfected with one of the oligonucleotides and then infected with A/California/7/09 strain or the cells were infected without prior transfection. The transfection doze was not toxic for the cells as it was proven by the cell viability assay. Twenty four hours post transfection and infection, the viral replication was assessed by developing the cell monolayer with anti-NP ELISA. We observed that only the decamer complementing the stem of the computationally predicted hairpin has significantly reduced the viral replication comparing to the controls. Neither the oligonucleotide complementing the loop, nor the two control oligos had a statistically significant effect on the *in vitro* viral replication (Figure 4).

**Figure 4:**
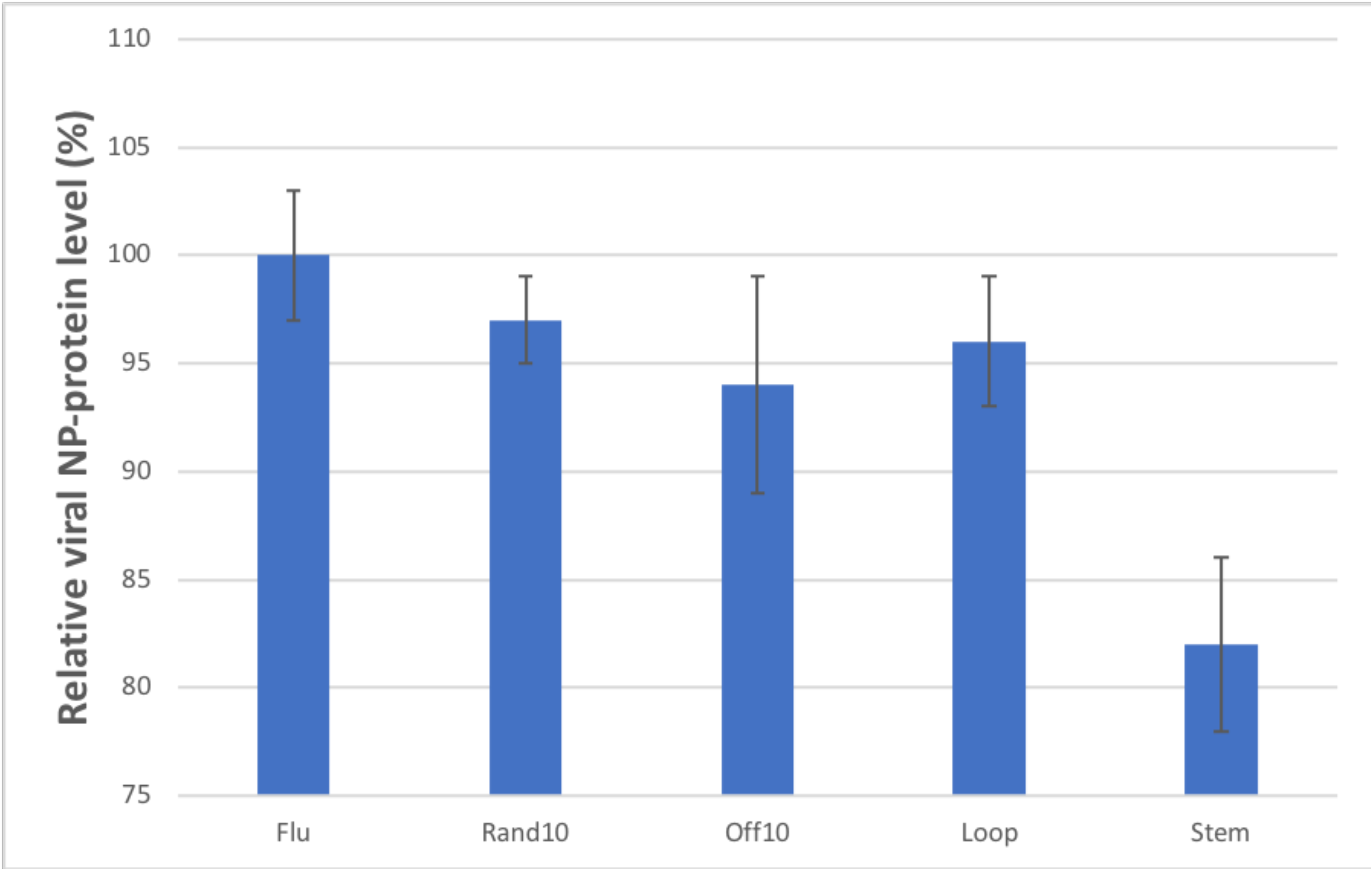
Influenza viral replication inhibition effect of antisense oligonucleotides, 24 hours post infection. P-value for the comparison between “Stem” and “Flu”is 0.0043 and is the only statistically significant difference.

### Location of the Most Mutable Positions

We distinguish between two types of positions in the influenza genome – the mutable positions which mutate quite frequently, and conserved positions. We tested if positions mutating more often cluster outside of PSRs, while conserved positions are predominantly located within the PSRs. The numbers for both types of positions in every mRNA of pandemic and non-pandemic H1N1 strain are provided in the Supplementary Table 1. Percentage of highly mutable positions in influenza mRNAs varies in a range from 7.9% in M1 to 15.5% in NA and from 6.8% in M2 to 15.8% in HA for non-pandemic and pandemic influenza strains respectively. Among the highly mutable positions from 56.1% for NS1 to 88.4% for NP and from 50.0% for M2 to 85.1% for M1 are third-codon positions for non-pandemic and pandemic influenza mRNAs respectively. Results presented at plots (c) and (f) of Figure 2 and Supplementary Figures 2-38 demonstrate that the most mutable positions are randomly distributed within each mRNA and do not form clusters. Absence of relationship between mutability value for every nucleotide position and corresponding value of moving average of individual standard deviations of the probabilities of nucleotides to be paired was confirmed by calculating Pearson correlation coefficients (Supplementary Table 4 and Supplementary Figures 39-58). All correlation coefficients were in a range from 0.006 to 0.222. This result refutes the intuitive notion that location of mutable positions would correspond to the least structured RNA regions, while sequence conserved positions would be collocated with the most structured RNA regions.

### Comparison of Mutation’s Effect on RNA PSRs

We generated two groups of *in silico* mutants by introducing synonymous mutations into influenza mRNAs. In the first group, mutations were introduced only into positions that are highly prone to being mutated; in the second one, mutations were introduced only into sequence conserved positions. The number of introduced *in silico* mutations was proportional to the length of every mRNA (Table 1). The effects of two groups of *in silico* mutations on structured RNA regions were compared, as described in the Materials and Methods section (Table 2). The results of statistical tests (Table 2) demonstrate that for majority of mRNAs the mutations introduced into sequence conserved positions have a greater effect on PSRs than mutations introduced into the mutable positions. This result stands out the most in mRNAs of non-pandemic NP, M2, and NS1 genes and pandemic NS2 gene.

**Table 2:** Comparative analysis of the effects on RNA PSRs elicited by *in silico* mutations in frequently vs. rarely mutating positions of H1N1 influenza mRNAs.

| Type | Gene name | Total length of PSRs | Percentage of mRNA covered by PSRs | Number of nucleotides within structured RNA regions, which changed their probability of being paired to a value outside of the naturally occurring range of probabilities for this position |  |  |  | P-value |
| --- | --- | --- | --- | --- | --- | --- | --- | --- |
|  |  |  |  | Mutations in positions that are prone to be mutated |  | Mutations in conserved positions |  |  |
|  |  |  |  | Mean | Standard deviation | Mean | Standard deviation |  |
| Non-pandemic | PB2 | 257 | 0.113 | 18.6 | 13.5 | 20.8 | 14.9 | 0.1395 |
|  | PB1 | 156 | 0.069 | 11.4 | 10.5 | 11.8 | 10.4 | 0.3264 |
|  | PA | 262 | 0.122 | 21.8 | 20.4 | 24.0 | 18.8 | 0.064 |
|  | HA | 85 | 0.050 | 5.9 | 5.4 | 5.3 | 5.6 | 0.0813 |
|  | NP | 300 | 0.200 | 24.6 | 17.2 | 31.3 | 17.7 | 0.001 |
|  | NA | 75 | 0.053 | 4.0 | 5.1 | 4.4 | 6.2 | 0.2283 |
|  | M1 | 334 | 0.440 | 24.3 | 23.3 | 21.1 | 21.2 | 0.1068 |
|  | M2 | 77 | 0.262 | 4.1 | 4.7 | 7.0 | 7.5 | 0.0022 |
|  | NS1 | 183 | 0.264 | 9.6 | 11.0 | 15.2 | 15.7 | 0.0008 |
|  | NS2 | 142 | 0.388 | 11.4 | 19.2 | 12.0 | 22.8 | 0.0311 |
| ic | PB2 | 232 | 0.102 | 16.7 | 14.0 | 16.0 | 13.0 | 0.2883 |
|  | PB1 | 251 | 0.110 | 15.8 | 14.3 | 17.0 | 14.6 | 0.1722 |
|  | PA | 138 | 0.064 | 12.0 | 10.7 | 12.5 | 10.8 | 0.3317 |
|  | HA | 205 | 0.121 | 17.5 | 18.0 | 15.9 | 18.5 | 0.0273 |
|  | NP | 351 | 0.234 | 41.2 | 27.5 | 44.8 | 27.8 | 0.0955 |
|  | NA | 84 | 0.060 | 4.7 | 6.8 | 5.5 | 7.7 | 0.3006 |
|  | M1 | 174 | 0.229 | 11.1 | 11.8 | 12.0 | 11.1 | 0.0917 |
|  | M2 | 77 | 0.262 | 10.5 | 8.7 | 9.8 | 11.2 | 0.0395 |
|  | NS1 | 31 | 0.047 | 2.8 | 5.7 | 3.3 | 5.7 | 0.1341 |
|  | NS2 | 146 | 0.399 | 10.4 | 14.7 | 21.5 | 28.0 | 0.0007 |

## DISCUSSION

Evolutionarily conserved RNA structural elements may perform important biological functions. Hence, identification and/or prediction of such elements can help in the understanding of the mechanism of RNA functions. This is true for identification of not only paired regions (stems), but loops too. In fact, kissing loop interactions are a common type of tertiary interaction motif in RNA that brings terminal loops together through Watson-Crick base pairing. Also, bulged nucleotides in the loop-loop interaction can be critical for ligand-dependent regulation. Yet, despite many efforts, it has still been a challenge to introduce an objective, quantitative, biologically meaningful and computationally friendly definition of what a “structured” RNA region is. Therefore, we had to propose a new definition and a new computational methodology free of these shortcomings.

In analyzing an individual RNA sequence, one has little chance to distinguish a biologically important structure formed by a folded molecule from simply a random shape with no biological importance. However, if one observes the same RNA configuration conserved and repeated across all related RNA sequences isolated from different strains and/or species, this increases the likelihood of biologically significant RNA structure. Following this logic, a definition of a structured RNA region should be based on a dataset of multiple aligned RNA sequences. Thus, assume that some structural element in a particular location is of such importance that it is present across all the strains. In this case, nucleotides in positions correspondent to the stem would have very high base pairing probabilities in all aligned RNA sequences of the dataset, while nucleotides in positions correspondent to the loop would have very low base pairing probabilities in all the strains. At the same time, nucleotides correspondent to a potential bistable structure would have their base pairing probabilities neither too high nor very low across all RNA sequences. Therefore, we propose that “structured RNA regions” are defined as the patterns of probability values of the nucleotides to be paired, which are manifested across the spectrum of strains and/or organisms. This definition equally considers the conservation of stems, loops and potential bistable structures while also providing a computationally friendly quantitative definition for the degree of RNA structure conservation.

Mathematically, the fact that nucleotides in a particular position in each RNA of a dataset are likely to belong to an evolutionarily conserved structural element means that if we collect values of pairing probabilities for this nucleotide from each RNA sequence in the dataset, and build a sample of these values to calculate its standard deviation, this standard deviation will be relatively low compared with the majority of standard deviations for other positions. Indeed, if this standard deviation is low, it means that mutations occurring in the analyzed RNA do not affect the base pairing probability of a nucleotide in this position across the spectrum of strains. Thus, it is most likely that mutations affecting pairing probability for this nucleotide are filtered out. This is a good indicator of evolutionary conservation and the biological importance of the RNA structure in this position. In contrary, if the standard deviation is high, it means that the correspondent nucleotide is very likely to be bonded in some strains but not in others; hence, the presence of any crucial RNA structure at this position is unlikely (unless there is a bistable secondary structure in this area playing roles in different functions). If an RNA contained five consecutive nucleotides with low standard deviations of their mean pairing probabilities, the region was considered structured.

### Applications of a Newly Introduced Computational Definition

Introduction of a new definition adequately describing the subject matter under study and development of a new technique for analysis, however, are only as good as they can be applied to a multitude of biological phenomena, generate new observations and experimentally testable hypotheses, explain old conundrums, and generate new questions (41). The presented approach was used to examine the existence of structured RNA regions in mRNAs of pandemic and non-pandemic influenza A H1N1 virus. This method revealed that influenza mRNAs contain nucleotide positions highly conserved in their base pairing probabilities. For every analyzed RNA type, such positions group together and constitute well-defined structured RNA regions, while the rest of the RNA molecule is significantly less structured. To the best of our knowledge, such mosaic structurization of RNA molecules was not reported previously. *In vitro* testing has confirmed that interfering with a stem of a previously unknown computationally predicted RNA structured region indeed reduces viral replication. We expect that future experimental testing will reveal the functions, these evolutionarily conserved RNA secondary structures, perform during the course of viral infection.

We hypothesized that mosaic structurization of influenza mRNAs may explain a long-standing conundrum of why different nucleotides in influenza genome mutate with such a varied frequency. The enormous influence of amino acid conservation could explain only a part of this phenomenon because many nucleotide substitutions are synonymous ones thus cannot be explained by amino acid conservation. The first hypothesis was that if a mutation happens within a structured RNA region it would disrupt the structure and be filtered out. Thus, even if a mutation rate was the same for all nucleotides, the only mutations observed in nature would be those happening outside of the structured RNA regions (PSR) and neutral for RNA structures. If an exact picture of each RNA structure was available, it would be possible to define structurally disruptive mutations visually as those that change the shape(s) of the structure(s). However, modern computational methods do not make it possible to predict exact RNA structures for long RNA molecules. Such predictions are inaccurate and cannot be relied upon (14, 42-44). Thus, we had to define structurally disruptive mutations based on the number of nucleotides in structured RNA regions, which would change their W-C pairing probabilities to a level aberrant of their naturally observed range. Contrary to original expectations, we showed that the nucleotide positions which are the least prone to being mutated do not collocate with regions of conserved RNA structures. Instead, the frequently and/or rarely mutating positions are randomly spread along the RNA sequences. Although it was demonstrated that the most frequently mutating positions within influenza genome are not collocated within unstructured RNA regions, this finding does not refute the main hypothesis that states: “Mutations, which occur in nucleotide positions that are the most prone to single nucleotide polymorphisms, have less of an effect on structured RNA regions than mutations, which occur in positions that are less likely to be changed”.

A mutation does not necessarily have to take place inside the PSR in order to be disruptive for a structure. For example, prior to mutation, a particular G was paired to a particular C forming a structure. If a mutation outside the structure changes some A to C, it may become a new paring partner for the G, thereby leading to an energetically more favorable RNA folding and disrupting the original structural element. This effect may be especially strong if mutations outside of the PSRs occur in combinations. Also, mutations in certain positions may have a greater effect on RNA structures than that of other positions. Thus, if RNA structures should indeed remain intact for successful viral propagation, all positions, SNPs in which would have a striking effect on the structures, would seem as rarely mutating compared to those positions, SNPs in which would have little effect on the structures. The results presented here support this hypothesis. We demonstrated for some influenza mRNAs that *in silico* mutations introduced into nucleotide positions, which mutate in the wild less frequently, would possess a greater disruptive effect on areas of conserved RNA structures than *in silico* mutations in positions which are known to mutate more frequently. As a result, mutations deleterious for vital RNA structures would be eliminated due to the negative selection pressure. This demonstrates that conservation of RNA structures could be a contributing mechanism defining a highly differential mutation rate for different influenza nucleotide positions. Additionally, the computational conclusion stipulates a direction for experimental testing. Although it is time/cost-consuming, it is possible to test RNA shapes experimentally (4, 45-49). If our hypothesis is correct, then influenza mRNAs observed in nature and those RNAs carrying mutations, which we predicted to be structurally non-disruptive, would possess similar RNA structures. By contrast, introducing into the RNA sequence mutations, which are predicted to disrupt structured RNA regions, would eliminate at least some of the RNA structures vital for a virus.

### Identification of Evolutionarily Conserved RNA Structural Elements

Plotting a graph with nucleotide positions on axis X and standard deviations of nucleotide paring probabilities for these positions on axis Y shows stretches along the RNA sequence with low standard deviations. These areas potentially have conserved RNA secondary structural elements. However, these graphs alone do not demonstrate whether the probability of a nucleotide to form bonds is high across different strains or low. In other words, a structurization profile may help identifying localization of RNA PSRs, but it does not indicate what kind of structure is there. Nevertheless, some assumptions about the RNA shape can be made if we complement structurization profiles with profiles presenting mean pairing probability for each nucleotide (i.e. for each nucleotide position in the RNA sequence, the pairing probability values from every RNA in the dataset would be used to calculate the mean for the position).

Extracting complex structures from comparing structurization profiles with profiles of mean pairing probabilities may require special analytical tools that are not a part of this first-stage study. However, discovering the simplest hairpin structure may not require additional instruments. Thus, when 10 nucleotides were found to possess very high means of probabilities to be bonded in the entire dataset, followed by 8 structurally conserved nucleotides which were apparently uncoupled, and then another 10 nucleotides that are likely to be paired and complementing the first 10 as W-C bonding partners, these findings showed existence of a previously unknown evolutionarily conserved RNA hairpin structure. *In vitro* testing has confirmed that interfering with a stem of a previously unknown computationally predicted RNA structured region indeed reduces viral replication. We expect that future experimental testing will reveal the functions, these evolutionarily conserved RNA secondary structures, perform during the course of viral infection.

It would be important to test whether pandemic and seasonal influenza strains indeed share some PSRs and whether the difference in RNA structurization may play a role in pandemic vs. non-pandemic viral phenotypes. Another direction of the future research would be to expand our computational definition of a structured RNA region to predict evolutionarily conserved RNA tertiary structures, especially in those RNAs that are hard to study by high-resolution experimental methods (50). In addition to helical segments, RNAs can fold into complex three-dimensional shapes. Computational modeling of RNA tertiary structures and determining of three-dimensional shapes of complex RNAs constitutes a major intellectual challenge (51-55). Thus, the most practical way to expand the proposed computational method to studying RNA 3D structures would be to incorporate RNA 3D structural modules that define sets of non-Watson-Crick base pairs embedded in WC pairs (56, 57).

### Novel Approach for the Rational Design of Live-Attenuated Vaccines and Anti-Viral Therapies

The method we proposed and applied to define structured RNA regions revealed several areas possessing conserved secondary structures in mRNAs of H1N1 influenza virus. As a next step, these structures have to be confirmed by *in vitro* analysis and their biological roles have to be assessed *in vitro* and/or *in vivo*. Potentially, structurally conserved RNA regions of viral RNAs may become a novel class of anti-viral drug targets. For example, anti-viral agents selectively disrupting RNA structures vital for a viral life cycle may become a new class of anti-viral therapies. As a preliminary proof of concept, we have demonstrated that an oligonucleotide binding the computationally predicted stem of a hairpin in a PSR, indeed acts as an anti-viral agent reducing *in vitro* viral replication. In contrast, statistically significant effect on viral replication was not observed if the infected cells transfected with the oligos of the same length, which bind outside of the predicted hairpin or do not bind to anything at all. Interestingly, even an oligonucleotide complementing the loop of this hairpin was unable to reduce viral replication in a statistically significant manner. Thus, the anti-viral effect was specific to disrupting the hairpin’s stem. RNA ISRAEU allows rapid rational design of oligonucleotide cocktails interfering with multiple computationally predicted structures, so no single or few mutations would result in a resistant viral strain.

Several approaches have been proposed for analysis of impact of SNPs on RNA structures and deleterious mutation prediction (1, 58), including RNAsnp (59, 60), SNPfold (61), RNAmute (62, 63), RNAmutants (64), and RDMAS (65). However, all these methods compare structures of the original and mutated RNAs assessing the distance, the effect on the RNA structure caused by SNPs. Although these methods are productive for the tasks they were developed for, they cannot be applied to our problem. We do not compare structures of an original and an altered RNA sequences. Instead, we compare structures of hundreds of RNA sequences without attributing any of them the “original” status. Therefore, the approach proposed here allows us to define: (i) a naturally occurring range of probabilities, which represents a range of probability values that are the most likely to be observed for natural RNA strains (see Quantitative Assessment of Mutation’s Effect on RNA PSRs in the Materials and Methods section for the specification) for every nucleotide position within an RNA region possessing an evolutionarily conserved structure; (ii) mutation(s) that would change base pairing probabilities within the structured RNA regions to an extent that the new probabilities would not belong to a naturally occurring range for corresponding positions.

Finally, we propose a new approach for the rational design of attenuated vaccines that would be based on predicting mutations disruptive for conserved RNA structures and introducing such mutations into viral genome. Indeed, disruption of an mRNA structure may serve as a functional gene knock out reducing expression of a viral gene to a level insufficient for viral cycle (12). Viral strain possessing such RNA can be administered to induce an immune response with little risk for a patient. Such attenuated viral strains can be grown on supporting cell lines actively expressing the limiting protein. Although LAVs are the most successful achievements in the history of public health (38), we believe there were no prior attempts to create LAVs based on perturbation of RNA structures.

## MATERIALS AND METHODS

### Data

Selecting H1N1 influenza mRNAs for this work constituted a crucial initial step. Influenza A genome consists of eight segments encoding seventeen proteins (66). Seven of those proteins were excluded from the analysis due to the limited information about them. It is known, however, that different influenza segments have different mutation rates (67). To eliminate potential bias that can be caused by disproportional representation of similar hemagglutinin (HA) and neuraminidase (NA) sequences (these two influenza genes are sequenced more often than the others because they constitute major viral antigens) and to compare evolutionary structure conservation between different influenza mRNAs, only completely sequenced influenza genomes were utilized in the analysis. An influenza genome was considered completely sequenced if it had no missing parts, no unknown nucleotides, and if sequences of the ten major mRNAs (namely, PB1, PB2, PA, HA, NP, NA, M1, M2, NS1, and NS2) were known. In order to further increase coherence of the dataset, only human influenza strains were utilized; other hosts were excluded because they demonstrate different characteristics (68). Finally, only those strains possessing the identical length of each influenza mRNA were selected. The fact that every mRNA of the same type has the same length in every viral genome selected eliminates potential mistakes, which could be introduced by effects of deletion and insertion polymorphisms (DIPs) on RNA secondary structures. Sequences of pandemic and non-pandemic complete influenza genomes satisfying the above mentioned criteria were downloaded from the Influenza Virus Resource (http://www.ncbi.nlm.nih.gov/genomes/FLU/FLU.html) (69).

### Filtering Redundant Sequences

Redundancy of data may introduce significant bias. To avoid it, one must only use a representative subset of sequences instead of analyzing all possible strains. Therefore, strains that were too similar were eliminated from further analysis; and, a non-redundant subset of strains was created. Any two strains in the non-redundant subset possess no less than 50 nucleotide differences per complete genome. In short, the first strain was chosen randomly from the dataset described in the previous section, then added to the non-redundant subset. Then, a different strain was randomly chosen and added to the non-redundant subset only if the newly chosen sequence had at least 50 nucleotide differences versus all strains in the non-redundant subset. This step was repeated until no more sequences could be added to the non-redundant subset. The described procedure was done separately for the pandemic and non-pandemic influenza datasets described above.

### Structural Conservation of a Nucleotide Position

As a first step, for each mRNA sequence in the datasets, the probability of every nucleotide within an RNA chain to be coupled via W-C bond was calculated. For that purpose, the RNAfold tool from the Vienna RNA package was used (70). RNAfold was used with the command line options −p that calculates the partition function and base pairing probability matrix, –noLP that disallows base pairs that can only occur as helices of length 1, and the default folding temperature fixed at 37°C. As a result, if a non-redundant dataset consisted of N sequences, a sample of N probability values would be created for each position (exactly N for a position, in which there is no deletion/insertion polymorphisms) within analyzed RNA. The standard deviation was calculated for every sample. The procedure described above was conducted for each of ten mRNAs from both subsets. Thus, we have calculated standard deviations of the probabilities of nucleotides to be paired for every nucleotide position within each of ten influenza mRNAs in both pandemic and non-pandemic datasets. To smooth stochastic fluctuations, moving averages of individual standard deviations with the sliding window of 5-nt length were calculated (Figure 2). To determine structure conserved positions, all moving average values of individual standard deviations from all mRNAs were combined to one dataset of moving averages, and the mean value and standard deviation of the values in that dataset were calculated. If an individual moving average calculated for a particular position was smaller than the overall mean of moving averages minus the overall standard deviation of moving averages, the correspondent position was considered “structure-conserved”.

### RNA Structurization and Structured RNA Regions

As described above, noticeable areas of structure-conserved positions possessing low standard deviation values were observed. Areas possessing at least five consequent structure-conserved nucleotides were defined as “structured RNA regions”. The described procedure was repeated separately for pandemic and non-pandemic influenza strains.

### Mutability

Intuitively, “mutability” demonstrates how likely it is for a nucleotide in a particular position to be mutated. Mathematically, this simple notion is defined as the value of Shannon entropy (35), which is calculated based on the frequencies of every ribonucleotide recorded in a particular position, with a pseudocount regularizers equal to 1 being added to the frequency of each of four ribonucleotides according to Laplace’s rule. To identify nucleotide positions that are the most/least prone to being mutated, the mutability value was computed for each nucleotide position. The more variable a set of ribonucleotides observed in a particular position, the higher the entropy. Then, all mutability values from all mRNAs were combined into one dataset. Those positions that had their mutability values higher than the 80th percentile of the dataset were considered as mutable positions. In contrast, those positions that did not contain SNPs among the sequences in the dataset were considered conserved positions. The described procedure was repeated separately for pandemic and non-pandemic influenza strains.

### Quantitative Assessment of Mutation’s Effect on RNA PSRs

Following the analysis discussed above, a new method was proposed and implemented, which defines “structurally disruptive mutations” based on their effect on structured RNA regions (PSRs). As described previously, two datasets of aligned influenza sequences were created. For each individual RNA sequence within the datasets, the probability of each nucleotide to be paired was computed. For every nucleotide position within coding regions of influenza mRNA sequences, the mean value and the standard deviation of the probabilities of nucleotides to be paired were calculated. Based on these values, a naturally occurring range of probabilities was calculated for every nucleotide position within a PSR. A naturally occurring range of probabilities was defined as a range of probabilities from the mean value decreased by two standard deviations to the mean value increased by two standard deviations.

Apparently, a mutation occurring in an RNA sequence may change probability of forming Watson-Crick pairs for multiple nucleotides within a particular sequence. For some of those nucleotides, their new probability values would still belong to the naturally occurring range of probabilities for this position. For other positions, the mutation would change their pairing probabilities to an extent that the new probabilities would not belong to their naturally occurring range. A quantitative effect of mutation(s) on RNA structurization is defined as a number of nucleotides within structured RNA regions (PSRs), which would change their probabilities to an extent that the new probabilities would not belong to a naturally occurring range for corresponding positions.

### Statistical Analysis: Do Mutations Taking Place in the Most vs. Least Often Mutating Positions Have a Different Effect on RNA Structurization?

Some positions in influenza genome are more prone to being mutated than others. The ability to define quantitatively effects of mutations on RNA structurization permitted the opportunity to propose a method for assessment, if mutations taking place in the frequently mutating positions have the same effect on RNA structurization as mutations occurring in the conserved ones. Two sets of *in silico* mutants were generated introducing synonymous mutations in nucleotide positions that are either the most or the least prone to being mutated. These two sets of mutations were compared for their effect on structured RNA regions.

In order to normalize for the length difference among influenza mRNAs, the number of changed nucleotides, which were introduced into each mRNA, was in proportion to the length of the mRNA (Table 1). The required number of synonymous SNPs was introduced into every mRNA sequence from the original datasets. In order to generate an *in silico* mutant from an original mRNA sequence, the required number of positions that are the most or the least prone to being mutated were randomly selected. Every codon, which contains the selected position, was changed to an alternative one encoding the same amino acid with the condition that the new codon is not observed in the particular position in any mRNA sequence from the datasets. Influenza mRNAs contain relatively high number of conserved positions and relatively few often mutating ones. As a result, for every mRNA, the number of *in silico* mutants with SNPs in conserved positions was equal to the number of wild type influenza strains in the datasets. However, due to a small number of frequently mutating positions, it was impossible for some mRNAs to generate the same number of unique mutants by introducing SNPs only to positions prone to being mutated. In those cases, all possible mutants were kept for further analysis - namely, 103 for non-pandemic M2, 95 for pandemic M2, and 109 for pandemic NS2.

For each computer-generated mutant, the probability of every nucleotide to be in a double-stranded conformation was calculated. Based on those probabilities, we calculated the number of nucleotides within structured RNA regions (PSRs), which changed their probability of being paired to a value outside of the naturally occurring range of probabilities for this position. Such numbers were combined into two datasets: one for mutations introduced into highly mutable positions and another – for mutations introduced in highly conserved positions. The Mann-Whitney U test was conducted for comparing these two datasets. The significance level for the test was Bonferroni-corrected by dividing the significance level of 5% by the total number of mRNAs in influenza virus, i.e. 10. The described procedure was repeated separately for pandemic and non-pandemic influenza strains.

### Profiles of Mean Pairing Probabilities

Profiles of mean pairing probabilities were created for influenza mRNAs (Figure 2 and Supplementary Figures 2-38). These profiles demonstrate how likely on average each nucleotide within an mRNA is to be paired based on an analysis of the entire dataset of sequences. As mentioned, for every RNA sequence in the dataset, the probability of every nucleotide within the RNA chain to be coupled via W-C bond was calculated. Then, for every nucleotide position, we computed the mean for probability values of this nucleotide based on all RNA sequences. The resulting series of means is used as a profile of mean pairing probabilities for a particular mRNA. The same work was performed for every influenza mRNA.

### Virus and Cells

Influenza virus A/California/7/09 (H1N1pdm) was provided by the Research Institute of Influenza museum of viruses, Saint-Petersburg, Russian Federation. The 50% tissue culture infective dose (TCID50) of this virus strain was defined by Reed–Muench method (72). The aliquots of virus were stored at −80°C. According to the American Tissue Madin-Darby canine kidney (MDCK) cell culture was provided from the cell collection of Research Institute of Influenza, Saint-Petersburg, Russian Federation. Cells were cultivated in cultural flasks using minimum essential medium Eagle alpha modification (aMEM, Biolot) with 2mM L-glutamine supplemented with 10% heat-inactivated fetal bovine serum (FBS, GIBCO, USA).

### Design of Antisense DNA-oligonucleotides

We designed antisense oligonucleotides, which may potentially disrupt the aforementioned predicted RNA-structure. A random oligonucleotide, “rand10”, with minimal probability of having targets in the human hosts and viral genome was used as a control. Another control, “off10”, was an oligonucleotide with a target to the adjacent region on the NS2 gene mRNA (Table 3).

**Table 3:**
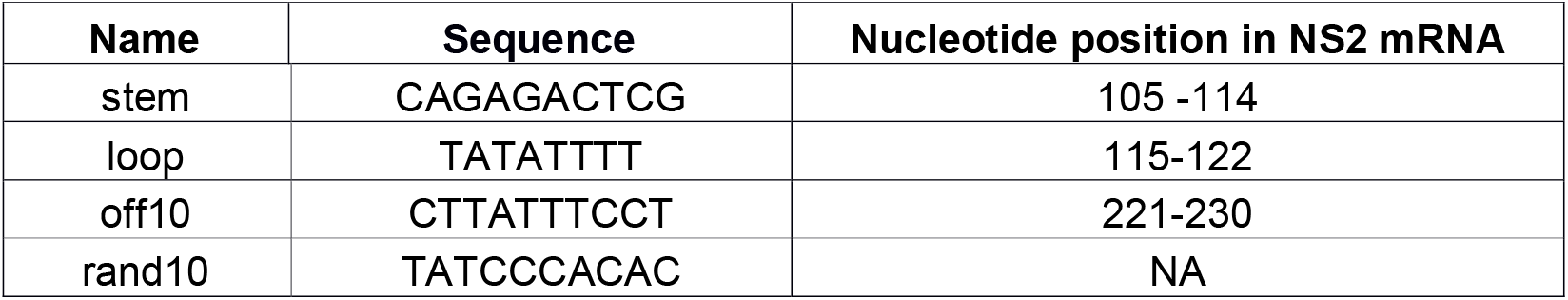
List of antisense oligonucleotides used to examine influenza viral replication inhibition.

### Cell Viability Assay

The cell viability was determined 24 hours post infection and transfection by microtetrazolium test (MTT assay). A solution of MTT [3-(4,5-dimethylthiazol-2-yl)-2,5-diphenyltetrazolium bromide] (Sigma) at a concentration of 2,5 mg/ml was prepared in PBS. The medium was removed, the cells were washed once with PBS, MTT solution was added into the wells (100 ul/well). The cells were incubated at 37°C and 5% CO 2 for 4 hours and then the solution was removed and 96% ethanol (100 ul/well) was added for formazan crystals dissolving. The absorbance signal was measured using multifunctional reader CLARIOstar ^®^(BMG LABTECH, Germany) at 535 nm.

### Virus Infection and Transfection

The cells were detached by 0,25% trypsin/EDTA solution (Invitrogen) for 5 min and plated in 96-well plates (Nucl) at 104 cells per well the day before the infection experiment. Cells were washed twice with Dulbecco’s Phosphate-Buffered Saline (DPBS, GIBCO) and infected with A/California/7/09 (H1N1pdm) viral strain in 100TCID50 dose per well. The medium for cells infection was minimum essential medium Eagle alpha modification (aMEM, Biolot) with 2mM L-glutamine, 2,5 ug/ml trypsin TPCK-treated from bovine pancreas (TPCK, Sigma) and 1:100 antibiotic-antimycotic (100X, GIBCO). Inoculation was conducted at 37°C and 5% CO2 for 60 minutes. Then the medium was removed and the cells were transfected using 100 μl of OptiPro SFM medium (GIBCO) contained 10 μM of DNA-oligonucleotides and 0.7 μl/well of Lipofectamine 2000 (Invitrogen) according to the manufacturer’s protocol. In addition, the transfection medium is also supplemented with 1:100 antibiotic-antimycotic (100X) and 2,5 ug/ml TPCK. Viral control samples were also transfected with lipofectamin 2000 only, without any oligonucleotides. Four hours post-transfection, the medium was replaced with fresh aMEM (Biolot) which contained 2mM L-glutamine, 2,5 ug/ml TPCK and 1:100 antibiotic-antimycotic (100X). Twenty four hours post-infection, cells were used for the further relative ELISA analyses. Each treatment was performed in triplicates.

### Enzyme-Linked Immunosorbent Assays (ELISA)

Twenty four hours post influenza virus infection and transfection with oligonucleotides, continuity of the cell monolayer was assessed microscopically. Then, medium was removed and the MDCK cells in 96-wells Nunc plates were fixed with 150 μl per well of cold 80% acetone at 4°C for 30 minutes. The fixed samples were washed three times with phosphate buffered saline containing 0.05% Tween (PBS-T) and blocked with 5% milk dissolved in PBST (200 ul/well) for 30 minutes at 37°C. The fixed cells were incubated with 1ug/ml mouse antibody against NP-protein (100 ul/well) produced in the Influenza Research Institute (clone 4H1) at 37°C for 1 hour. After the next three washes the secondary goat anti-mouse antibody conjugated with horseradish peroxidase (GAM-HRP, BioRad, USA) was added at 1ug/ml (100 ul/well) and incubated for 1 hour at 37°C. Cells were washed three times with PBS-T followed by adding TMB Peroxidase EIA Substrate Kit (Bio-Rad, USA) according to manufacturer’s instructions for further absorbance analysis. The absorbance was measured using multifunctional reader CLARIOstar ^®^ (BMG LABTECH) as delta optical density OD 450 – OD 655. The absorbance signal from uninfected cells was taken as zero and was subtracted from the obtained values of the samples. The results were presented relative to infection control.

### Statistical Analysis of Viral Replication Inhibition Assay

Data shown are means +/- SD as percentage of untreated “Flu” group. P-values for comparing the four treatment groups with the untreated group (Flu) were calculated using student T test. The significance level for the test was Bonferroni-corrected by dividing the significance level of 0.05 by the total number of group comparisons, i.e. 10. Analysis was performed using the R software.

## Supporting information

## ACKNOWLEDGEMENTS

The authors would like to thank Prof. Dmitrij Frishman, a Ph.D. advisor of Andrey Chursov. This work was partially performed during AC term in Dmitrij Frishman’s lab. The authors would also like to thank Prof. Adolfo Garcia-Sastre for illuminating comments on the article and the methodology, and Prof. Andrey Mironov and Prof. Ilya Muchnik for intellectually stimulating discussions.

