## Supplementary material for "Novel Computational Method to Define RNA PSRs Explains Influenza A Virus Nucleotide Conservation"

**Supplementary Table 1**: Length of coding region and numbers of highly mutable and conserved positions identified in every influenza mRNA.

| **Gene name** | **Non-pandemic** | | | | | | **Pandemic** | | | | | |
|---|---|---|---|---|---|---|---|---|---|---|---|---|
| | **Coding region length** | **Non-mutating positions** | **Highly mutable positions** | | **Mutable third codon positions** | | **Coding region length** | **Non-mutating positions** | **Highly mutable positions** | | **Mutable third codon positions** | |
| | | | **Number** | **Percentage of total length, %** | **Number** | **Percentage of highly mutable, %** | | | **Number** | **Percentage of total length, %** | **Number** | **Percentage of highly mutable, %** |
| PB2 | 2280 | 1763 | 262 | 11.5 | 211 | 80.5 | 2280 | 1717 | 289 | 12.7 | 232 | 80.3 |
| PB1 | 2274 | 1762 | 277 | 12.2 | 237 | 85.6 | 2274 | 1679 | 286 | 12.6 | 226 | 79.0 |
| PA | 2151 | 1682 | 244 | 11.3 | 207 | 84.8 | 2151 | 1583 | 266 | 12.4 | 209 | 78.6 |
| HA | 1698 | 1254 | 241 | 14.2 | 173 | 71.8 | 1701 | 1164 | 269 | 15.8 | 179 | 66.5 |
| NP | 1497 | 1187 | 164 | 11.0 | 145 | 88.4 | 1497 | 1180 | 150 | 10.0 | 125 | 83.3 |
| NA | 1413 | 1008 | 219 | 15.5 | 154 | 70.3 | 1410 | 1000 | 201 | 14.3 | 128 | 63.7 |
| M1 | 759 | 635 | 60 | 7.9 | 51 | 85.0 | 759 | 606 | 67 | 8.8 | 57 | 85.1 |
| M2 | 294 | 232 | 28 | 9.5 | 18 | 64.3 | 294 | 230 | 20 | 6.8 | 10 | 50.0 |
| NS1 | 693 | 515 | 82 | 11.8 | 46 | 56.1 | 660 | 467 | 82 | 12.4 | 44 | 53.7 |
| NS2 | 366 | 296 | 38 | 10.4 | 28 | 73.7 | 366 | 297 | 27 | 7.4 | 18 | 66.7 |

**Supplementary Table 2:** Non-pandemic and pandemic H1N1 influenza A strains used for the analysis.

| # | Non-pandemic | Pandemic |
|---|---|---|
| 1 | A/England/192/2000 | A/Minneapolis/INS3_675/2012 |
| 2 | A/California/VRDL209/2009 | A/Helsinki/720M/2014 |
| 3 | A/Alabama/UR06-0455/2007 | A/Aalborg/INS132/2009 |
| 4 | A/Arkansas/WRAIR1249P/2009 | A/Aarhus/INS254/2009 |
| 5 | A/Auckland/579/2000 | A/Aarhus/INS3_653/2011 |
| 6 | A/Auckland/580/2000 | A/Aarhus/INS3_654/2011 |
| 7 | A/Auckland/605/2001 | A/Aarhus/INS609/2011 |
| 8 | A/Auckland/619/2005 | A/Aarhus/INS612/2011 |
| 9 | A/Boston/1/2009 | A/Alabama/01/2010 |
| 10 | A/Boston/10/2009 | A/Alaska/38/2009 |
| 11 | A/Boston/12/2007 | A/Arizona/20/2009 |
| 12 | A/Boston/23/2009 | A/Athens/INS163/2009 |
| 13 | A/Boston/34/2008 | A/Athens/INS271/2009 |
| 14 | A/Boston/46/2009 | A/Athens/INS345/2009 |
| 15 | A/Boston/49/2008 | A/Athens/INS398/2010 |
| 16 | A/Boston/6/2009 | A/Athens/INS412/2010 |
| 17 | A/Boston/67/2009 | A/Athens/INS416/2010 |
| 18 | A/Boston/93/2009 | A/Athens/INS571/2011 |
| 19 | A/California/UR06-0125/2007 | A/Bangkok/INS3_681/2012 |
| 20 | A/California/UR06-0393/2007 | A/Bangkok/INS424/2010 |
| 21 | A/California/VRDL134/2009 | A/Bangkok/INS425/2010 |
| 22 | A/California/VRDL151/2009 | A/Bangkok/INS478/2010 |
| 23 | A/California/VRDL152/2009 | A/Bangkok/INS481/2010 |
| 24 | A/California/VRDL175/2009 | A/Bangkok/INS490/2010 |
| 25 | A/California/VRDL191/2009 | A/Bangkok/INS505/2010 |
| 26 | A/California/VRDL193/2009 | A/Bangkok/INS511/2010 |
| 27 | A/California/VRDL252/2009 | A/Bogota/WRAIR0435N/2009 |

| | | |
|---|---|---|
| 28 | A/California/VRDL256/2009 | A/Boston/685/2009 |
| 29 | A/Canada/591/2004 | A/Boston/DOA08/2011 |
| 30 | A/Canterbury/01/2001 | A/Boston/DOA14/2011 |
| 31 | A/Canterbury/126/2001 | A/Boston/DOA2-099/2012 |
| 32 | A/Canterbury/27/2000 | A/Boston/DOA40/2011 |
| 33 | A/Chile/8885/2001 | A/Boston/DOA90/2012 |
| 34 | A/Christchurch/1/2003 | A/Boston/YGA_00037/2013 |
| 35 | A/Colorado/UR06-0053/2007 | A/Boston/YGA_01185/2013 |
| 36 | A/DaNang/DN238/2008 | A/Boston/YGA_01217/2013 |
| 37 | A/DaNang/DN345/2008 | A/Brooklyn/INS549/2011 |
| 38 | A/DaNang/DN431/2008 | A/California/VRDL107/2009 |
| 39 | A/England/493/2006 | A/California/VRDL4/2010 |
| 40 | A/England/494/2006 | A/California/VRDL81/2009 |
| 41 | A/England/545/2007 | A/California/WR1316P/2009 |
| 42 | A/England/593/2006 | A/Cambridge/INS528/2010 |
| 43 | A/Florida/UR06-0208/2007 | A/Changchun/01/2009 |
| 44 | A/Florida/UR07-0022/2008 | A/Chicago/YGA_04019/2012 |
| 45 | A/HaNoi/Q421/2006 | A/Chile/115/2010 |
| 46 | A/HaNoi/TX200/2008 | A/Chile/6/2010 |
| 47 | A/Hanoi/ISBM31/2005 | A/Darlinghurst/INS3_643/2011 |
| 48 | A/Hong Kong/1870/2008 | A/District of Columbia/INS525/2010 |
| 49 | A/Hue/H386/2008 | A/District of Columbia/INS600/2011 |
| 50 | A/Illinois/UR06-0146/2007 | A/District of Columbia/WRAIR0313/2011 |
| 51 | A/Johannesburg/159/1997 | A/England/05120538/2010 |
| 52 | A/Kentucky/UR06-0057/2007 | A/England/280/2010 |
| 53 | A/Kentucky/UR06-0539/2007 | A/England/859/2009 |
| 54 | A/Kentucky/UR07-0061/2008 | A/Finland/102/2014 |
| 55 | A/Kyoto/08K056/2009 | A/Finland/1520N/2011 |
| 56 | A/Malaysia/11641/1997 | A/Finland/30/2014 |

| | | |
|---|---|---|
| 57 | A/Malaysia/14075/2000 | A/Finland/61/2014 |
| 58 | A/Malaysia/15042/1998 | A/Gainesville/05/2014 |
| 59 | A/Malaysia/1686034/2006 | A/Georgia/T51700/2012 |
| 60 | A/Malaysia/1706215/2007 | A/Guangzhou/GIRD74/2010 |
| 61 | A/Malaysia/1798564/2007 | A/Gunma/267/2009 |
| 62 | A/Malaysia/2143035/2009 | A/Gunma/287/2009 |
| 63 | A/Malaysia/30025/2004 | A/Hamburg/INS92/2009 |
| 64 | A/Malaysia/32110/2005 | A/Helsinki/100/2013 |
| 65 | A/Malaysia/33132/2005 | A/Helsinki/1127/2014 |
| 66 | A/Malaysia/34450/2006 | A/Helsinki/11IH2213/2011 |
| 67 | A/Malaysia/35164/2006 | A/Helsinki/1289/2013 |
| 68 | A/Managua/1038.01/2008 | A/Helsinki/147/2013 |
| 69 | A/Managua/107.01/2008 | A/Helsinki/220M/2014 |
| 70 | A/Managua/2055.01/2008 | A/Helsinki/2430/2012 |
| 71 | A/Managua/254.01/2008 | A/Helsinki/39/2013 |
| 72 | A/Managua/3153.01/2008 | A/Helsinki/473N/2014 |
| 73 | A/Managua/3759.02/2008 | A/Helsinki/490/2013 |
| 74 | A/Memphis/1/2001 | A/Helsinki/771M/2014 |
| 75 | A/Memphis/15/2000 | A/Houston/JMM_131/2013 |
| 76 | A/Memphis/6/2001 | A/Hubei/75/2009 |
| 77 | A/Memphis/7/2001 | A/Hubei/76/2009 |
| 78 | A/Mississippi/UR06-0378/2007 | A/India/Nag132467/2013 |
| 79 | A/Nagasaki/07N035/2008 | A/India/Nsk12388/2012 |
| 80 | A/Nanchang/11/1996 | A/India/P1112874/2011 |
| 81 | A/Nanchang/16A/1999 | A/India/P1114854/2011 |
| 82 | A/Nanchang/8/1996 | A/India/P121773/2012 |
| 83 | A/New Caledonia/20-JY2/1999 | A/India/P121778/2012 |
| 84 | A/New South Wales/26/2000 | A/India/P12946/2012 |
| 85 | A/New York/08-1253/2008 | A/India/P131027/2013 |

| | | |
|---|---|---|
| 86 | A/New York/1062/2007 | A/India/P131845/2013 |
| 87 | A/New York/1104/2008 | A/India/P132194/2013 |
| 88 | A/New York/1159/2009 | A/Iowa/04/2010 |
| 89 | A/New York/1692/2009 | A/Jiangsu/1/2009 |
| 90 | A/New York/205/2001 | A/Kazan/CRIE-02/2013 |
| 91 | A/New York/281/2001 | A/Khon Kaen/INS3_649/2012 |
| 92 | A/New York/306/2001 | A/Liaoning/1/2009 |
| 93 | A/New York/442/2001 | A/Lima/INS3_671/2012 |
| 94 | A/New York/494/2002 | A/Lima/WRAIR0672F/2009 |
| 95 | A/New York/UR06-0199/2007 | A/Managua/0305_10/2010 |
| 96 | A/Shanghai/2/1997 | A/Managua/1244.01/2009 |
| 97 | A/Singapore/14/2001 | A/Managua/3246.01/2010 |
| 98 | A/South Australia/58/2005 | A/Melbourne/INS471/2010 |
| 99 | A/St. Petersburg/8/2006 | A/Mexico/InDRECTRLA/2010 |
| 100 | A/Taiwan/123/2002 | A/Moscow/WRAIR1627T/2009 |
| 101 | A/Taiwan/5072/1999 | A/Nepal/VIROAF5/2012 |
| 102 | A/Tennessee/UR06-0080/2007 | A/New York/2372/2010 |
| 103 | A/Texas/UR06-0026/2007 | A/New York/3230/2010 |
| 104 | A/Thailand/CU-B589/2009 | A/New York/6530/2010 |
| 105 | A/Thailand/CU-H223/2009 | A/New York/7480/2010 |
| 106 | A/Waikato/11/2005 | A/New York/INS150/2009 |
| 107 | A/Western Australia/18/2001 | A/New York/WC-LVD-13-001/2013 |
| 108 | | A/New York/WC-LVD-13-007/2013 |
| 109 | | A/New York/WC-LVD-13-028/2013 |
| 110 | | A/New York/WC-LVD-14-001/2014 |
| 111 | | A/New York/WC-LVD-14-027/2014 |
| 112 | | A/New York/WC-LVD-14-034/2014 |
| 113 | | A/New York/WC-LVD-14-044/2014 |
| 114 | | A/New York/WC-LVD-14-058/2014 |

| | | |
|---|---|---|
| 115 | | A/Nicaragua/4136_07/2013 |
| 116 | | A/Nicaragua/4211_07/2013 |
| 117 | | A/Nicaragua/6123_01/2011 |
| 118 | | A/Nicaragua/6263_05/2013 |
| 119 | | A/Nicaragua/AGA2-52/2011 |
| 120 | | A/North Carolina/05/2010 |
| 121 | | A/North Fitzroy/INS464/2010 |
| 122 | | A/Northern Ireland/04380108/2010 |
| 123 | | A/Novosibirsk/KSH/2011 |
| 124 | | A/Odense/INS143/2009 |
| 125 | | A/Ontario/720545/2010 |
| 126 | | A/Quito/WRAIR0617N/2009 |
| 127 | | A/San Diego/INS101/2009 |
| 128 | | A/San Diego/INS105/2009 |
| 129 | | A/Santiago/p13d2/2013 |
| 130 | | A/Santiago/p15d1/2011 |
| 131 | | A/Santiago/p18d0/2013 |
| 132 | | A/Santiago/p9d1/2011 |
| 133 | | A/Singapore/EN193/2010 |
| 134 | | A/Singapore/GP10/2010 |
| 135 | | A/Singapore/GP3839/2009 |
| 136 | | A/Singapore/GP413/2010 |
| 137 | | A/Singapore/GP4406/2010 |
| 138 | | A/Singapore/GP489/2010 |
| 139 | | A/Singapore/GP496/2011 |
| 140 | | A/Singapore/GP868/2011 |
| 141 | | A/Singapore/GP908/2011 |

| | | |
|---|---|---|
| 142 | | A/Singapore/ON2136/2009 |
| 143 | | A/Singapore/SS34/2010 |
| 144 | | A/Singapore/TT157/2011 |
| 145 | | A/Sydney/DD3-10/2010 |
| 146 | | A/Sydney/DD3-11/2010 |
| 147 | | A/Sydney/DD3-27/2010 |
| 148 | | A/Sydney/DD3-37/2010 |
| 149 | | A/Tallinn/INS182/2010 |
| 150 | | A/Tennessee/F1093A/2010 |
| 151 | | A/Texas/JMS393/2009 |
| 152 | | A/Thailand/CU-B2357/2010 |
| 153 | | A/Thailand/CU-H2176/2010 |
| 154 | | A/Thailand/H1255/2010 |
| 155 | | A/Uganda/MUWRP-059/2009 |
| 156 | | A/Uganda/MUWRP-102/2009 |
| 157 | | A/Uganda/MUWRP-176/2010 |
| 158 | | A/Uganda/MUWRP-212/2010 |
| 159 | | A/Uganda/MUWRP-224/2010 |
| 160 | | A/Uganda/MUWRP-240/2011 |
| 161 | | A/Utah/02/2010 |
| 162 | | A/Viet Nam/11032010/2009 |
| 163 | | A/Viet Nam/15032004/2009 |
| 164 | | A/Warsaw/INS312/2009 |
| 165 | | A/Warsaw/INS3_657/2011 |
| 166 | | A/Wisconsin/629-D00134/2009 |
| 167 | | A/Wisconsin/629-D02424/2009 |
| 168 | | A/Wurzburg/INS382/2009 |

| | | |
|---|---|---|
| 169 | | A/Yaroslavl/IIV-196/2009 |
| 170 | | A/Zhejiang/3/2009 |
| 171 | | A/Zhejiang/88/2009 |
| 172 | | A/Zhejiang/X1/2009 |
| 173 | | A/Zhejiang/X2/2009 |

**Supplementary Table 3**: Coordinates of identified structured RNA regions.

| Gene Name | Non-pandemic | | Pandemic | |
|---|---|---|---|---|
| | Start Position | End Position | Start Position | End Position |
| M1 | 8 | 12 | 59 | 63 |
| | 26 | 30 | 113 | 117 |
| | 50 | 55 | 138 | 156 |
| | 57 | 63 | 185 | 214 |
| | 89 | 93 | 247 | 254 |
| | 112 | 142 | 284 | 293 |
| | 149 | 189 | 361 | 366 |
| | 208 | 217 | 372 | 379 |
| | 232 | 288 | 449 | 454 |
| | 320 | 337 | 512 | 518 |
| | 340 | 344 | 520 | 531 |
| | 350 | 367 | 535 | 540 |
| | 371 | 393 | 610 | 628 |
| | 397 | 410 | 678 | 699 |
| | 492 | 503 | 707 | 717 |
| | 512 | 520 | | |
| | 539 | 547 | | |
| | 610 | 614 | | |
| | 620 | 628 | | |
| | 646 | 660 | | |
| | 667 | 676 | | |
| | 678 | 697 | | |
| M2 | 2 | 8 | 59 | 72 |
| | 17 | 50 | 148 | 153 |
| | 187 | 200 | 184 | 192 |

| | | | | |
|---|---|---|---|---|
| | 237 | 251 | 223 | 227 |
| | 258 | 264 | 236 | 278 |
| NS1 | 198 | 220 | 537 | 541 |
| | 270 | 298 | 577 | 602 |
| | 337 | 343 | | |
| | 377 | 412 | | |
| | 439 | 443 | | |
| | 469 | 483 | | |
| | 505 | 550 | | |
| | 563 | 584 | | |
| NS2 | 9 | 129 | 7 | 30 |
| | 208 | 221 | 33 | 38 |
| | 347 | 353 | 40 | 87 |
| | | | 104 | 136 |
| | | | 210 | 214 |
| | | | 311 | 318 |
| | | | 321 | 334 |
| | | | 343 | 350 |
| PA | 262 | 273 | 99 | 125 |
| | 278 | 286 | 147 | 152 |
| | 307 | 317 | 312 | 318 |
| | 513 | 517 | 407 | 415 |
| | 855 | 868 | 914 | 930 |
| | 874 | 885 | 953 | 959 |
| | 1321 | 1326 | 967 | 976 |
| | 1332 | 1338 | 989 | 1002 |
| | 1384 | 1388 | 1004 | 1011 |
| | 1471 | 1480 | 1113 | 1117 |

| | | | | |
|---|---|---|---|---|
| | 1551 | 1556 | 1549 | 1556 |
| | 1575 | 1579 | 1832 | 1842 |
| | 1682 | 1699 | 2047 | 2055 |
| | 1728 | 1764 | | |
| | 1797 | 1801 | | |
| | 1818 | 1822 | | |
| | 1824 | 1852 | | |
| | 1892 | 1896 | | |
| | 1944 | 1950 | | |
| | 2061 | 2114 | | |
| NP | 3 | 7 | 100 | 166 |
| | 72 | 83 | 183 | 187 |
| | 187 | 210 | 200 | 213 |
| | 224 | 230 | 225 | 234 |
| | 244 | 248 | 241 | 247 |
| | 262 | 267 | 264 | 276 |
| | 284 | 288 | 323 | 344 |
| | 290 | 297 | 369 | 374 |
| | 325 | 329 | 384 | 389 |
| | 369 | 377 | 434 | 439 |
| | 419 | 423 | 504 | 508 |
| | 452 | 463 | 512 | 520 |
| | 637 | 656 | 539 | 543 |
| | 662 | 666 | 588 | 592 |
| | 680 | 688 | 672 | 678 |
| | 740 | 745 | 736 | 743 |
| | 751 | 760 | 769 | 777 |
| | 762 | 767 | 808 | 815 |
| | 1034 | 1038 | 824 | 832 |

| | | | | |
|---|---|---|---|---|
| | 1053 | 1069 | 1048 | 1057 |
| | 1072 | 1087 | 1071 | 1076 |
| | 1090 | 1101 | 1183 | 1187 |
| | 1127 | 1139 | 1218 | 1222 |
| | 1189 | 1202 | 1310 | 1314 |
| | 1329 | 1344 | 1324 | 1340 |
| | 1423 | 1438 | 1368 | 1372 |
| | 1443 | 1451 | 1384 | 1392 |
| | 1460 | 1482 | 1396 | 1400 |
| | | | 1408 | 1440 |
| | | | 1444 | 1450 |
| | | | 1461 | 1483 |
| PB1 | 45 | 51 | 67 | 169 |
| | 314 | 332 | 552 | 563 |
| | 338 | 349 | 579 | 598 |
| | 351 | 356 | 707 | 713 |
| | 412 | 416 | 812 | 825 |
| | 550 | 561 | 854 | 858 |
| | 606 | 611 | 1067 | 1071 |
| | 619 | 632 | 1229 | 1241 |
| | 1066 | 1071 | 1288 | 1296 |
| | 1151 | 1159 | 1326 | 1330 |
| | 1386 | 1417 | 1482 | 1489 |
| | 1429 | 1433 | 1498 | 1519 |
| | 1592 | 1598 | 1861 | 1865 |
| | 1947 | 1951 | 2020 | 2035 |
| | 1956 | 1966 | 2048 | 2054 |
| PB2 | 6 | 12 | 8 | 14 |
| | 38 | 55 | 200 | 233 |

| | | | | |
|---|---|---|---|---|
| | 95 | 99 | 464 | 476 |
| | 121 | 126 | 501 | 505 |
| | 281 | 286 | 788 | 794 |
| | 361 | 365 | 962 | 967 |
| | 372 | 378 | 1102 | 1124 |
| | 384 | 390 | 1450 | 1454 |
| | 417 | 422 | 1456 | 1465 |
| | 594 | 599 | 1498 | 1521 |
| | 779 | 789 | 1534 | 1555 |
| | 843 | 873 | 1618 | 1622 |
| | 909 | 922 | 1666 | 1676 |
| | 929 | 949 | 1682 | 1688 |
| | 978 | 986 | 1694 | 1708 |
| | 988 | 993 | 1844 | 1848 |
| | 998 | 1006 | 1938 | 1964 |
| | 1233 | 1239 | 2048 | 2053 |
| | 1443 | 1473 | | |
| | 1844 | 1848 | | |
| | 1973 | 1977 | | |
| | 2014 | 2040 | | |
| | 2113 | 2120 | | |
| HA | 349 | 353 | 135 | 140 |
| | 367 | 371 | 224 | 228 |
| | 559 | 564 | 470 | 482 |
| | 645 | 653 | 703 | 707 |
| | 1048 | 1056 | 767 | 772 |
| | 1065 | 1074 | 898 | 905 |
| | 1120 | 1124 | 943 | 954 |
| | 1141 | 1147 | 966 | 981 |

| | | | | |
|---|---|---|---|---|
| | 1200 | 1208 | 1090 | 1094 |
| | 1266 | 1275 | 1140 | 1144 |
| | 1601 | 1610 | 1174 | 1185 |
| | | | 1217 | 1222 |
| | | | 1224 | 1243 |
| | | | 1255 | 1278 |
| | | | 1280 | 1288 |
| | | | 1291 | 1300 |
| | | | 1303 | 1307 |
| | | | 1313 | 1320 |
| | | | 1328 | 1346 |
| | | | 1570 | 1574 |
| | | | 1661 | 1666 |
| NA | 63 | 77 | 13 | 23 |
| | 83 | 92 | 564 | 585 |
| | 97 | 102 | 590 | 596 |
| | 550 | 561 | 1047 | 1056 |
| | 564 | 568 | 1064 | 1073 |
| | 643 | 660 | 1268 | 1291 |
| | 1226 | 1234 | | |

**Supplementary Table 4:** Pearson correlation coefficients between the mutability value (i.e. Shannon entropy) for every nucleotide position and corresponding value of moving average of individual standard deviations of the base-pairing probabilities of nucleotides. The p-value column shows the measure of statistical significance of the corresponding correlation coefficient.

| | **Gene Name** | **Correlation Coefficient** | **P-value** |
|---|---|---|---|
| **Non pandemic** | PB2 | 0.120 | 8.86E-009 |
| | PB1 | 0.102 | 1.20E-006 |
| | PA | 0.106 | 7.85E-007 |
| | HA | 0.103 | 2.04E-005 |
| | NP | 0.118 | 5.16E-006 |
| | NA | 0.067 | 0.0118 |
| | M1 | 0.081 | 0.0265 |
| | M2 | 0.140 | 0.0171 |
| | NS1 | 0.110 | 3.97E-003 |
| | NS2 | 0.129 | 0.0143 |
| **Pandemic** | PB2 | 0.123 | 3.63E-009 |
| | PB1 | 0.095 | 5.91E-006 |
| | PA | 0.059 | 6.35E-003 |
| | HA | 0.098 | 5.73E-005 |
| | NP | 0.149 | 7.76E-009 |
| | NA | 0.090 | 7.44E-004 |
| | M1 | 0.074 | 0.0423 |
| | M2 | 0.006 | 0.9240 |
| | NS1 | 0.155 | 7.01E-005 |
| | NS2 | 0.222 | 2.02E-005 |

**Supplementary Figure 1:** Number of non-pandemic and pandemic influenza strains left after filtering very similar sequences depending on the identity threshold.

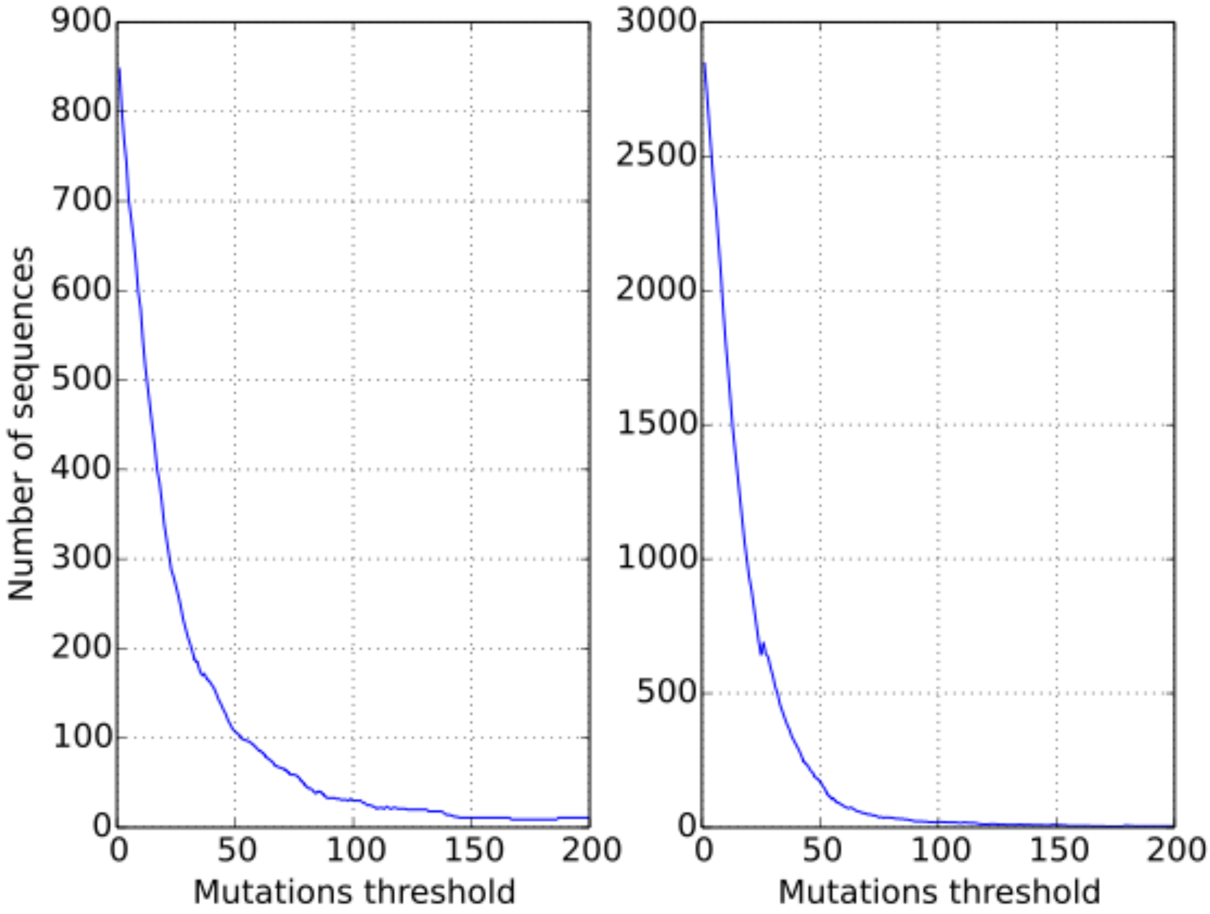

**Supplementary Figures 2-38:** Structure variability and mutability profiles for non-pandemic (a, b, and c) and pandemic (d, e, and f) influenza mRNAs. Plots a and d demonstrate structure conservation profiles; namely, they show the moving average that was calculated by applying a sliding window approach to smooth individual fluctuations of standard deviations of nucleotide base pairing probabilities. The blue solid line demonstrates the mean level of all moving average values, and the blue dashed line demonstrates the level equal to the mean of all moving average values decreased by the standard deviation of all moving average values. In this case, the mean and the standard deviation were computed based on all moving average values from all mRNAs of a particular type (pandemic or non-pandemic) of influenza strains. According to our definition, when the moving average goes below the blue dashed line, it is a structured RNA region. Such regions are colored with green across the plots. Plots b and e demonstrate profiles of the mean values of probabilities of nucleotide positions to be in a double-stranded conformation. If this value is close to 1, it means that in most strains in the dataset the correspondent nucleotide has a very high probability to be paired; and, if this value is close to 0, the correspondent nucleotide is very likely to be unpaired in most strains in the dataset. Plots c and f demonstrate mutability profiles for influenza mRNAs. Mutability of every nucleotide position is computed as a value of Shannon entropy which is calculated based on frequency of every ribonucleotide in a particular position.

**Supplementary Figure 2**

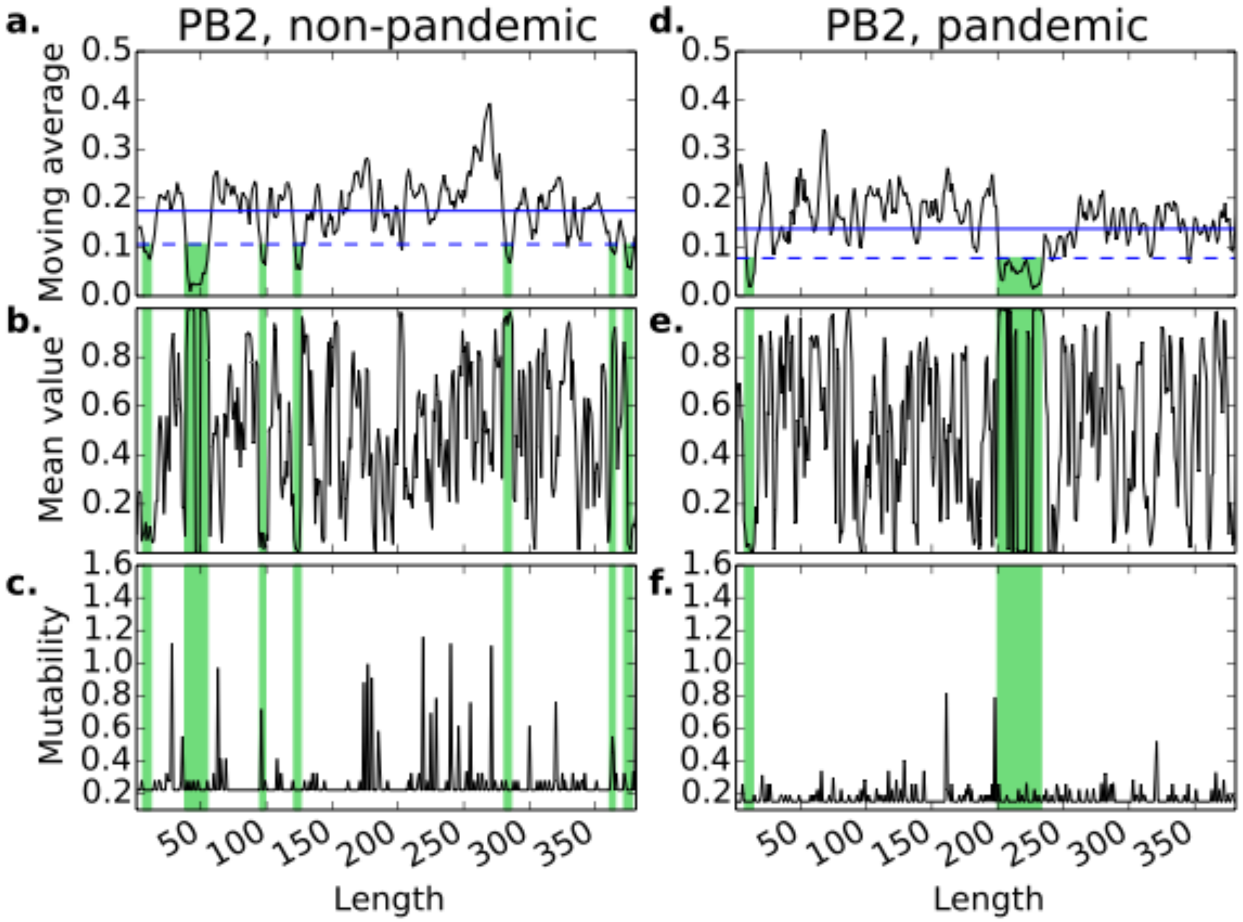

**Supplementary Figure 3**

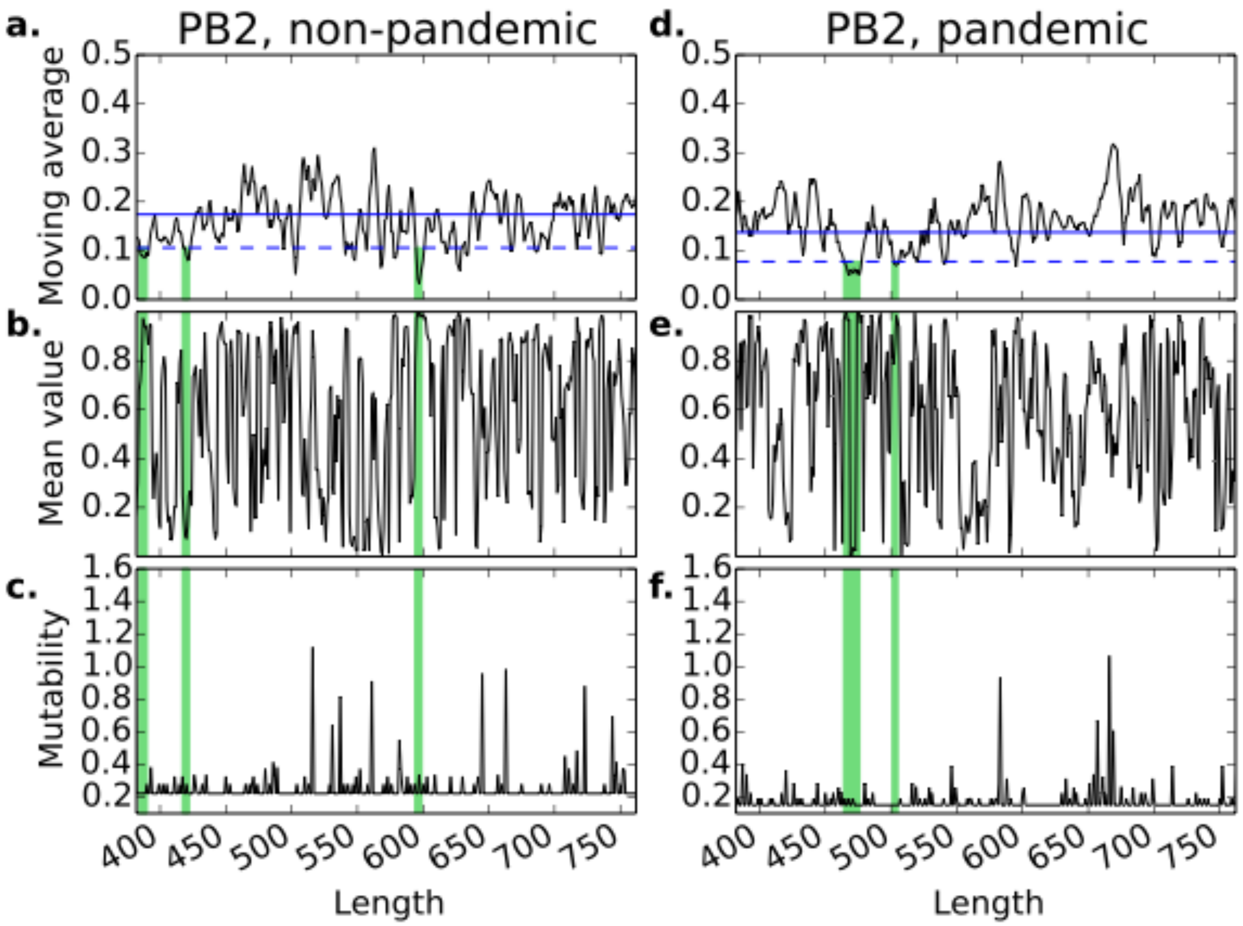

**Supplementary Figure 4**

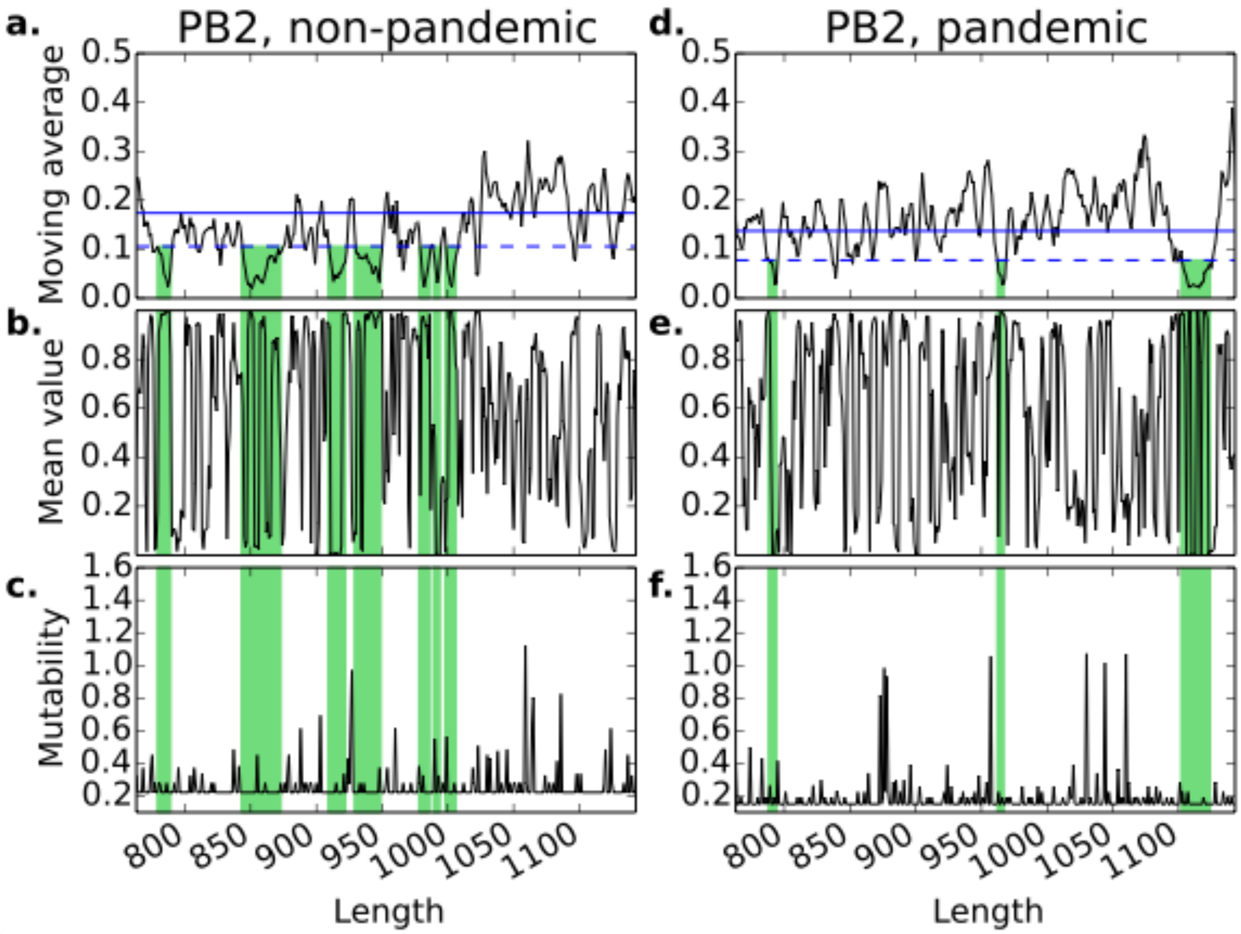

**Supplementary Figure 5**

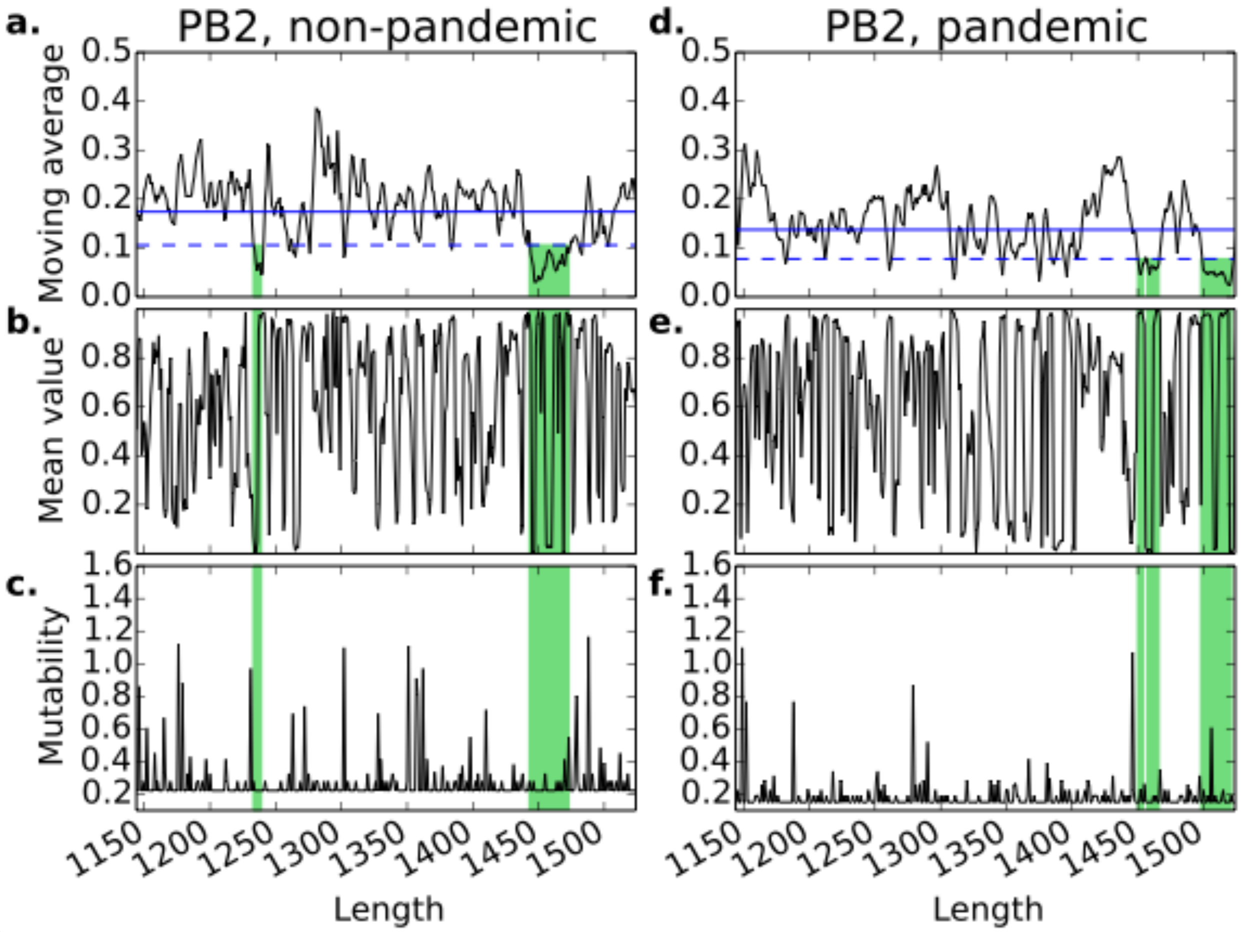

**Supplementary Figure 6**

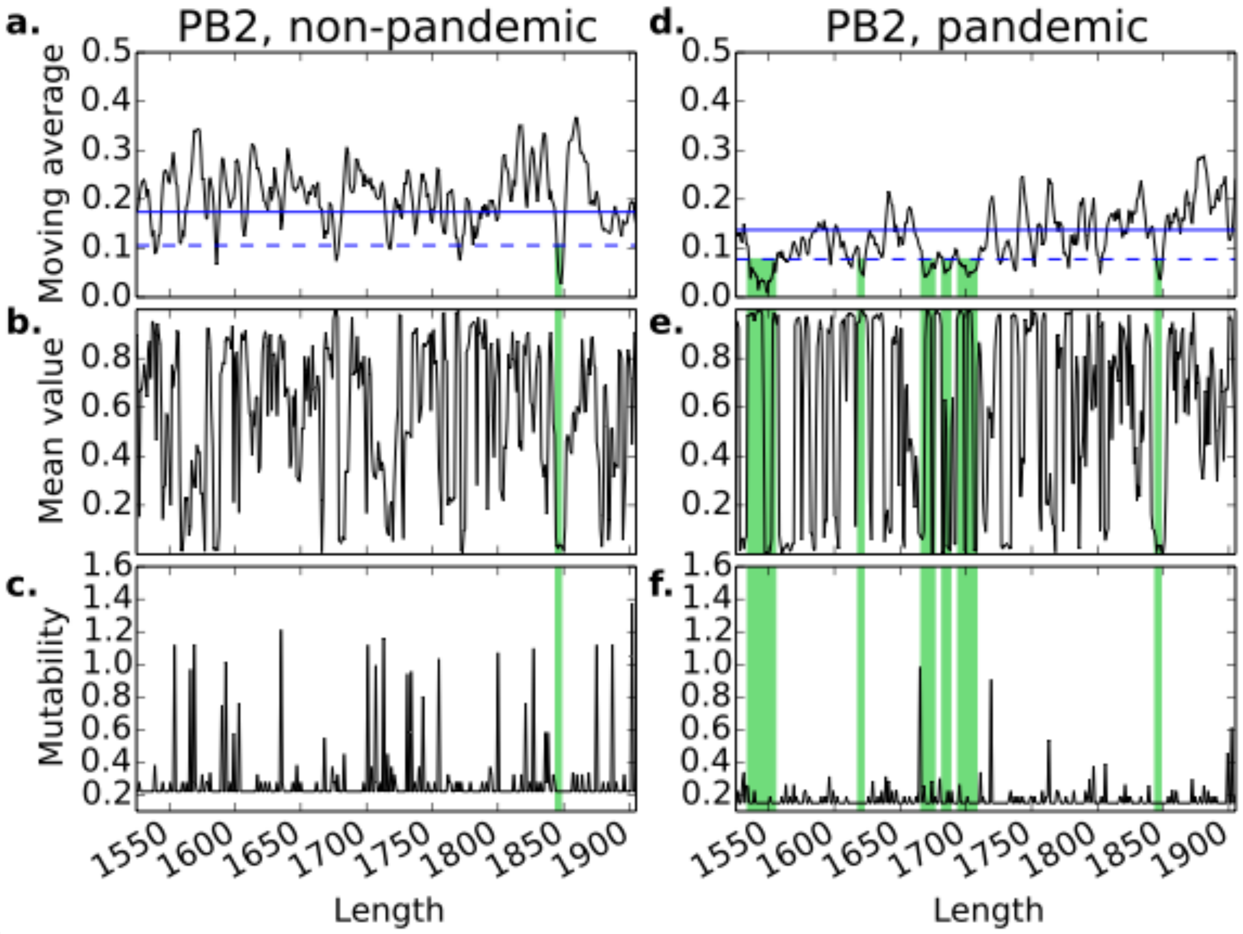

**Supplementary Figure 7**

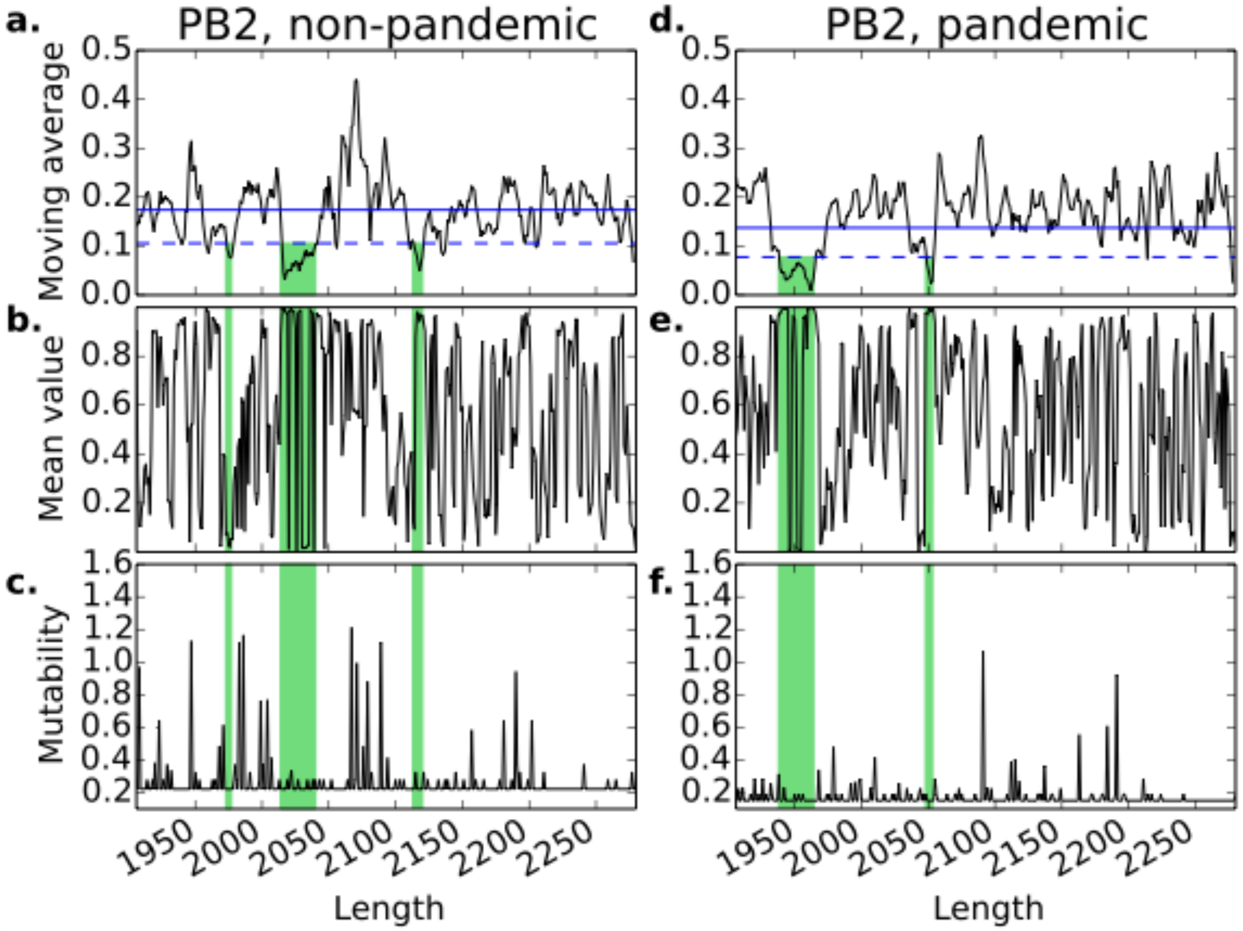

**Supplementary Figure 8**

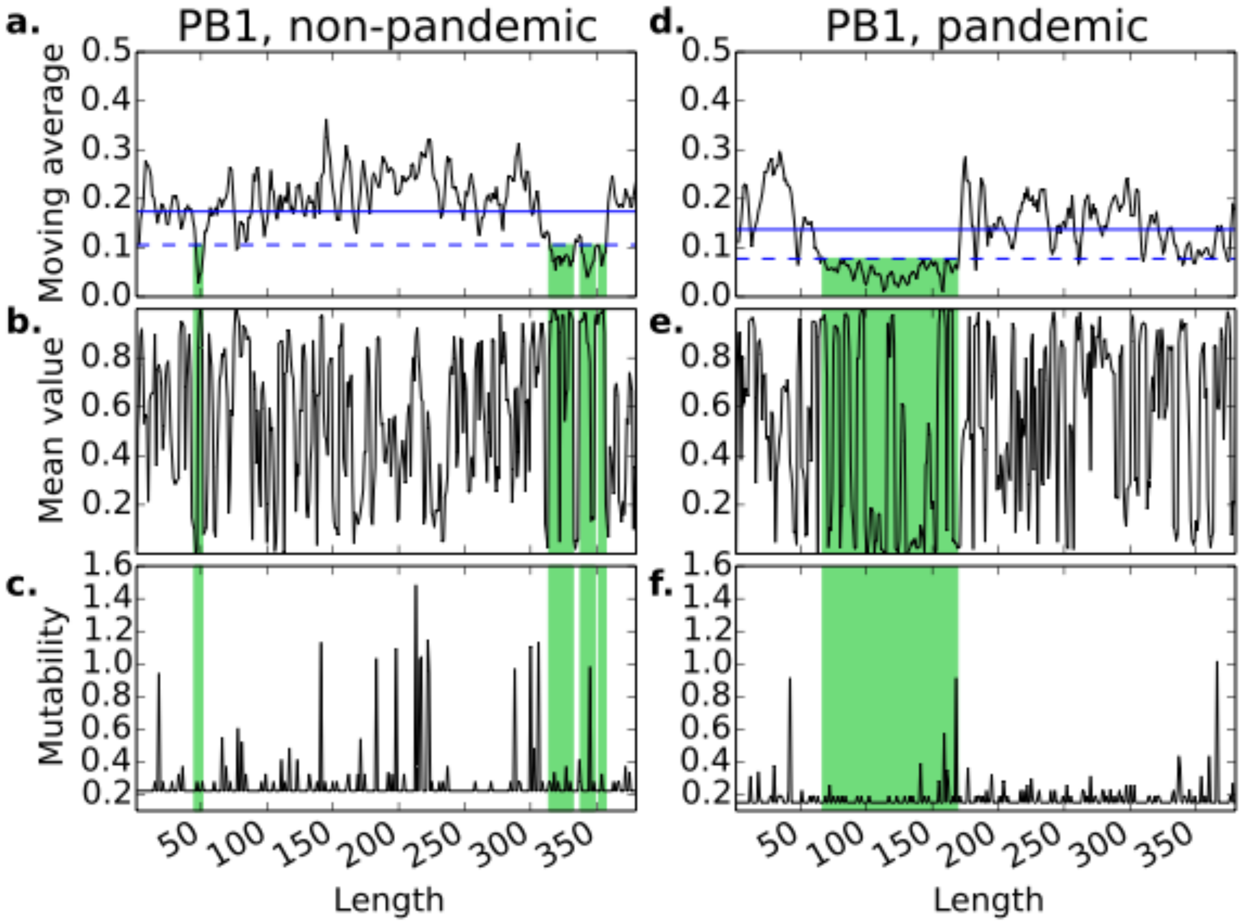

**Supplementary Figure 9**

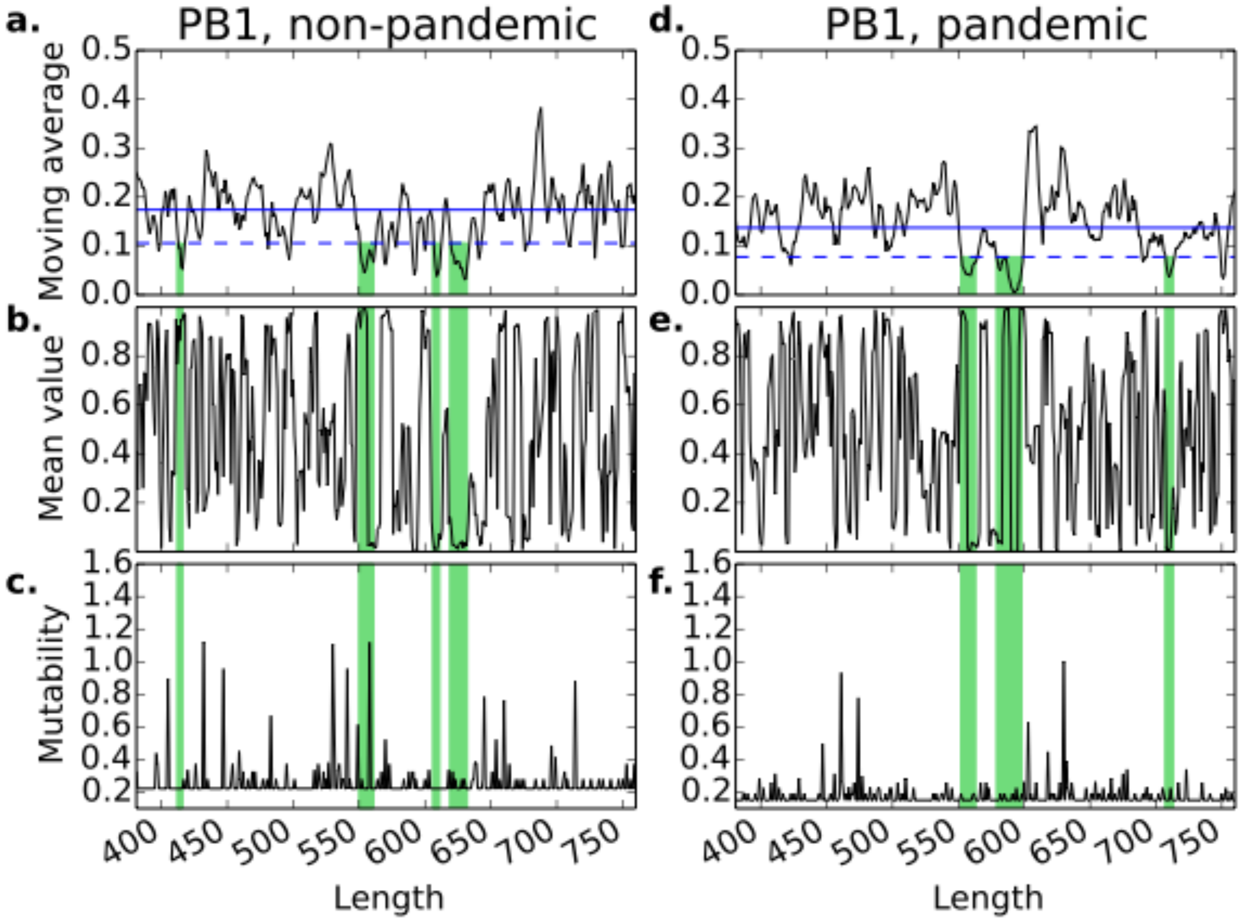

**Supplementary Figure 10**

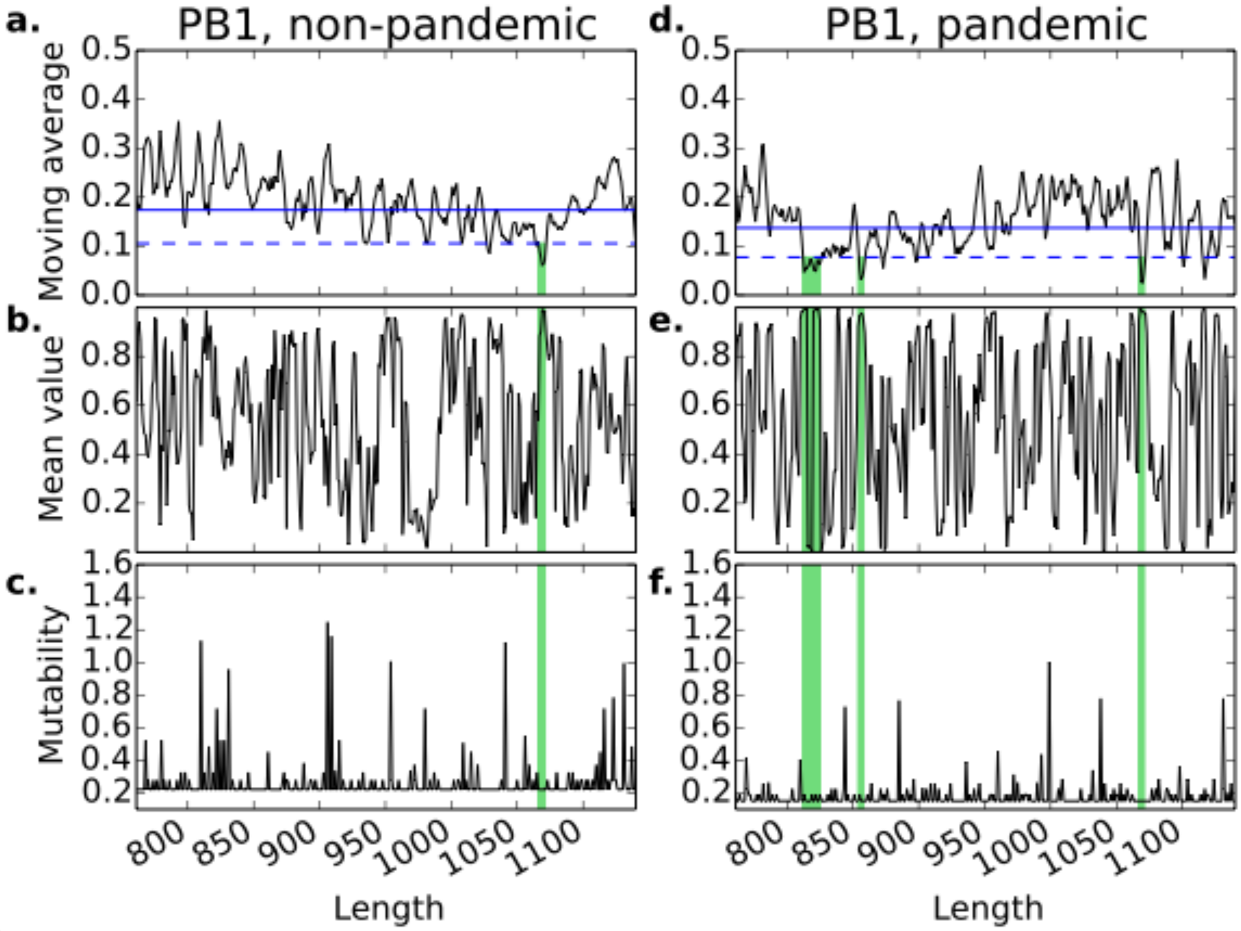

**Supplementary Figure 11**

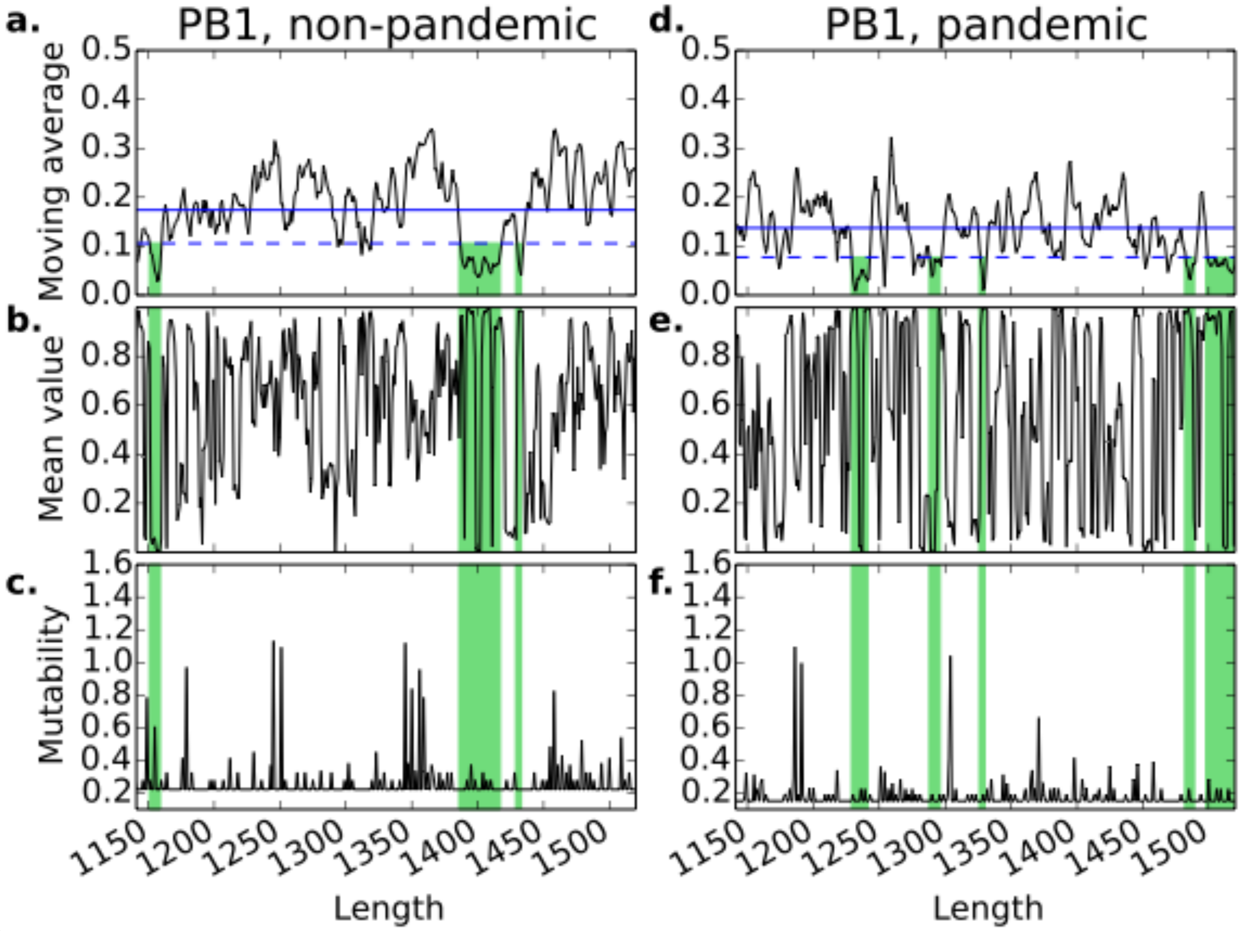

**Supplementary Figure 12**

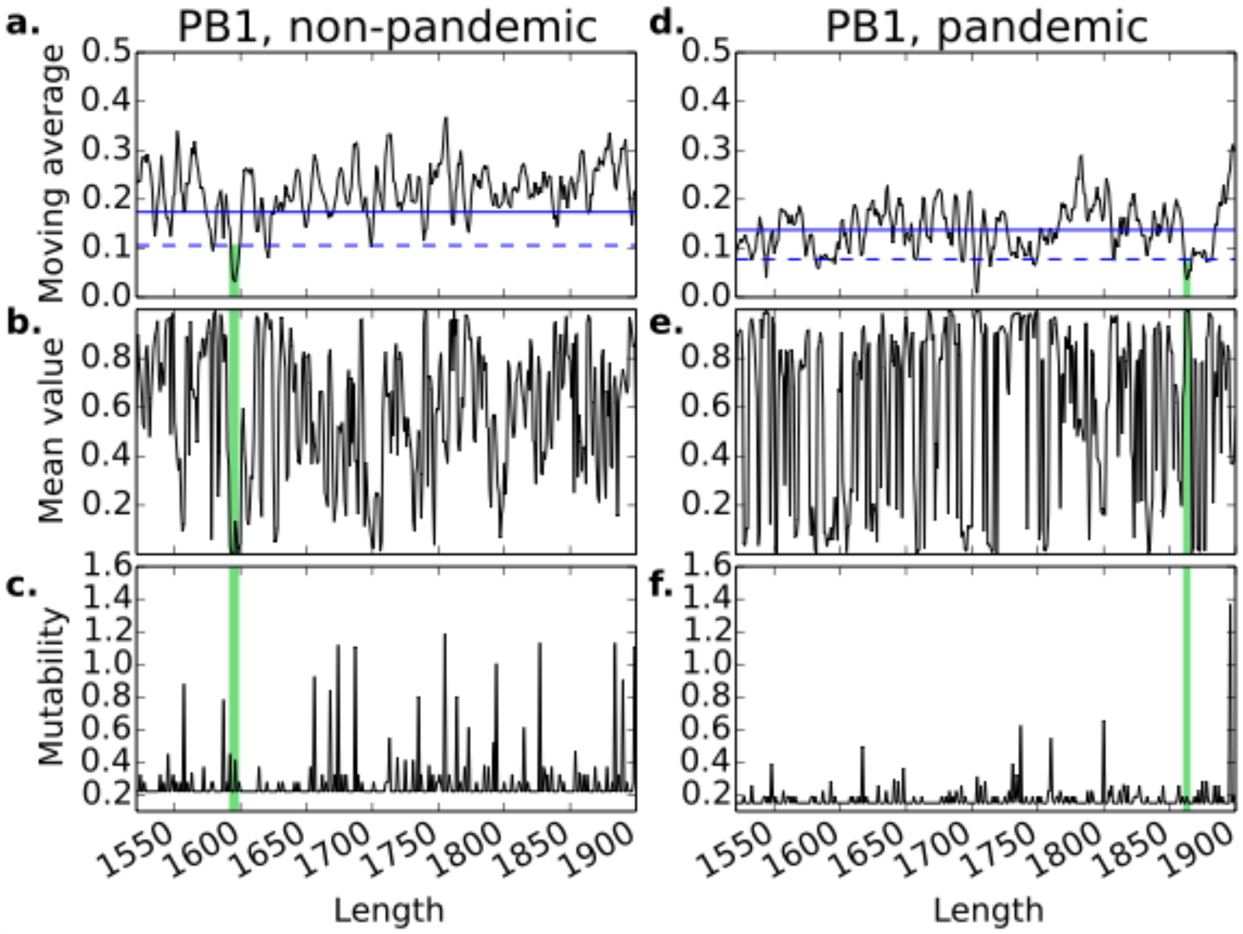

**Supplementary Figure 13**

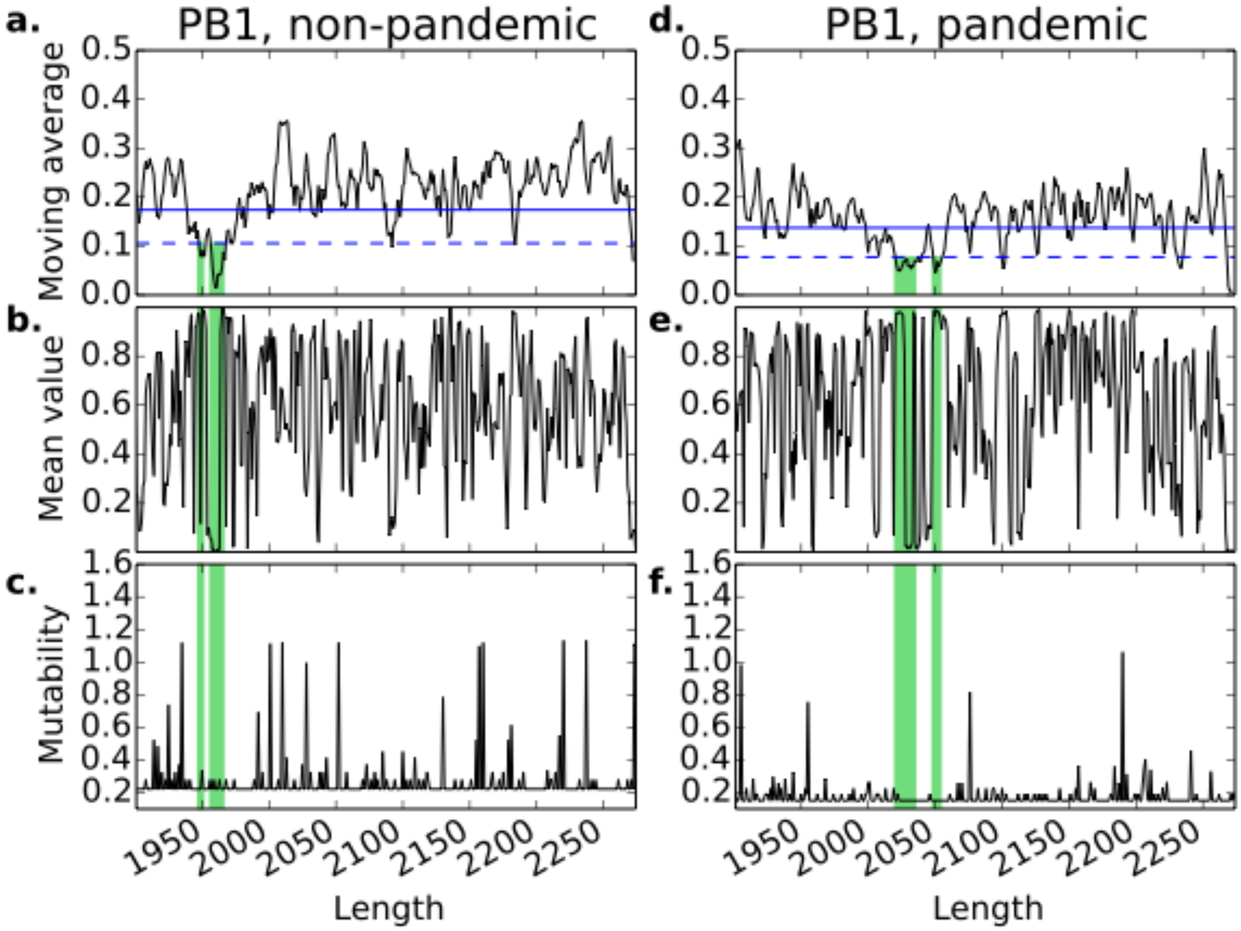

**Supplementary Figure 14**

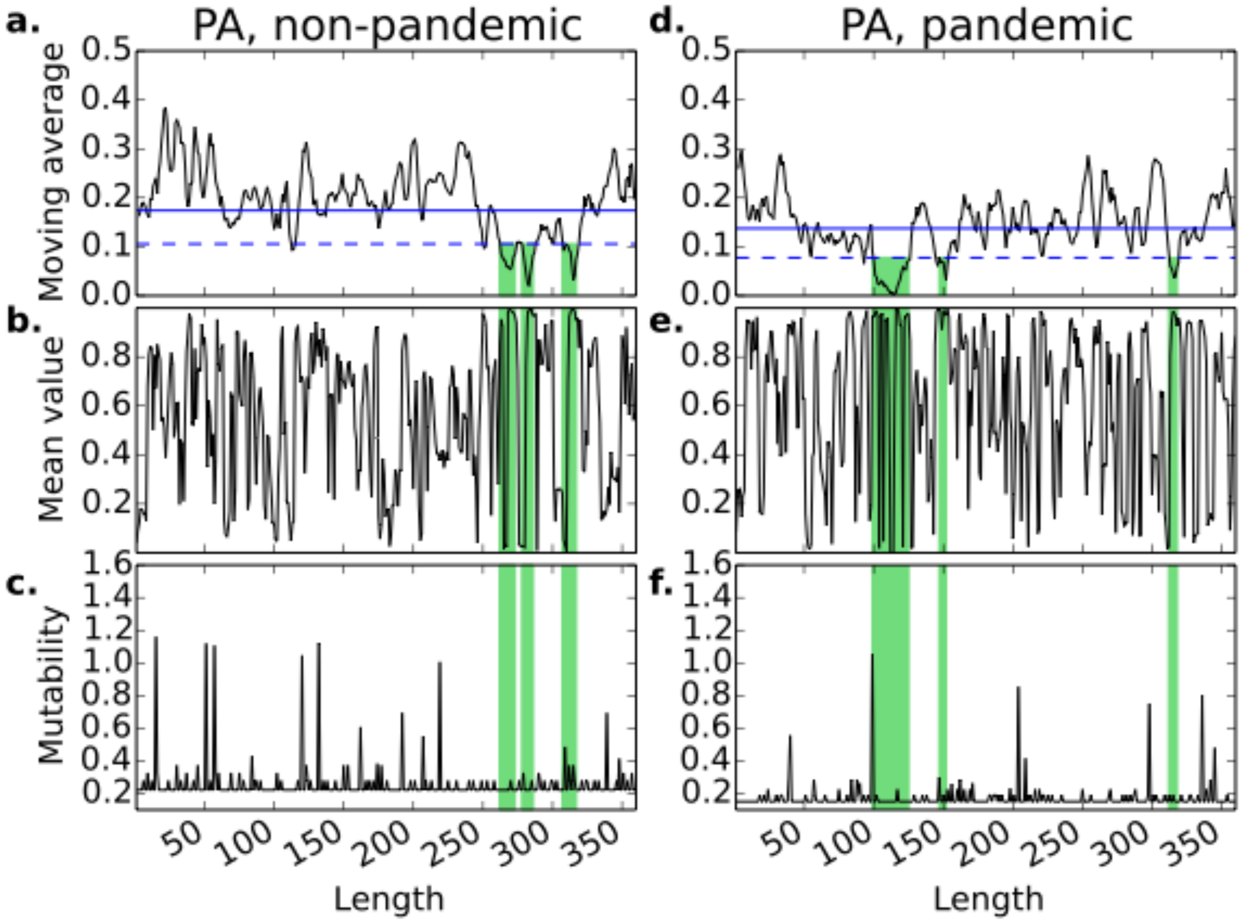

**Supplementary Figure 15**

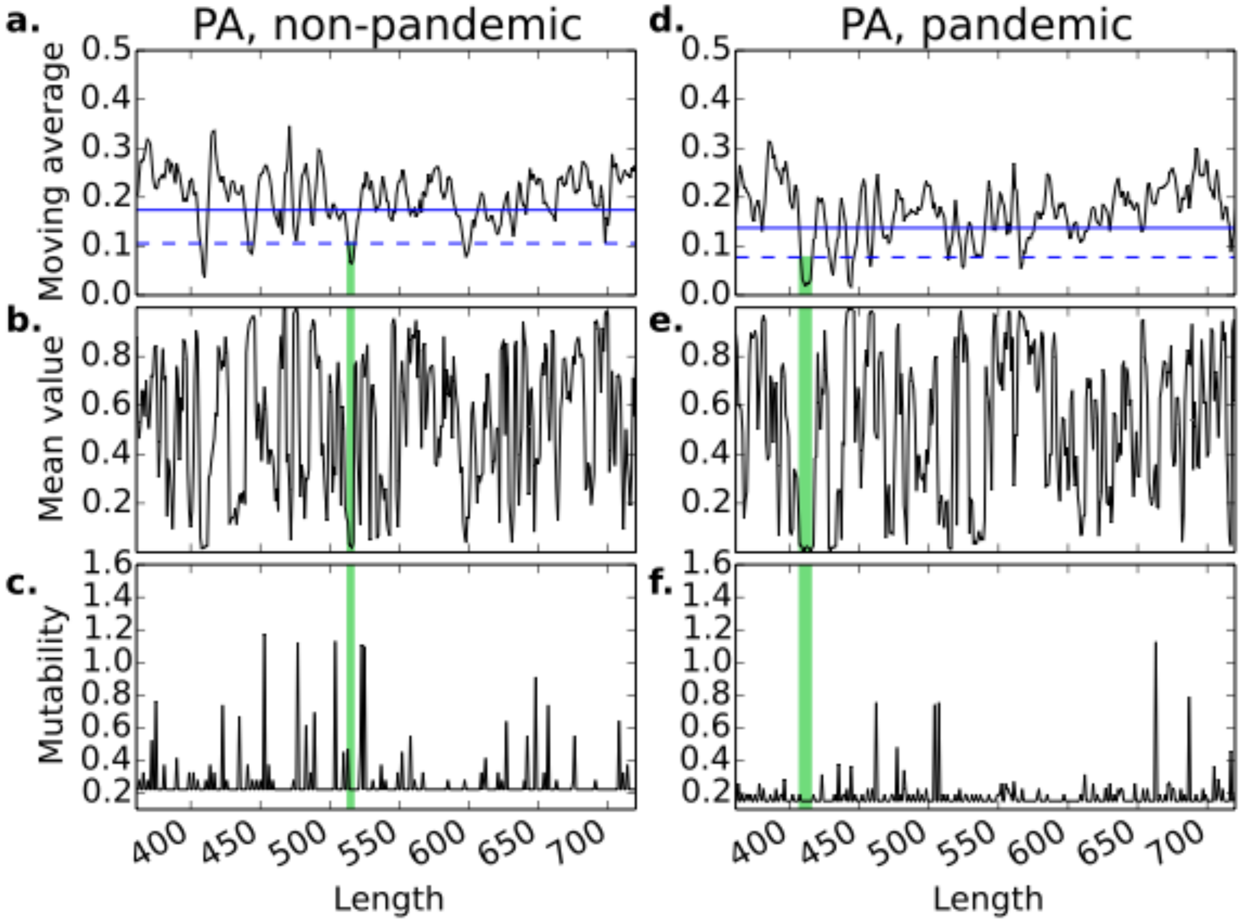

**Supplementary Figure 16**

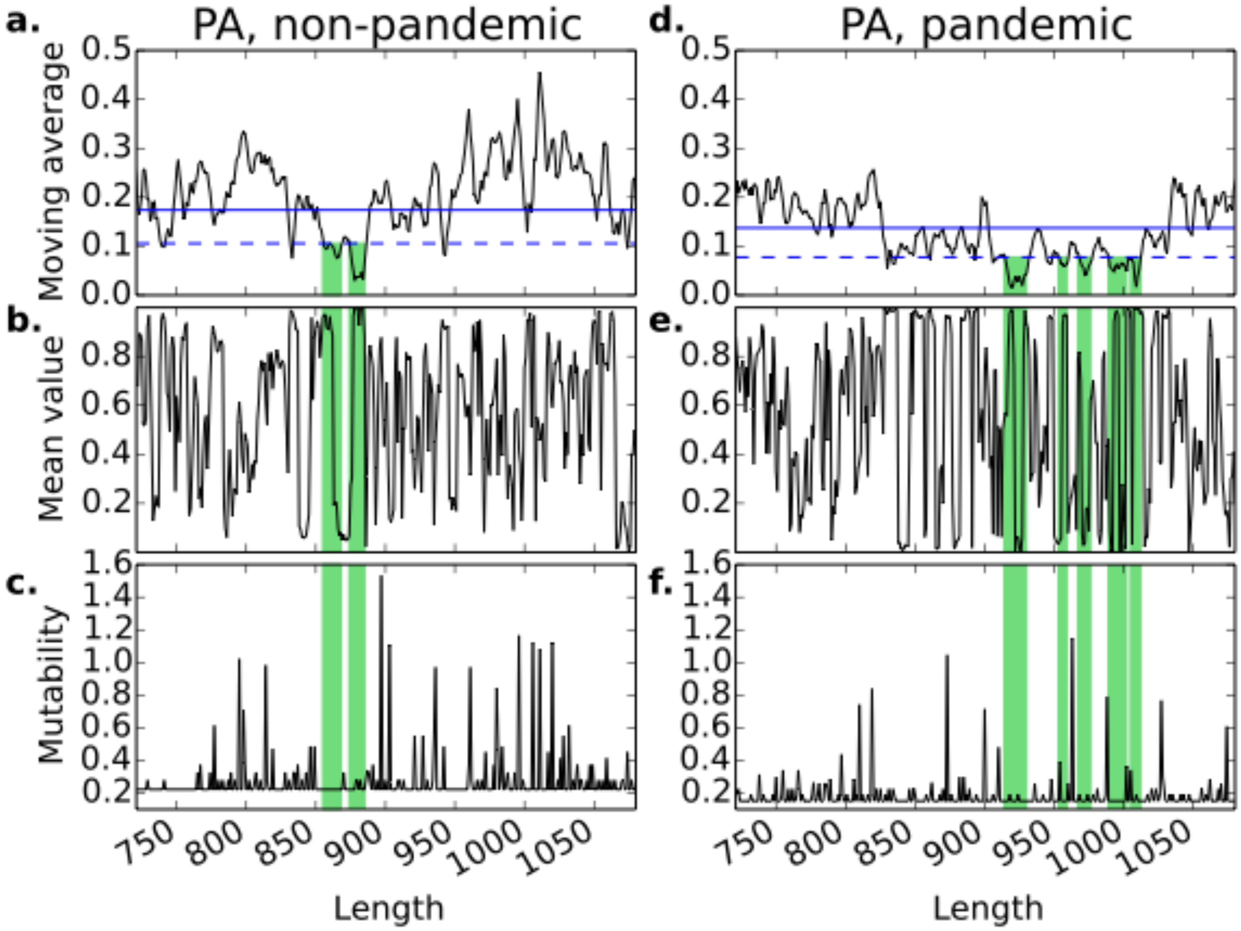

**Supplementary Figure 17**

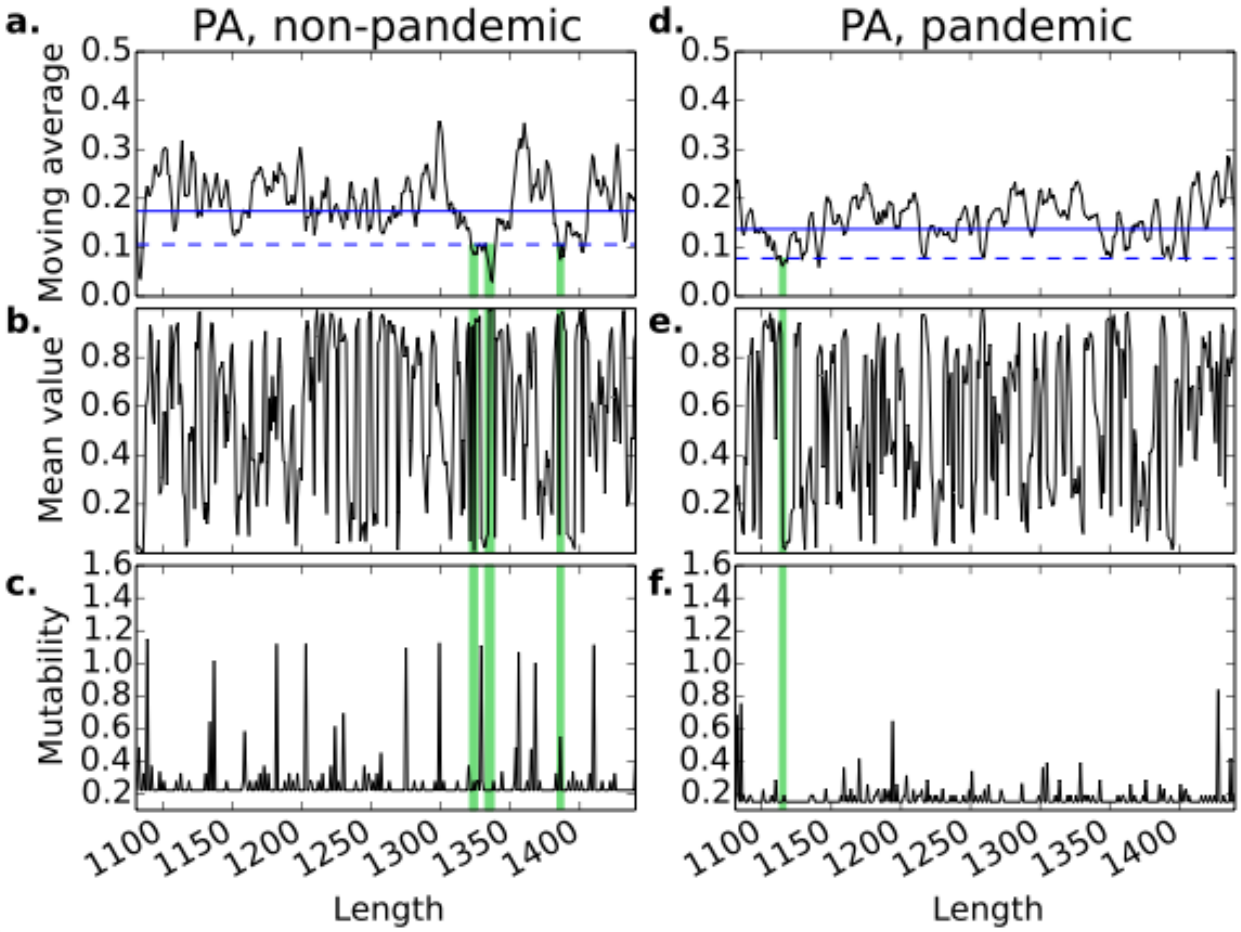

**Supplementary Figure 18**

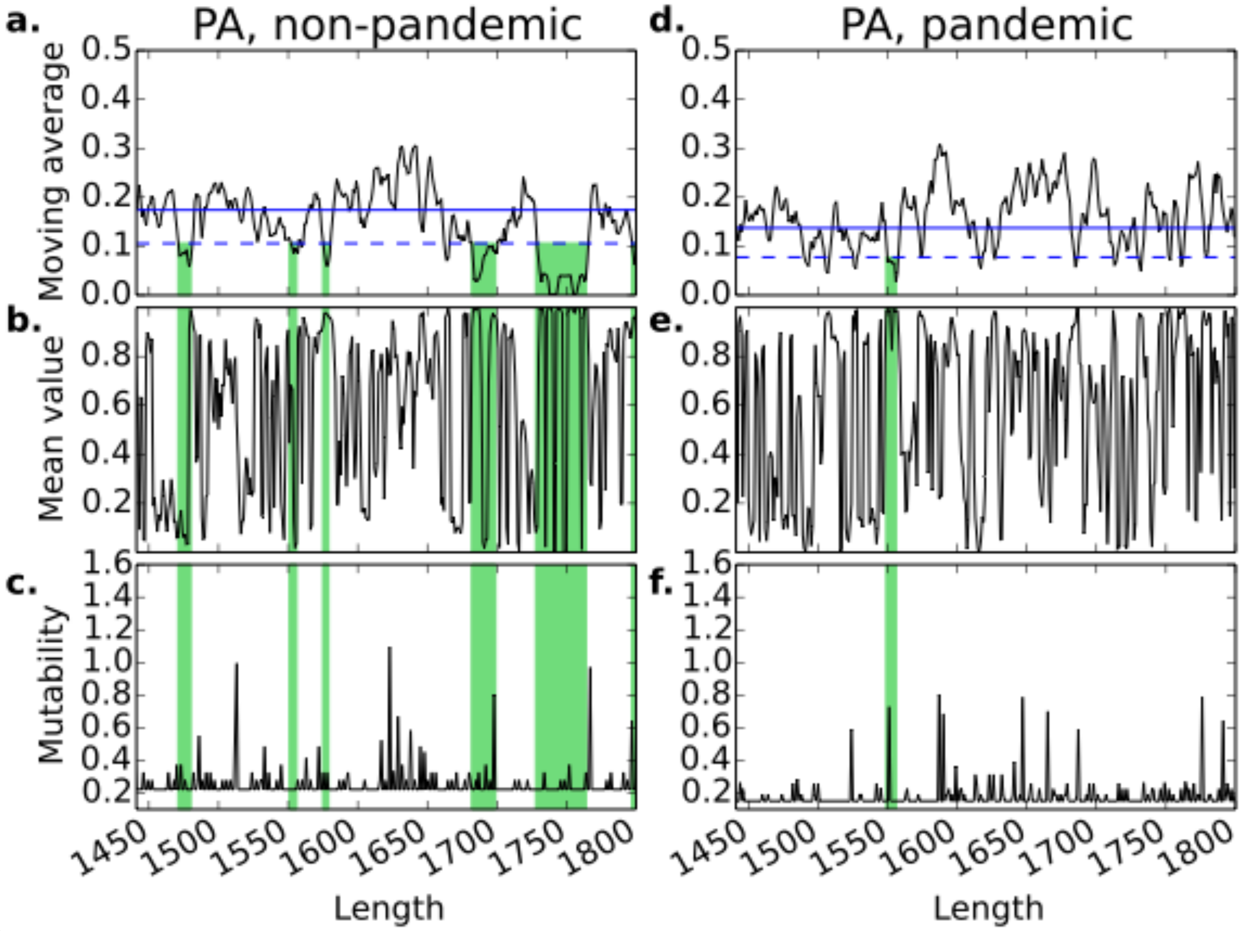

**Supplementary Figure 19**

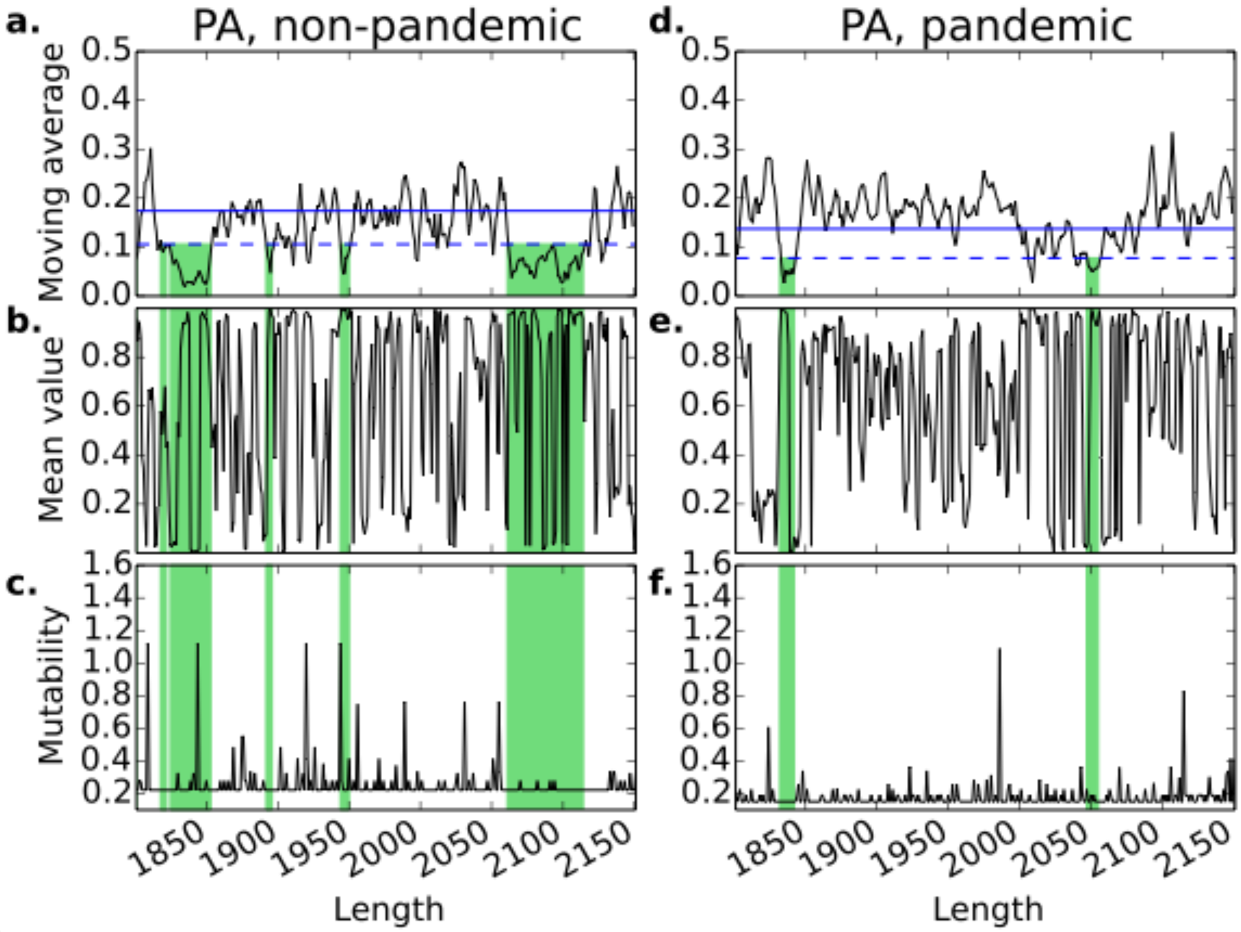

**Supplementary Figure 20**

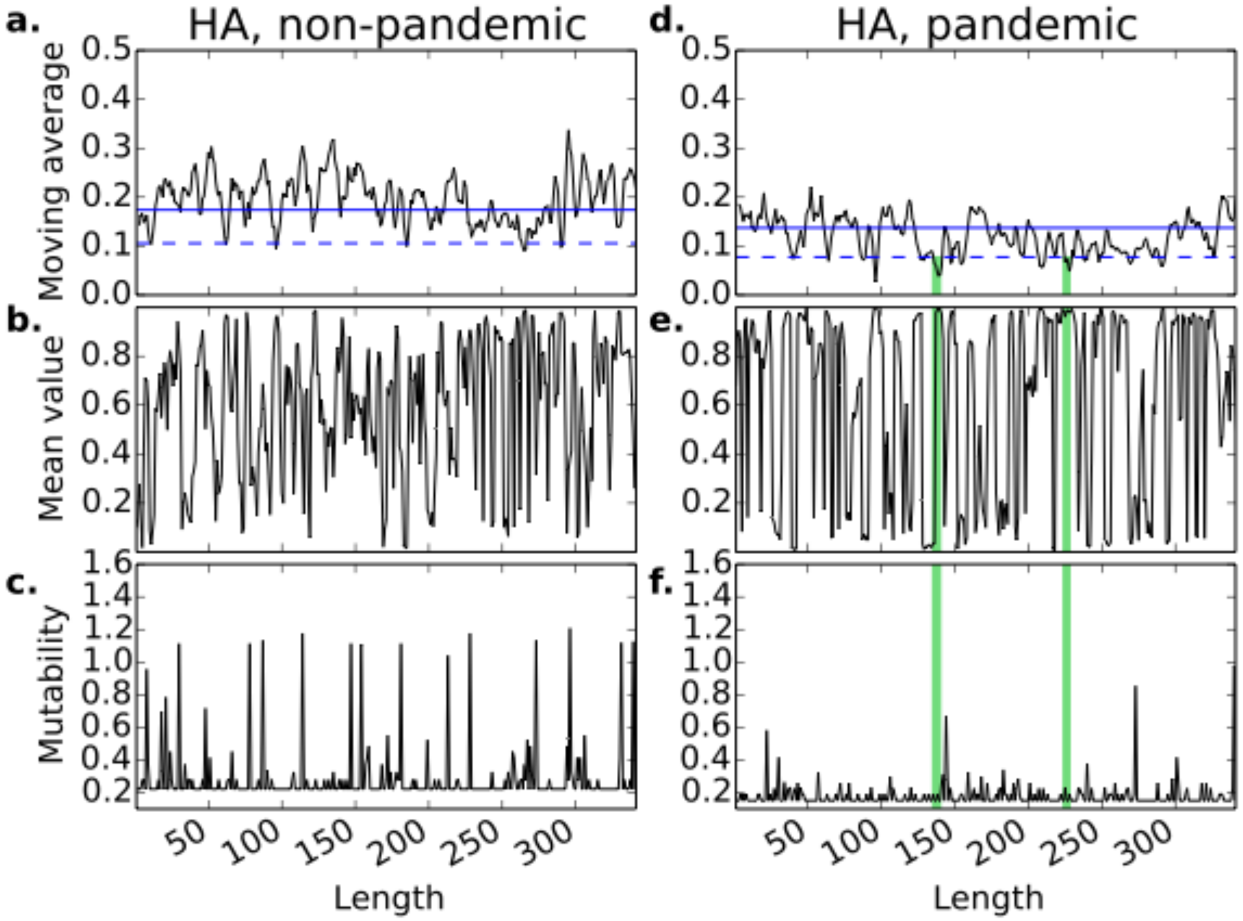

**Supplementary Figure 21**

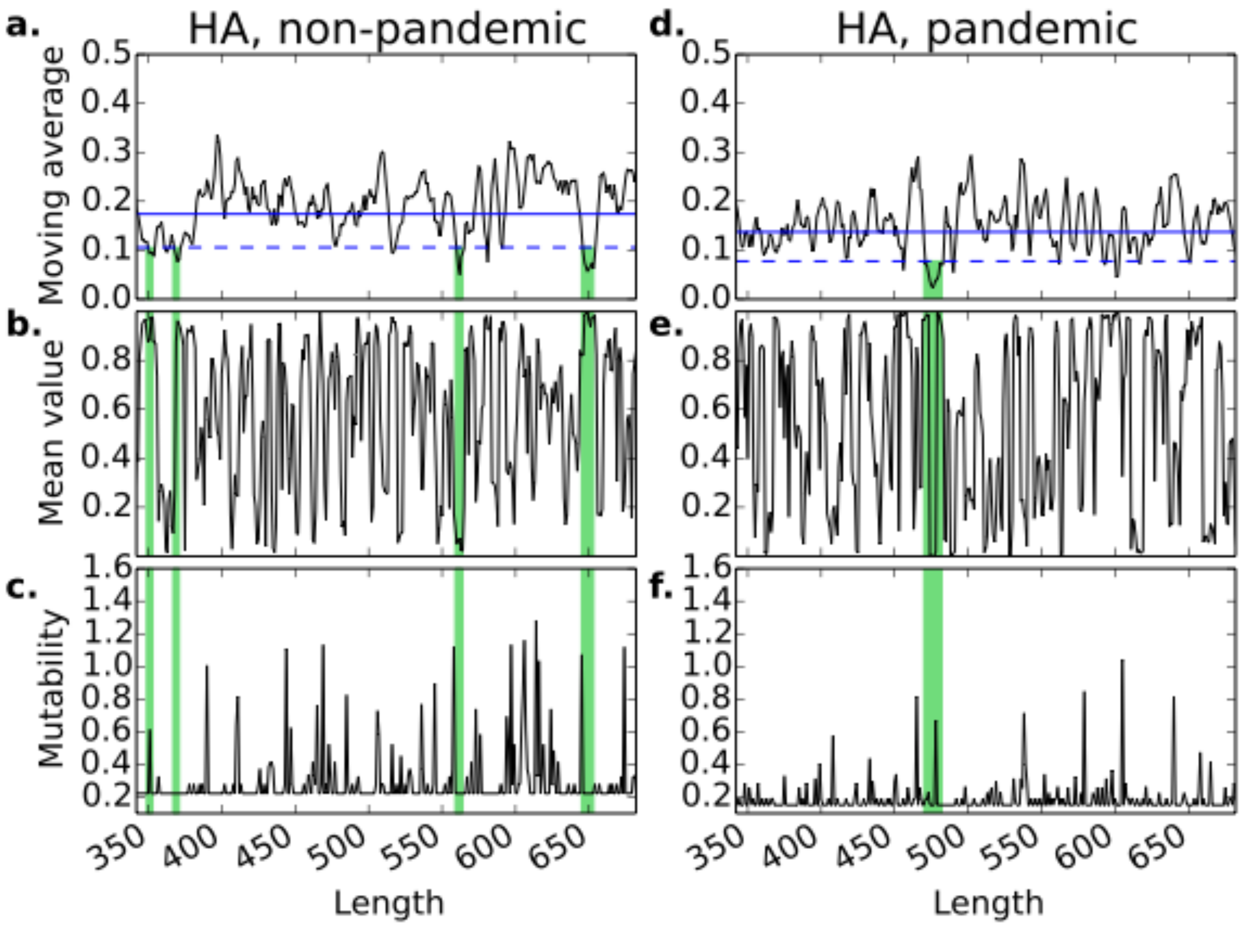

**Supplementary Figure 22**

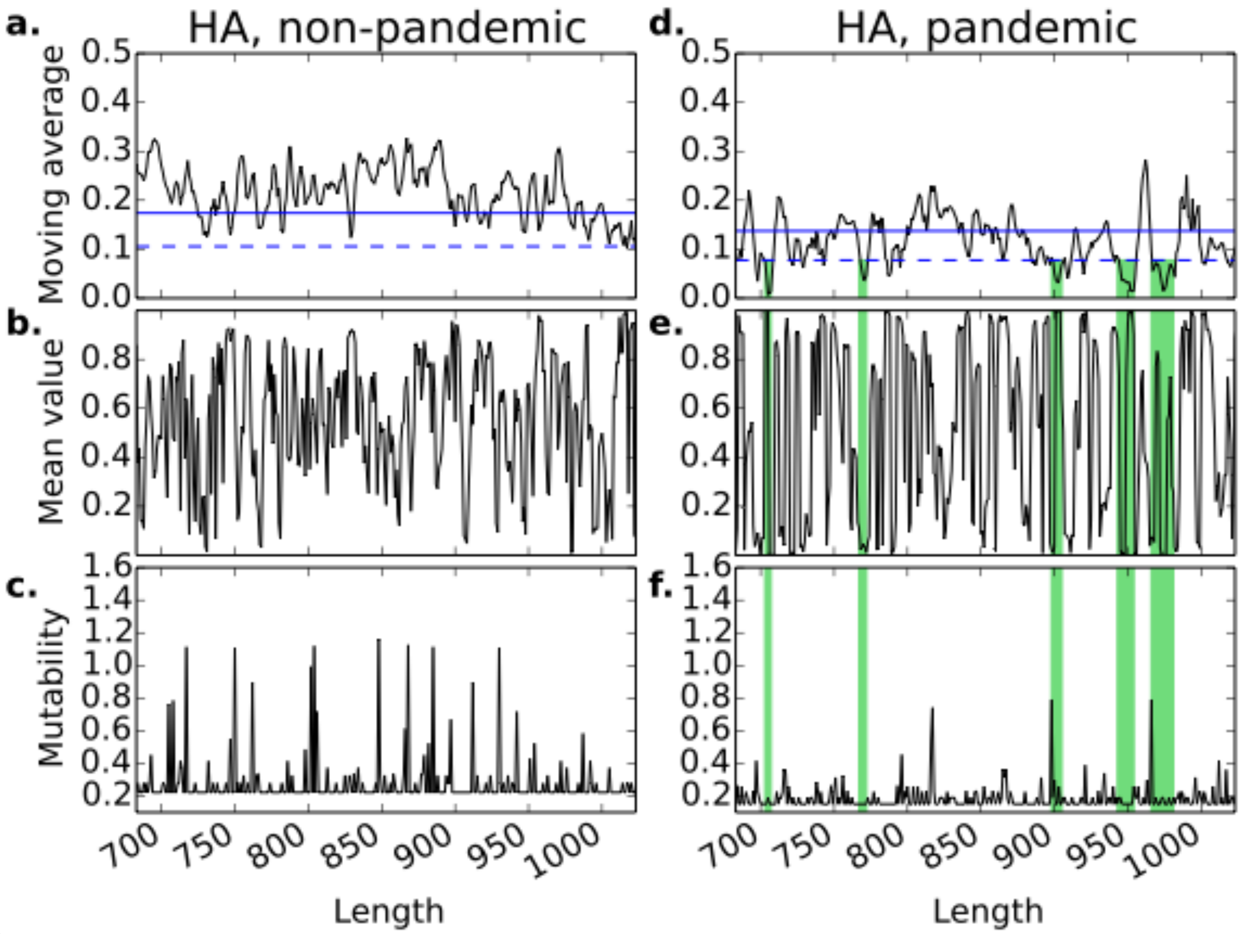

**Supplementary Figure 23**

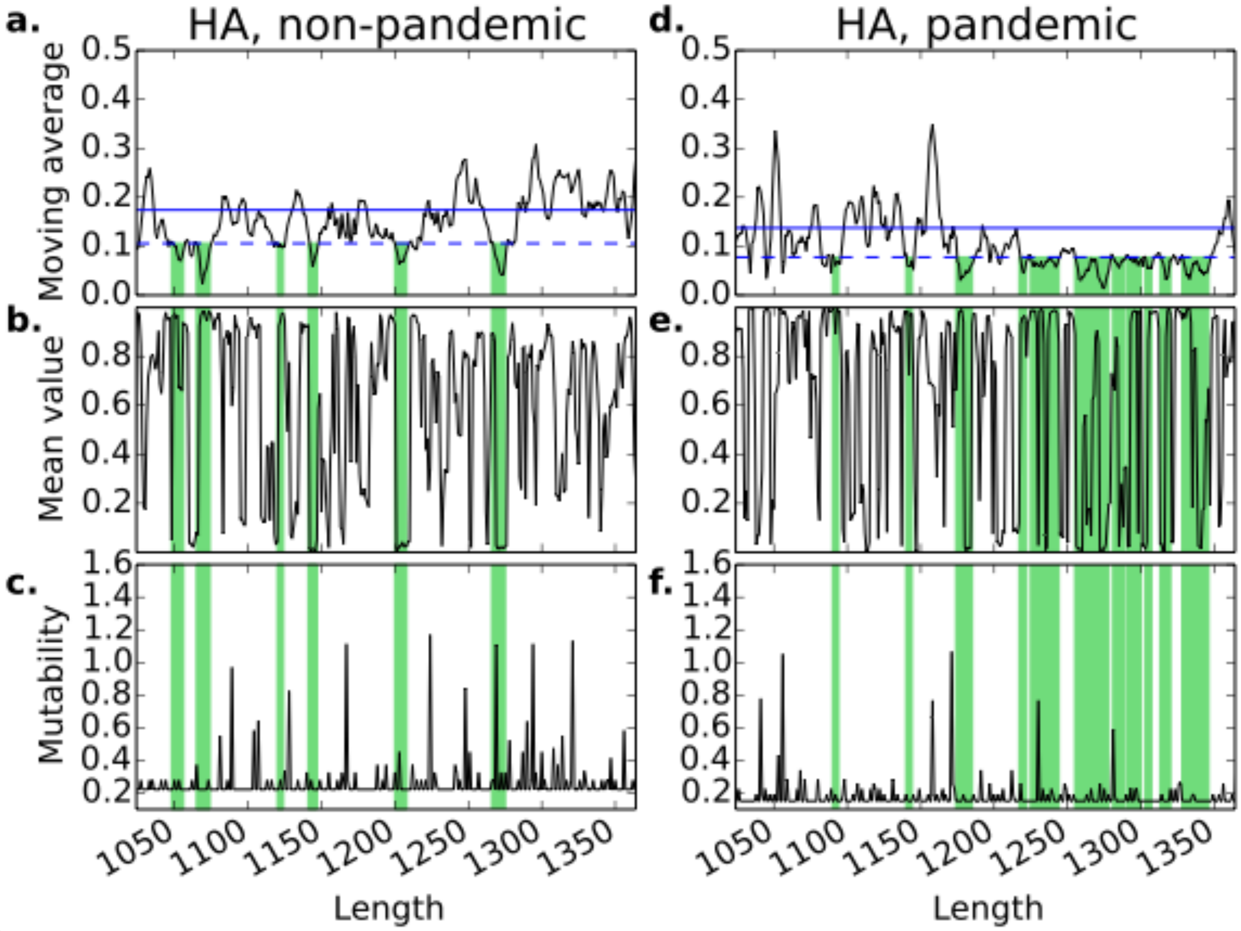

**Supplementary Figure 24**

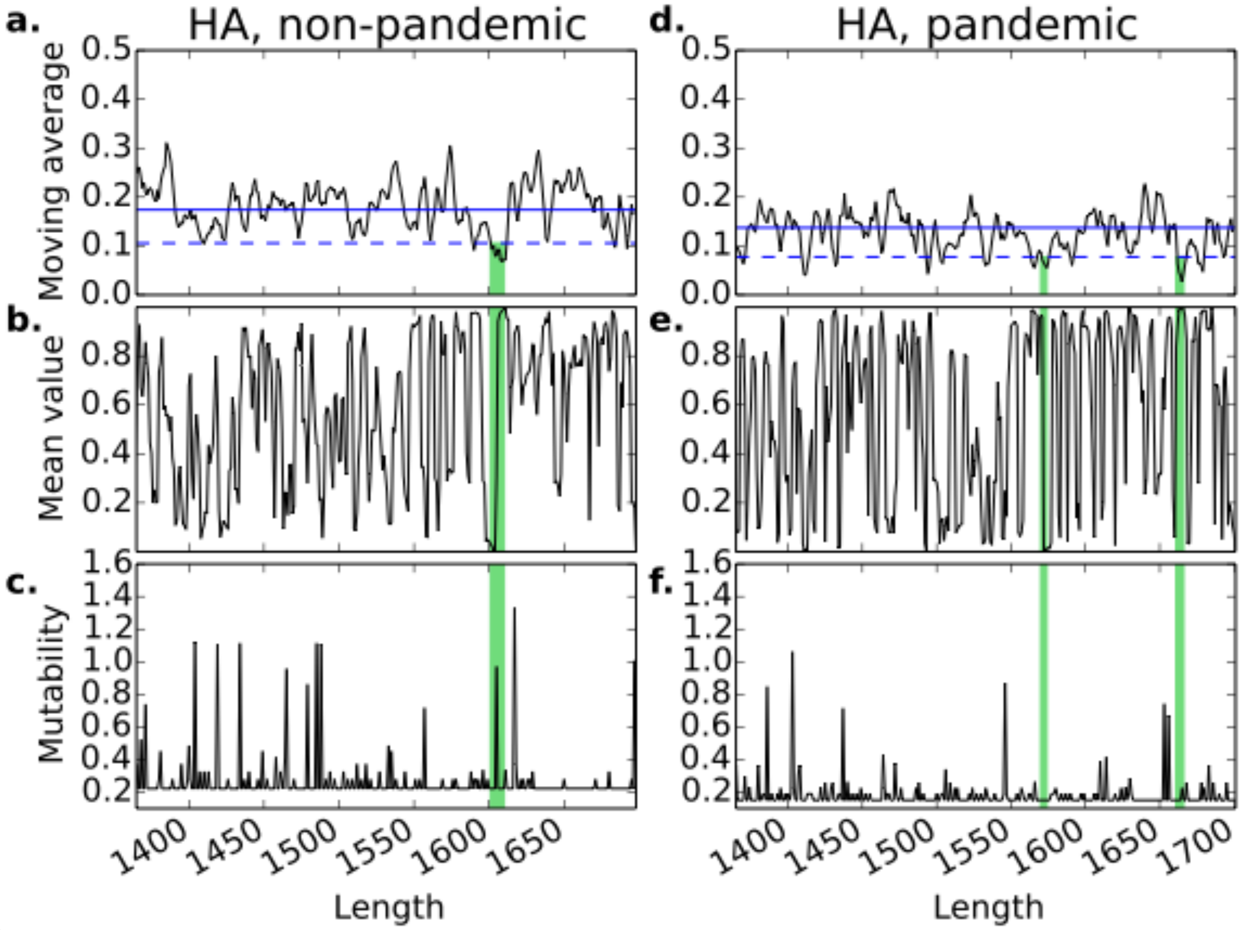

**Supplementary Figure 25**

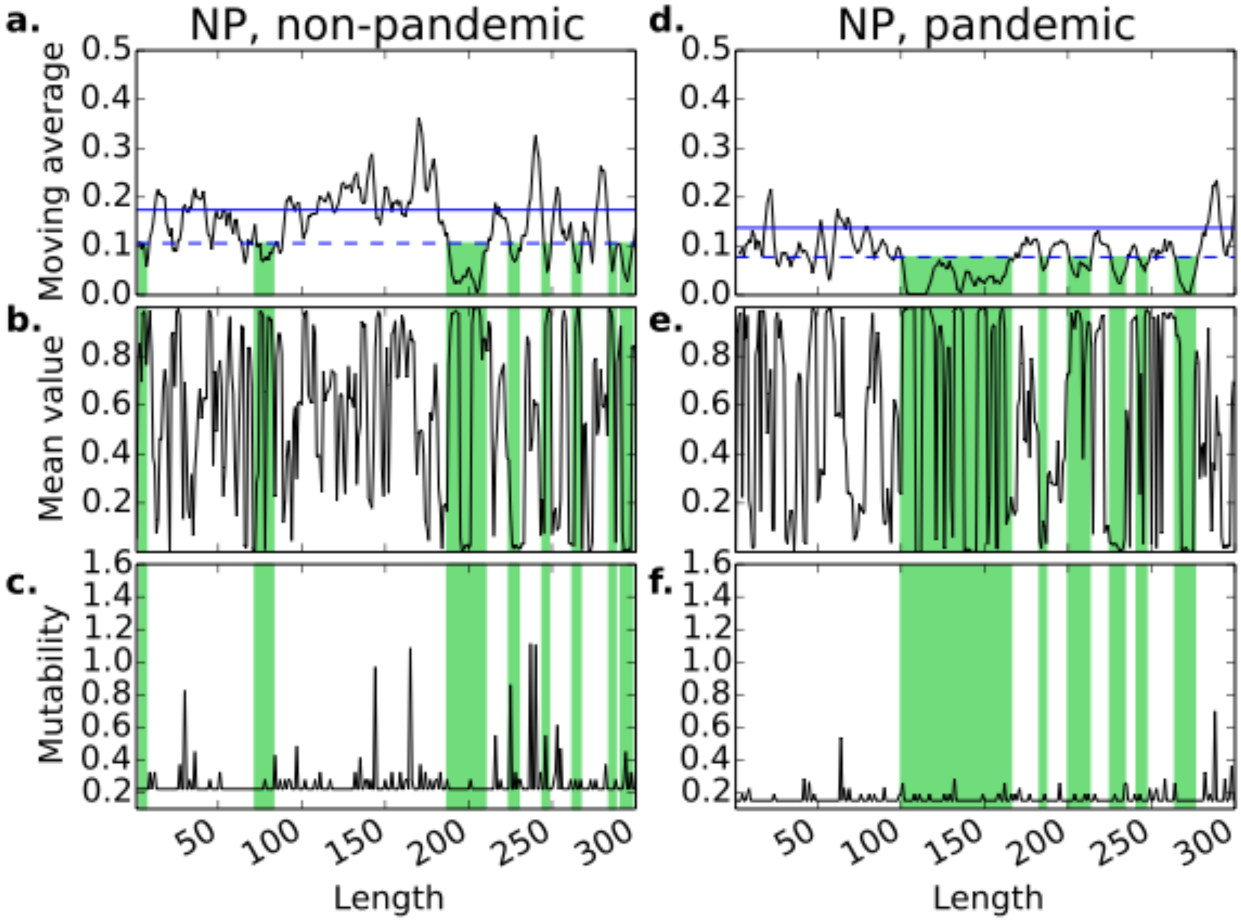

**Supplementary Figure 26**

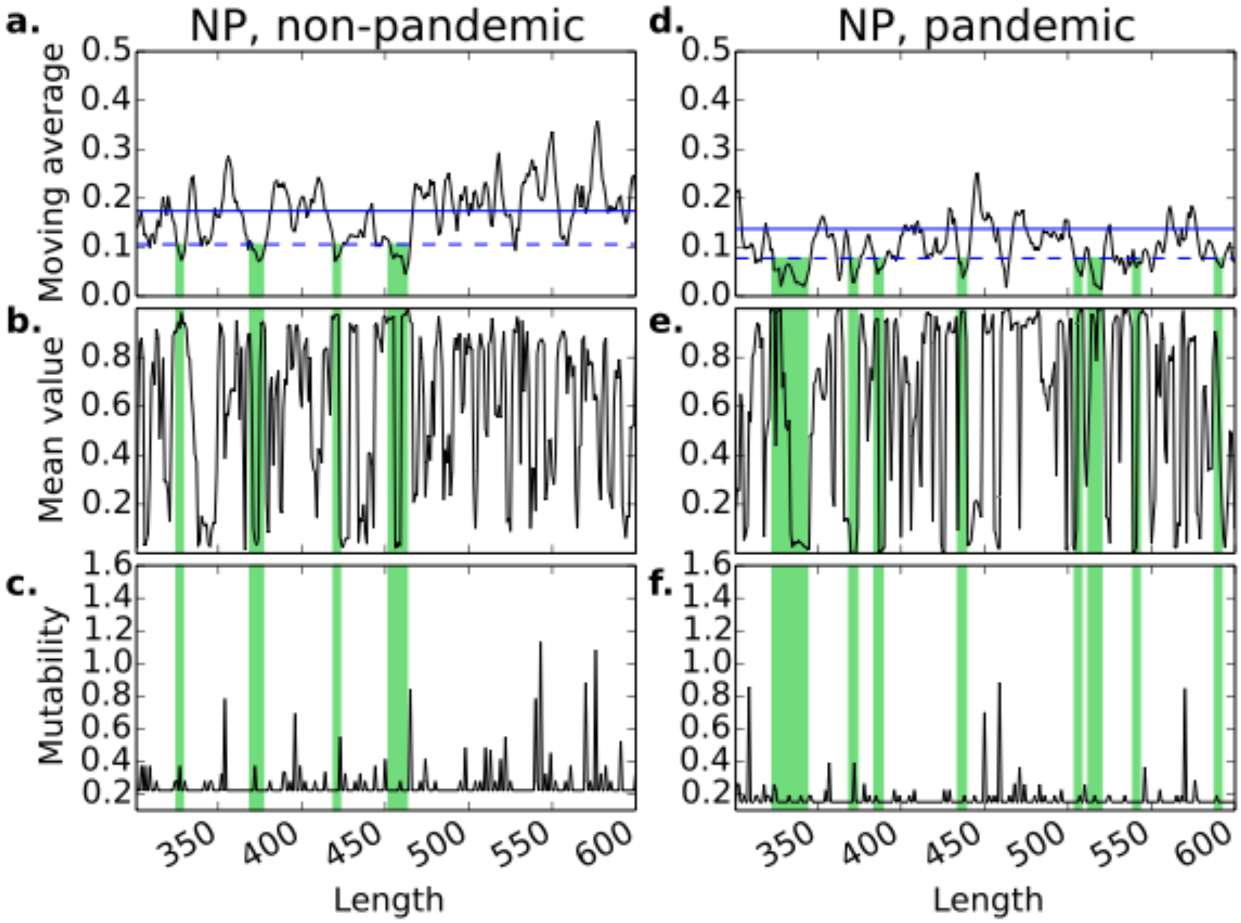

**Supplementary Figure 27**

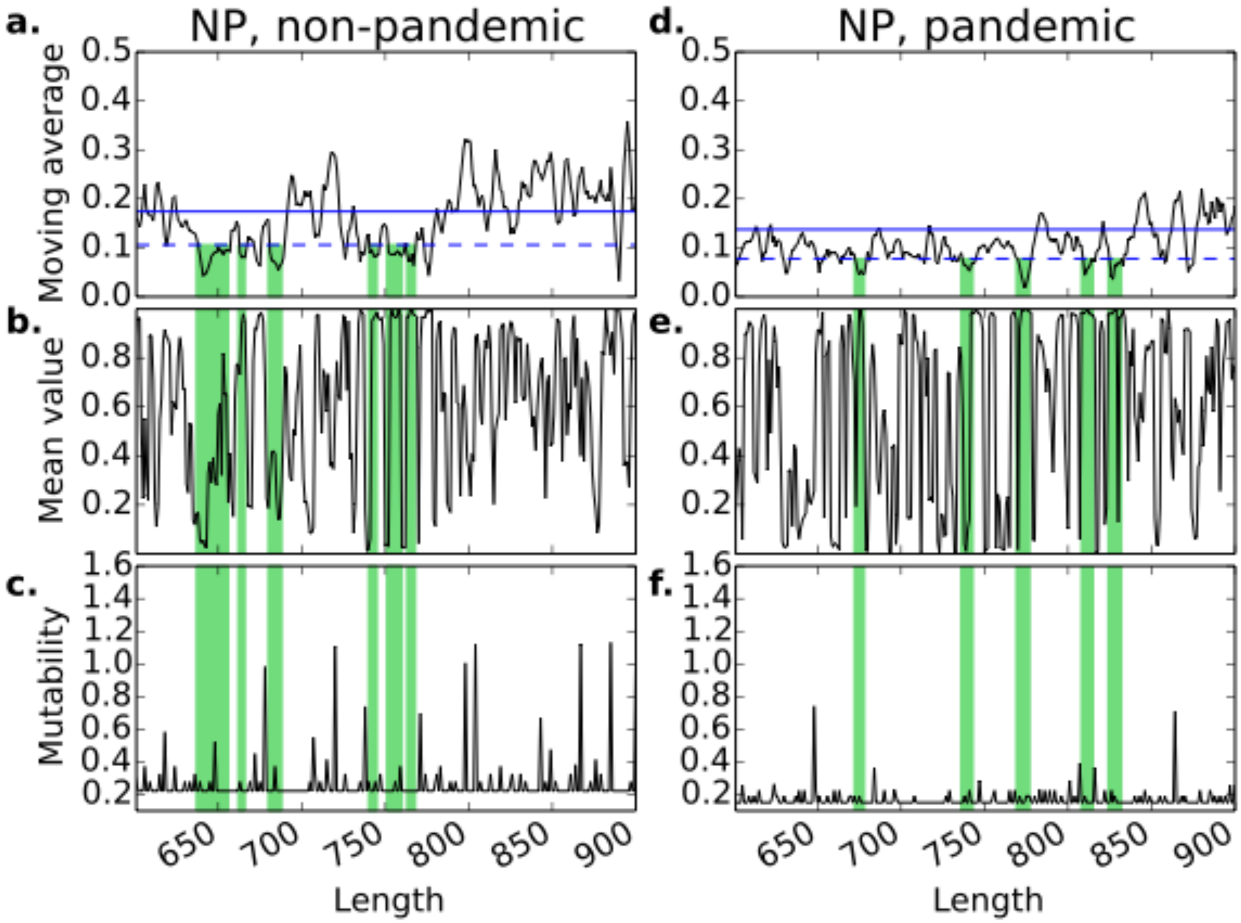

**Supplementary Figure 28**

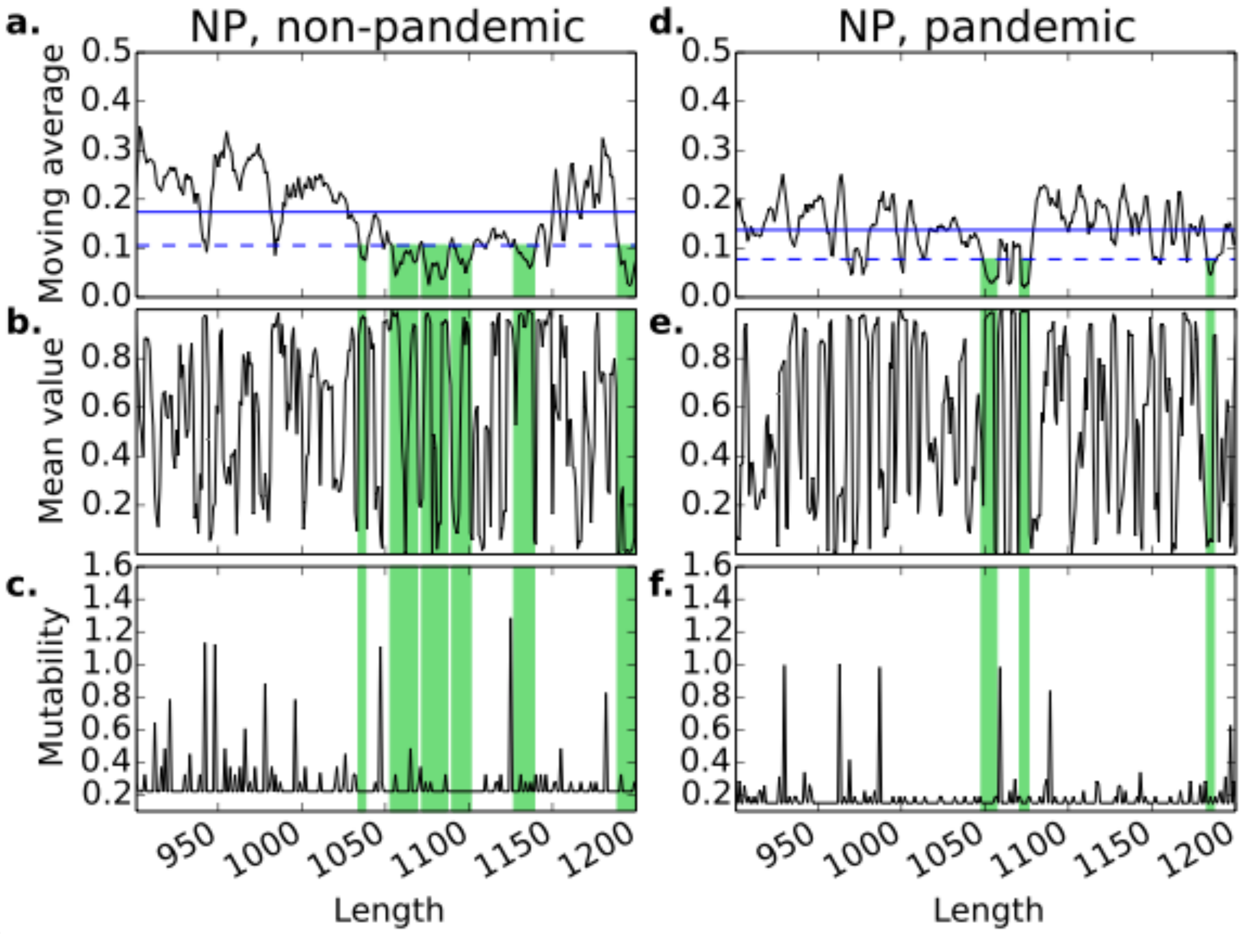

**Supplementary Figure 29**

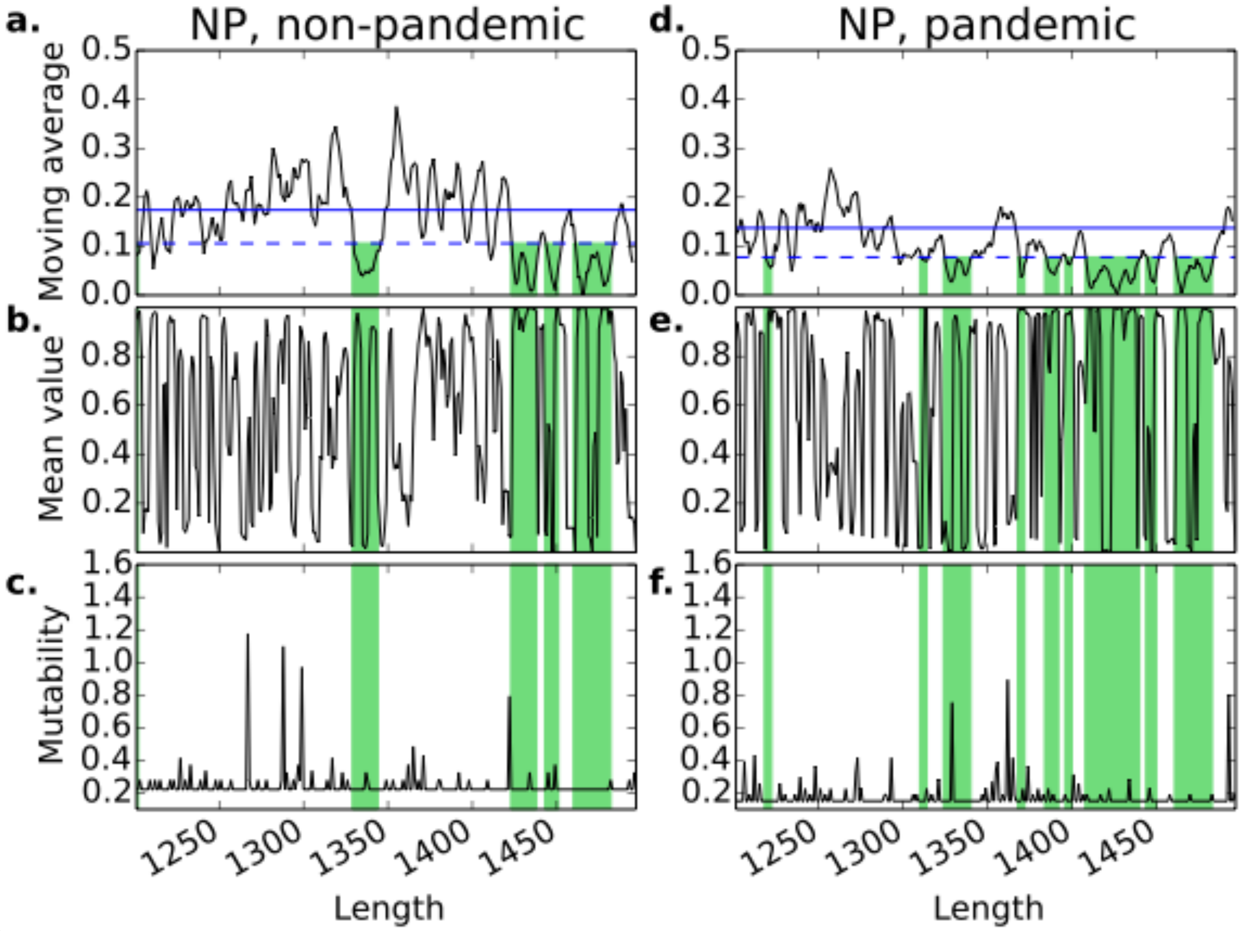

**Supplementary Figure 30**

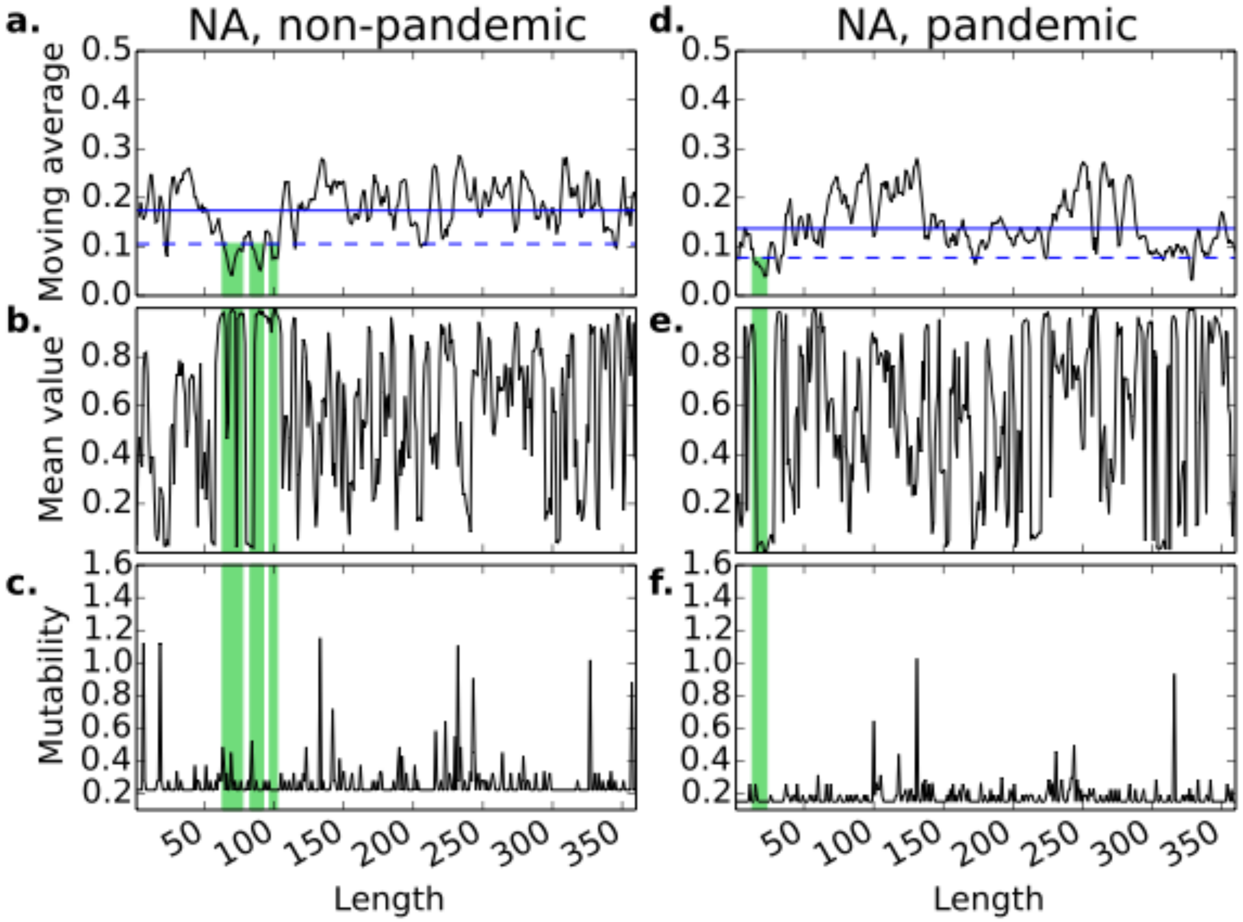

**Supplementary Figure 31**

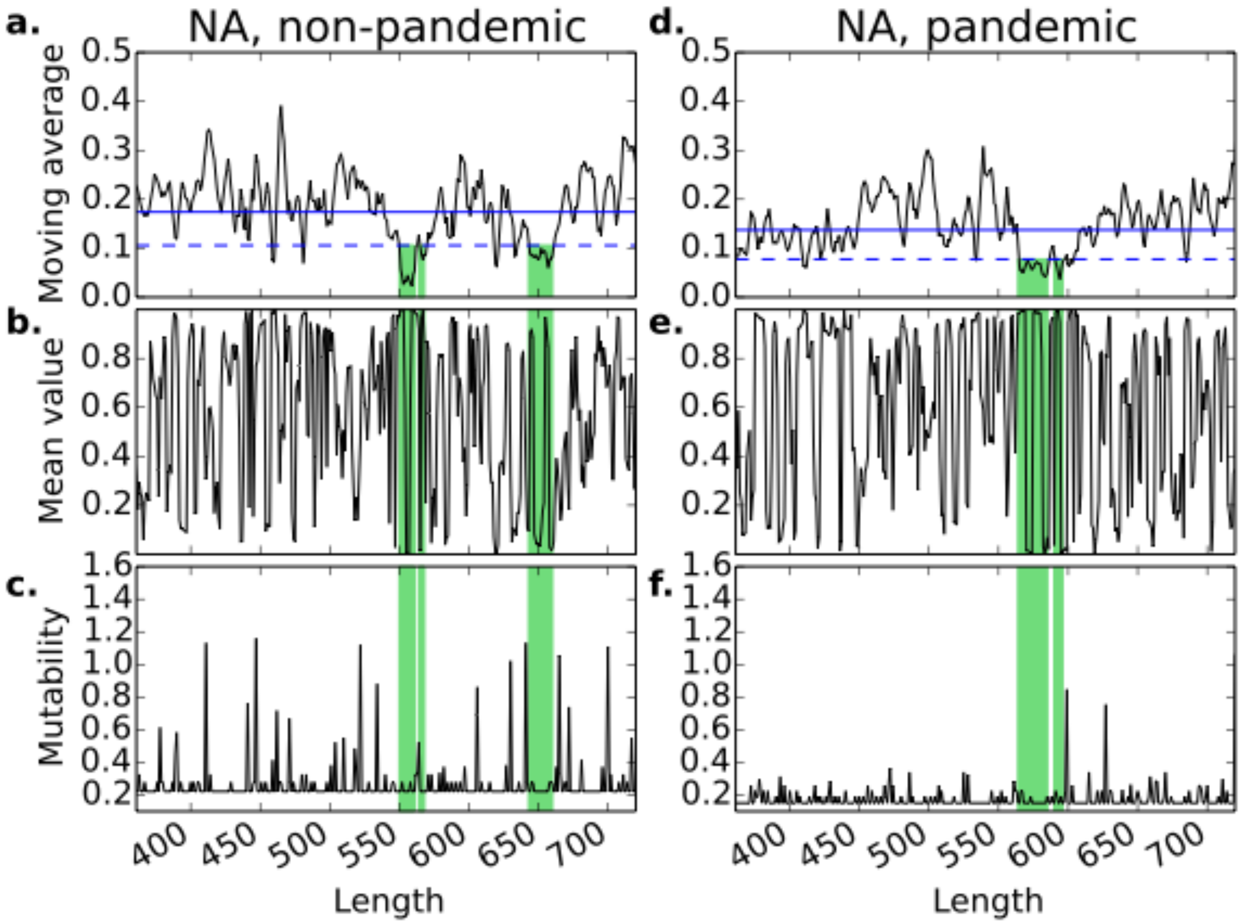

**Supplementary Figure 32**

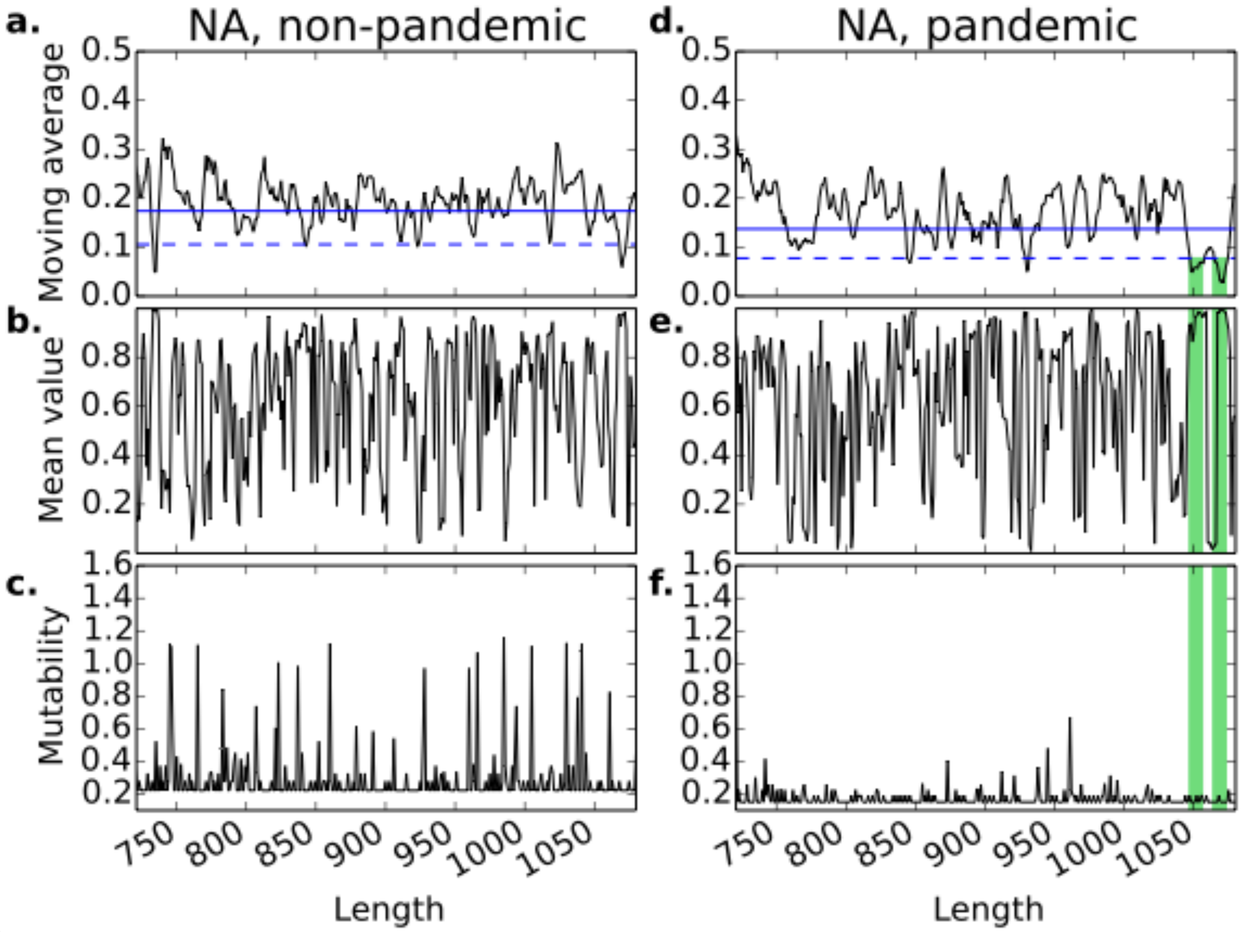

**Supplementary Figure 33**

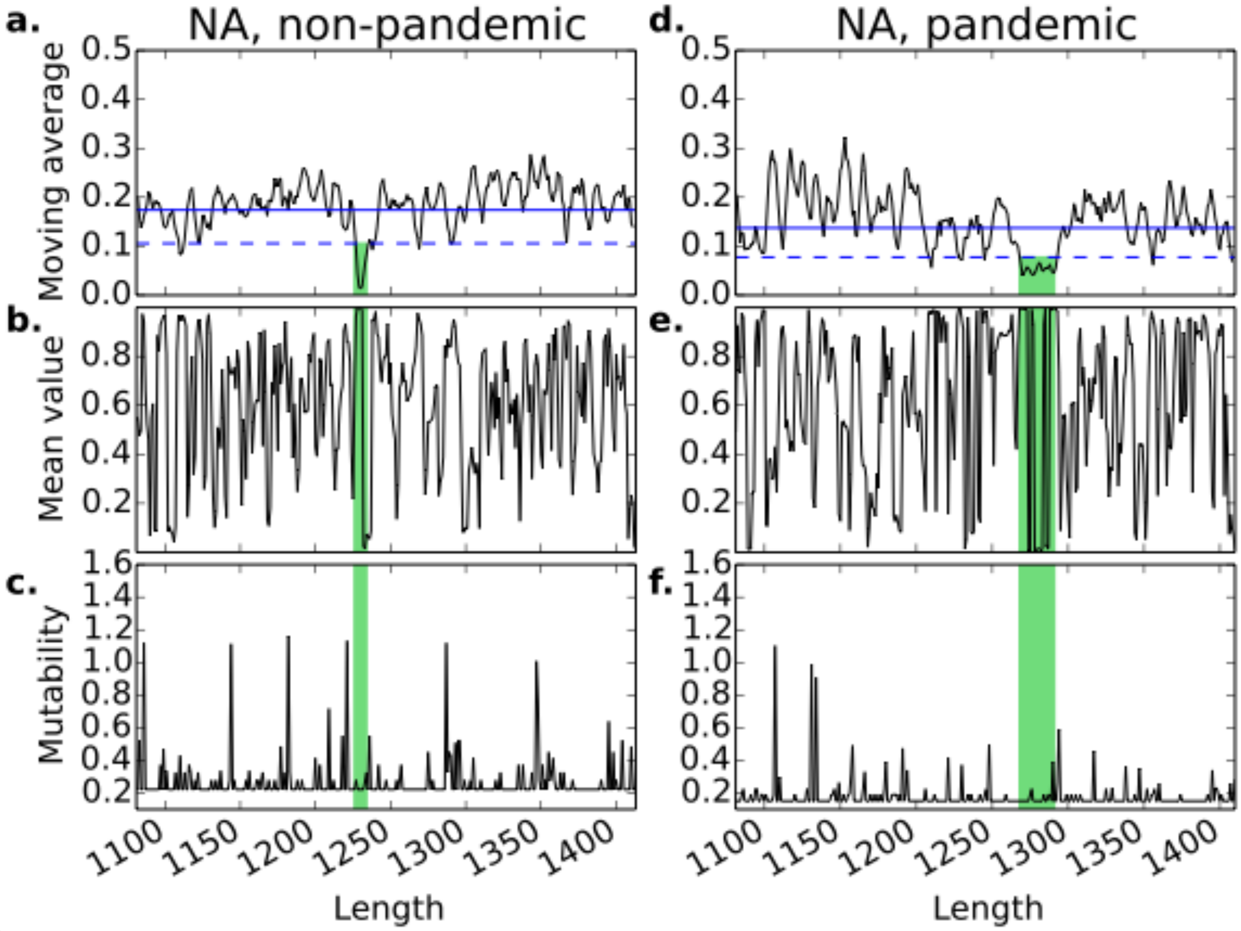

**Supplementary Figure 34**

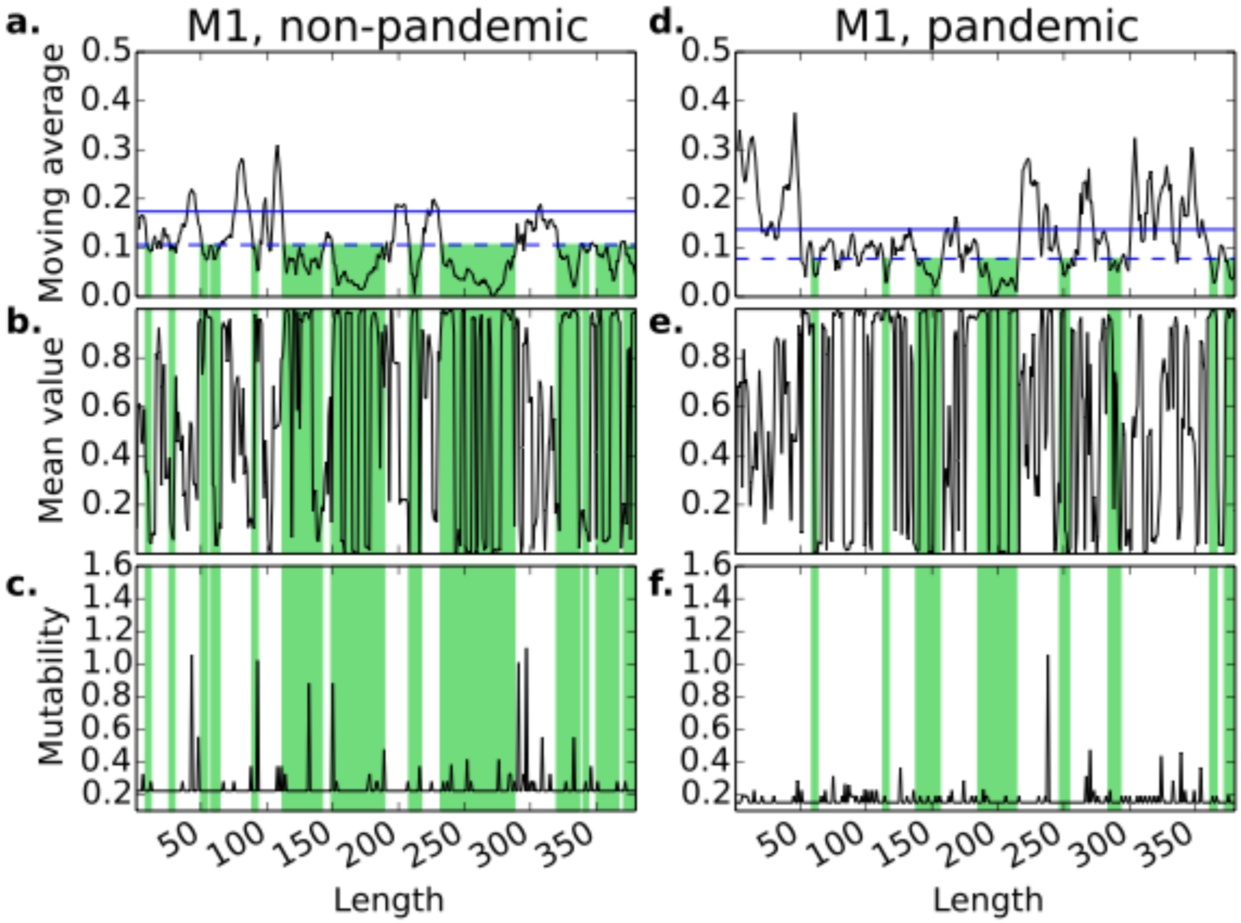

**Supplementary Figure 35**

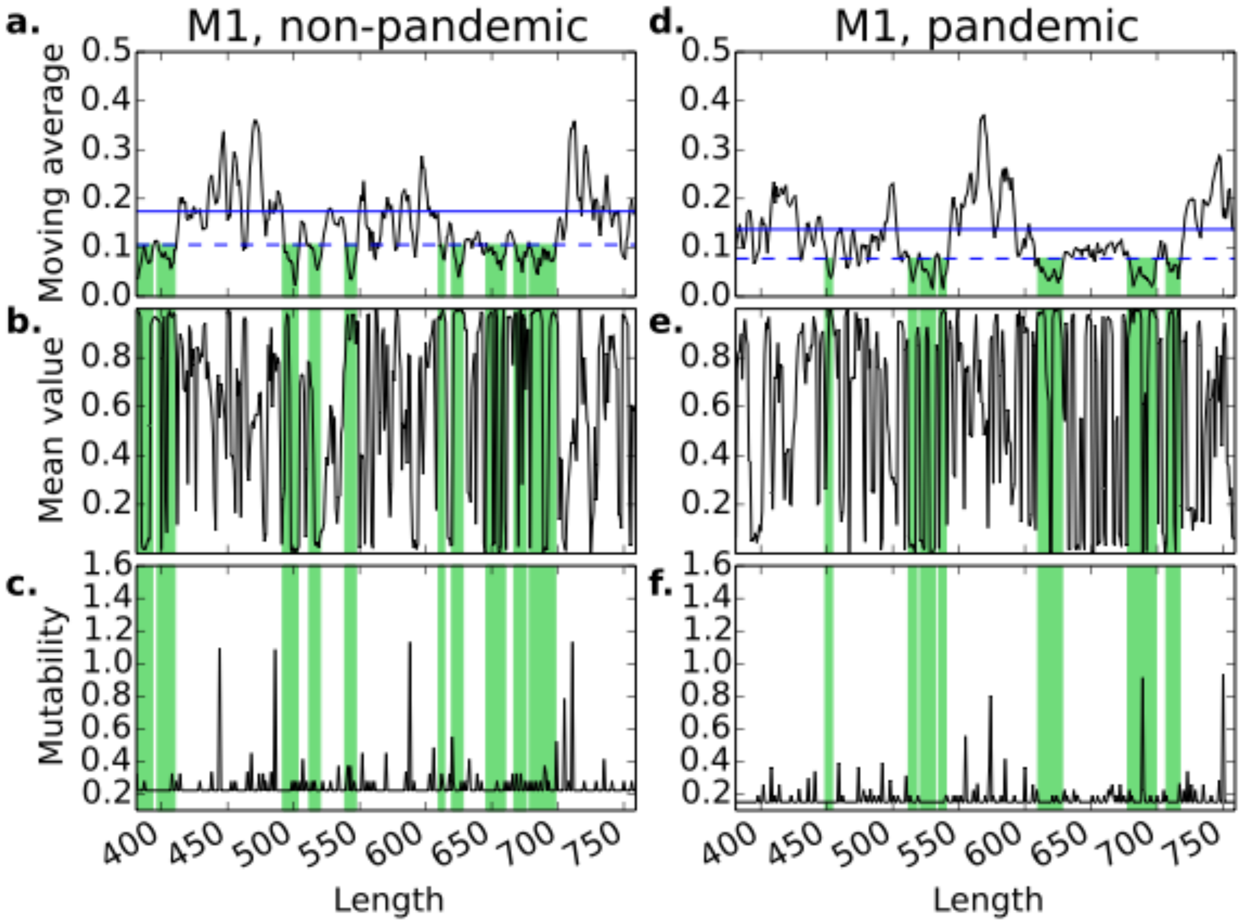

**Supplementary Figure 36**

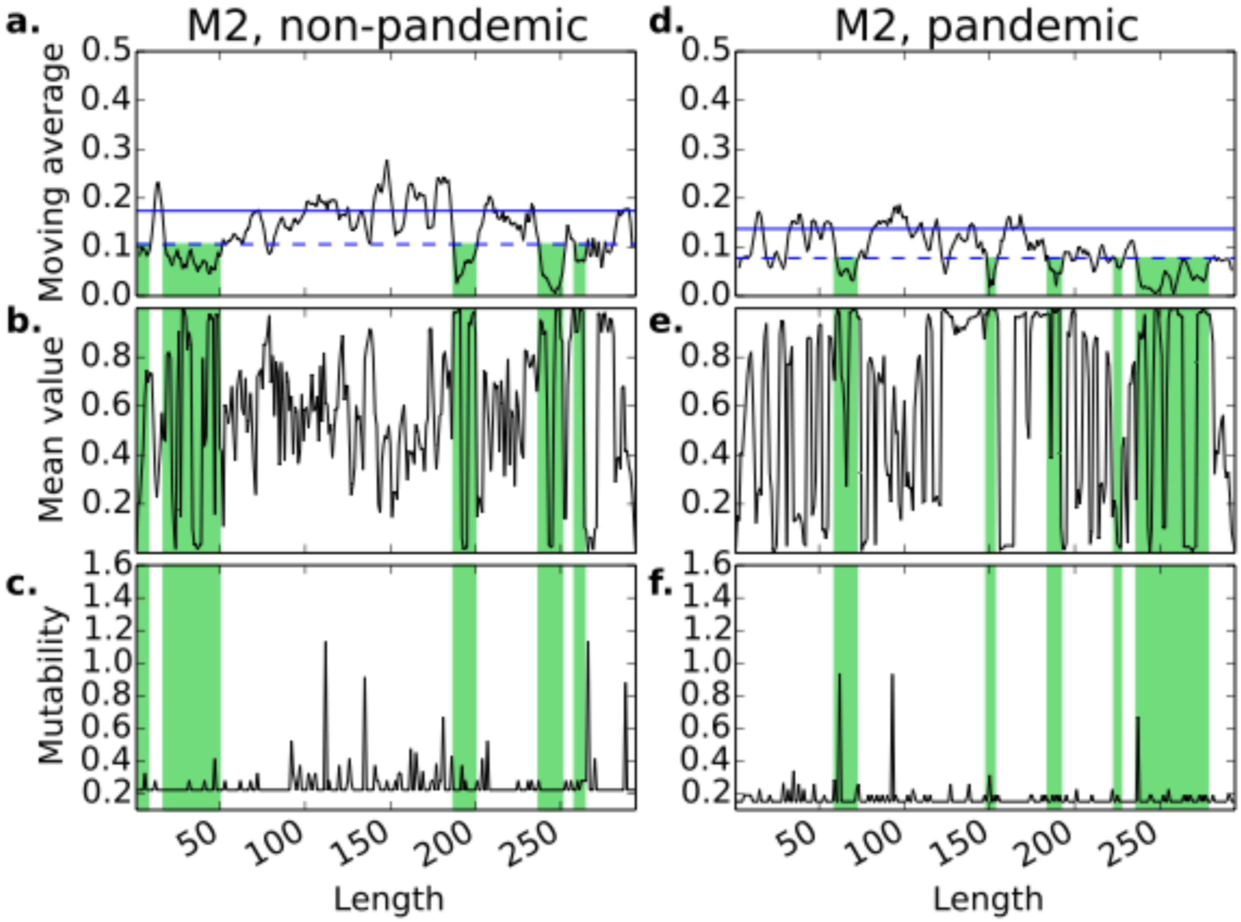

**Supplementary Figure 37**

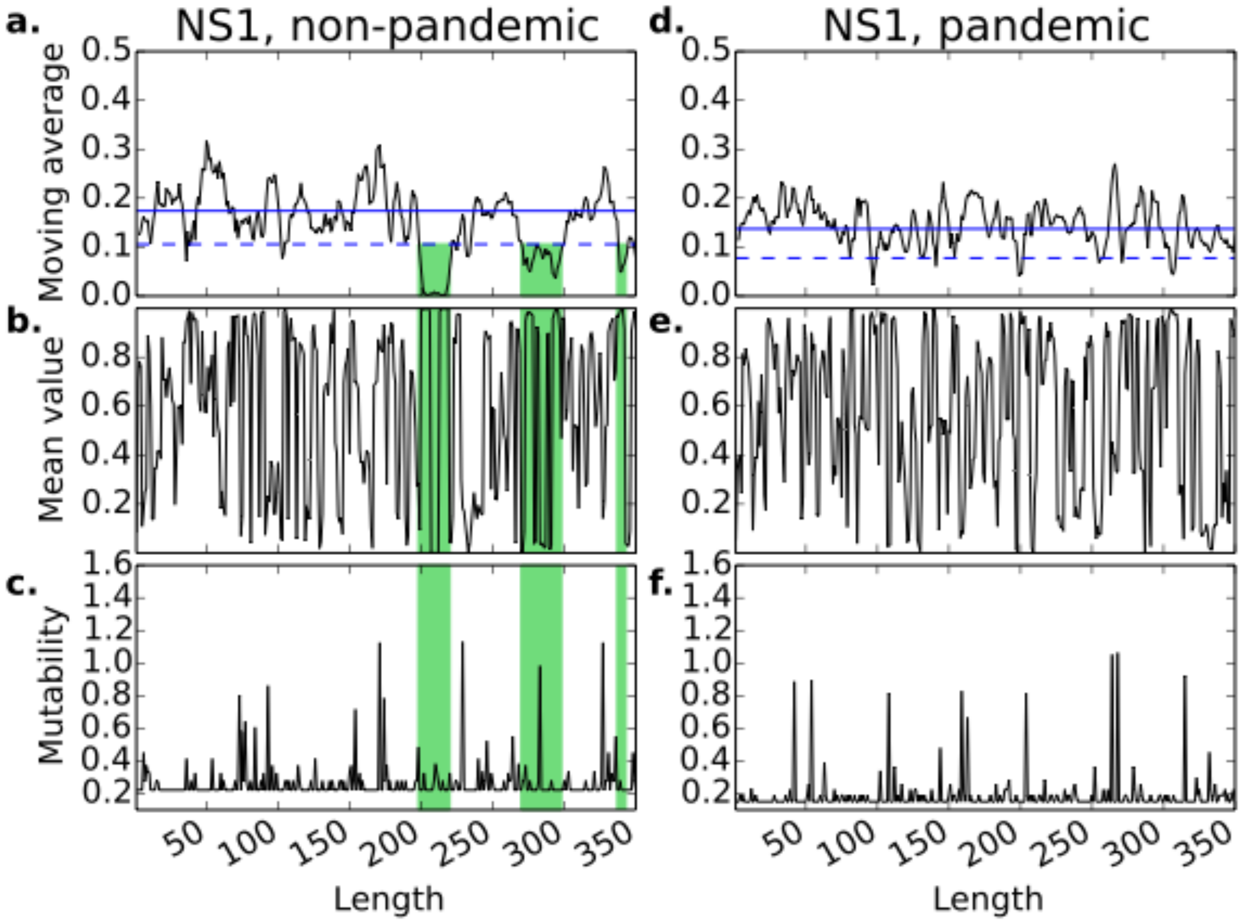

**Supplementary Figure 38**

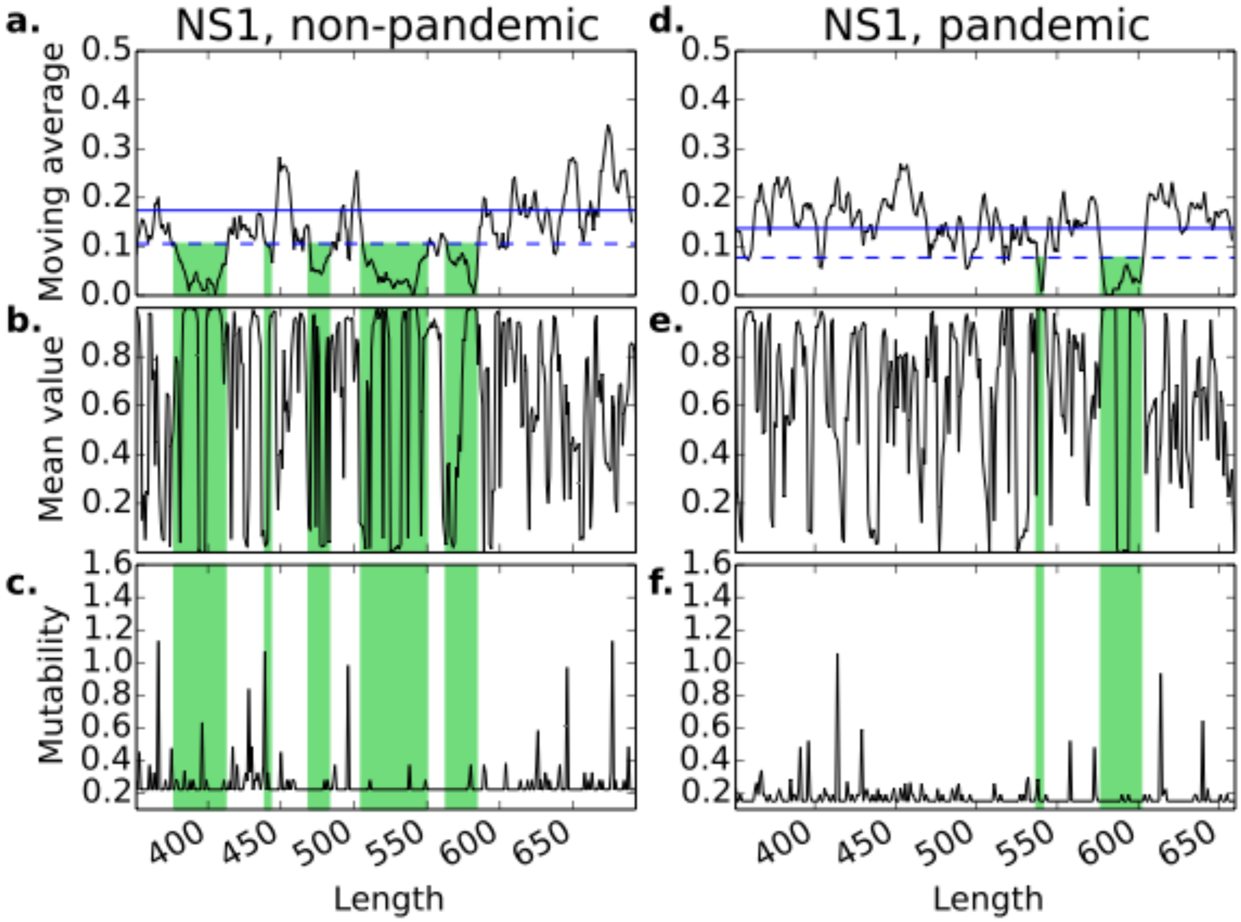

**Supplementary Figures 39-58:** Associations between mutability value for every nucleotide position (position mutability) and corresponding value of moving average of individual standard deviations of the probabilities of nucleotides to be paired (structure variability). X represents structure variability, while Y represents position mutability. The observed correlations are very low, which demonstrates the absence of relationship between these parameters.

**Supplementary Figure 39**

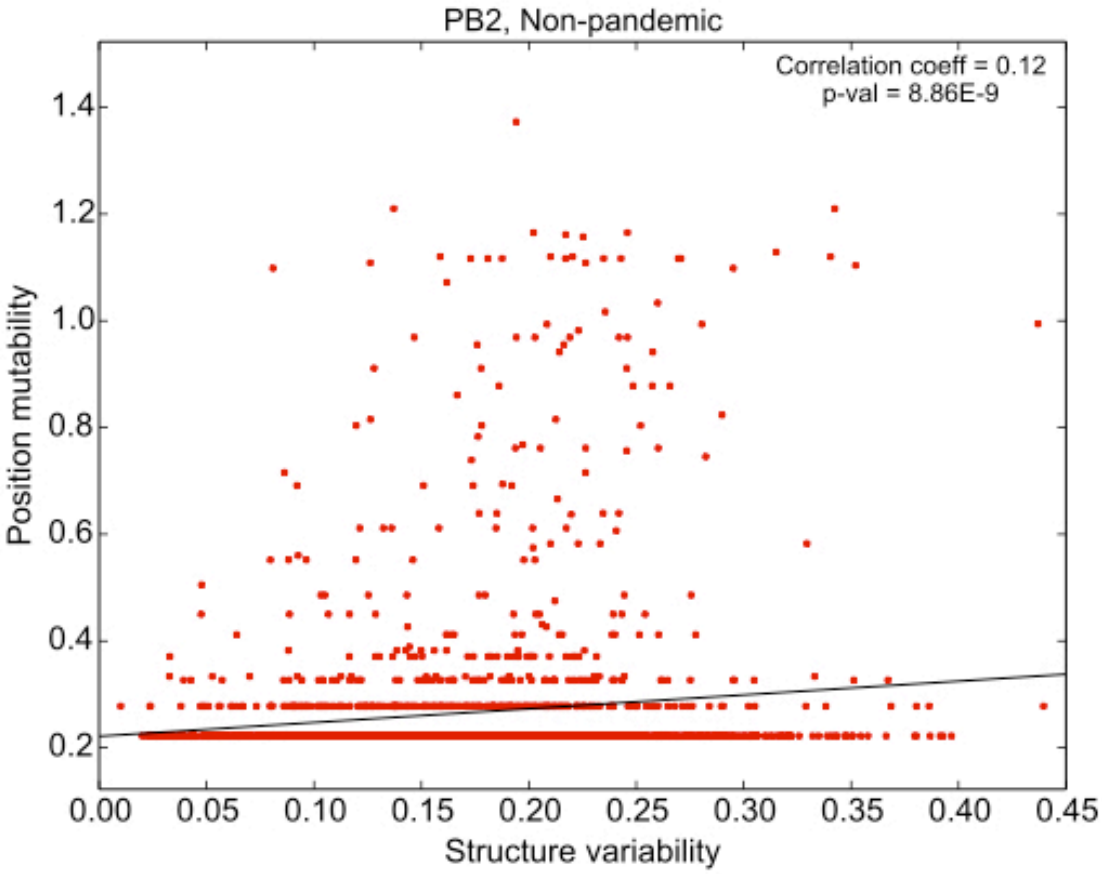

**Supplementary Figure 40**

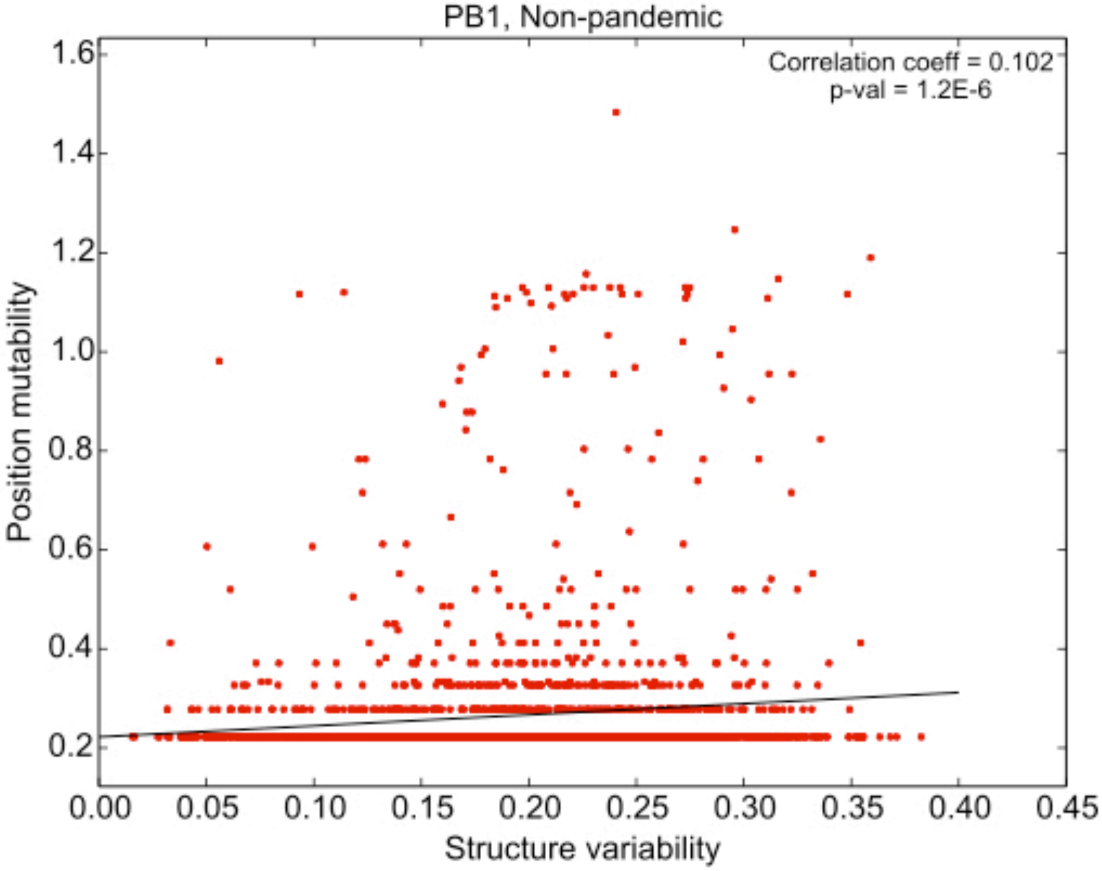

**Supplementary Figure 41**

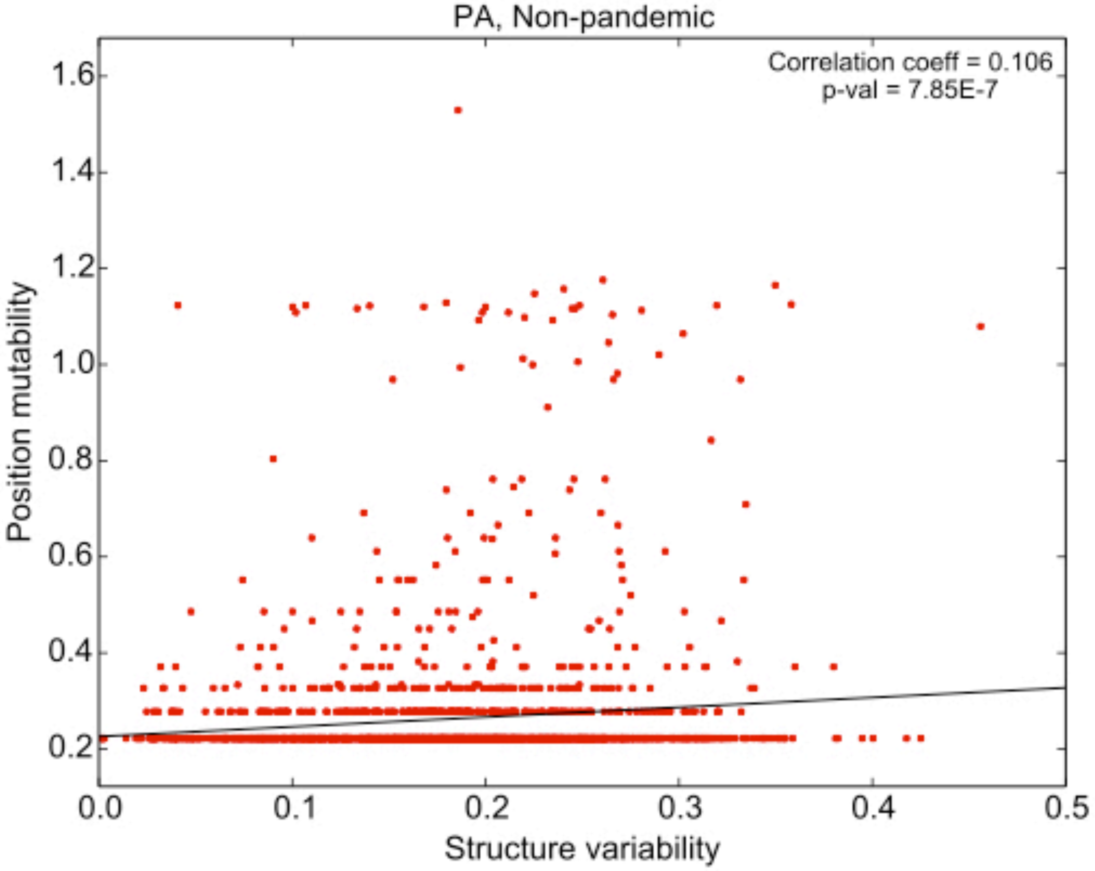

**Supplementary Figure 42**

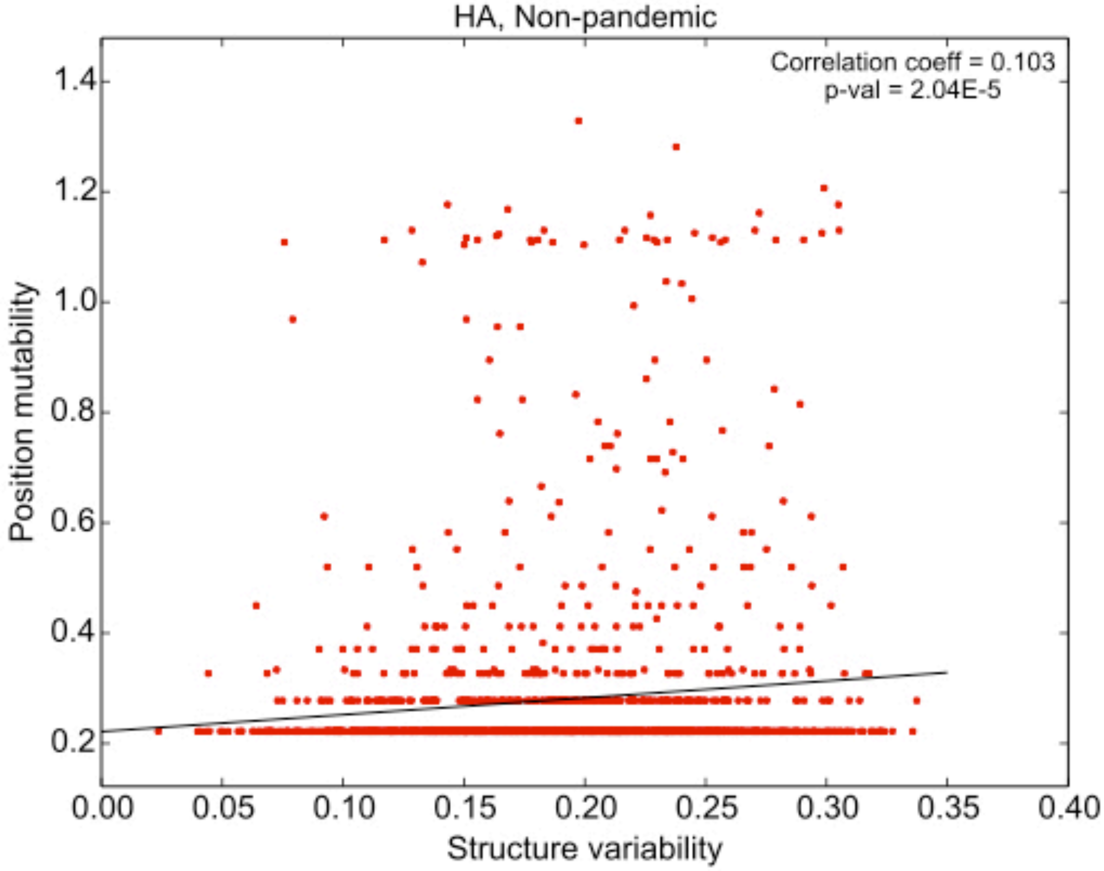

**Supplementary Figure 43**

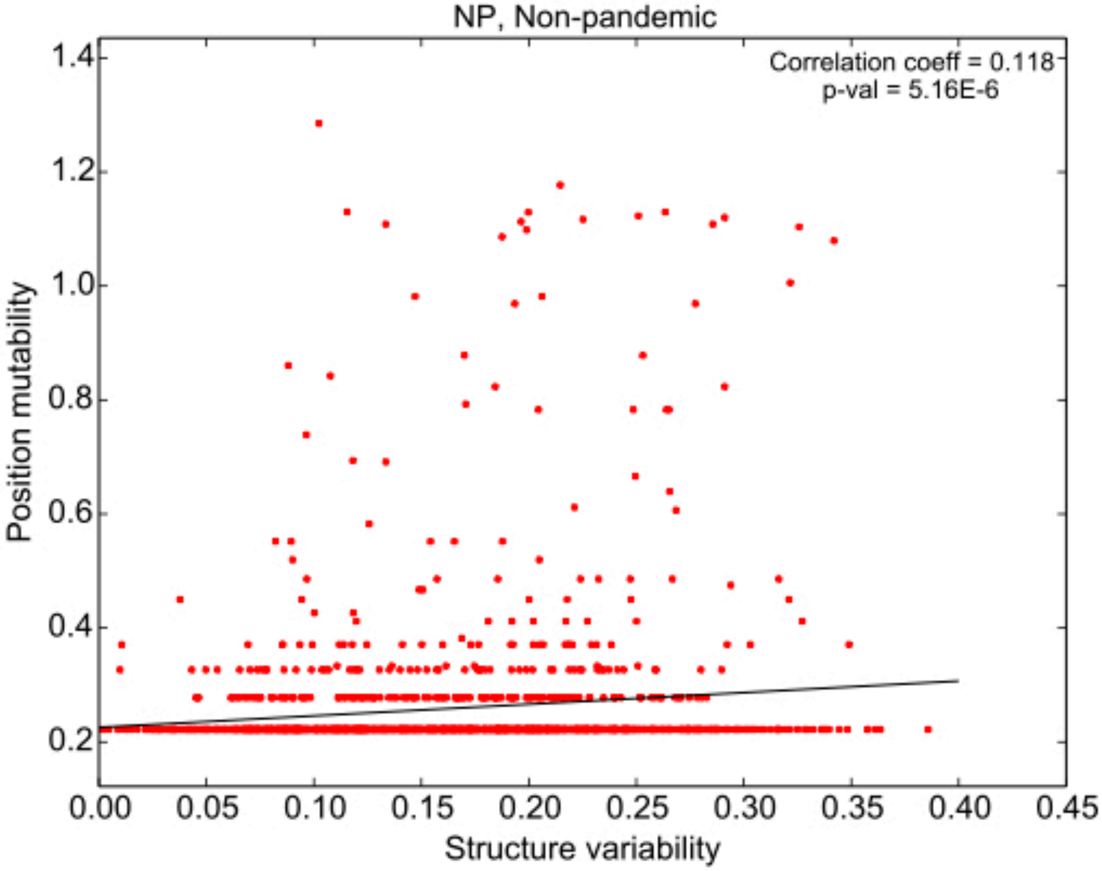

**Supplementary Figure 44**

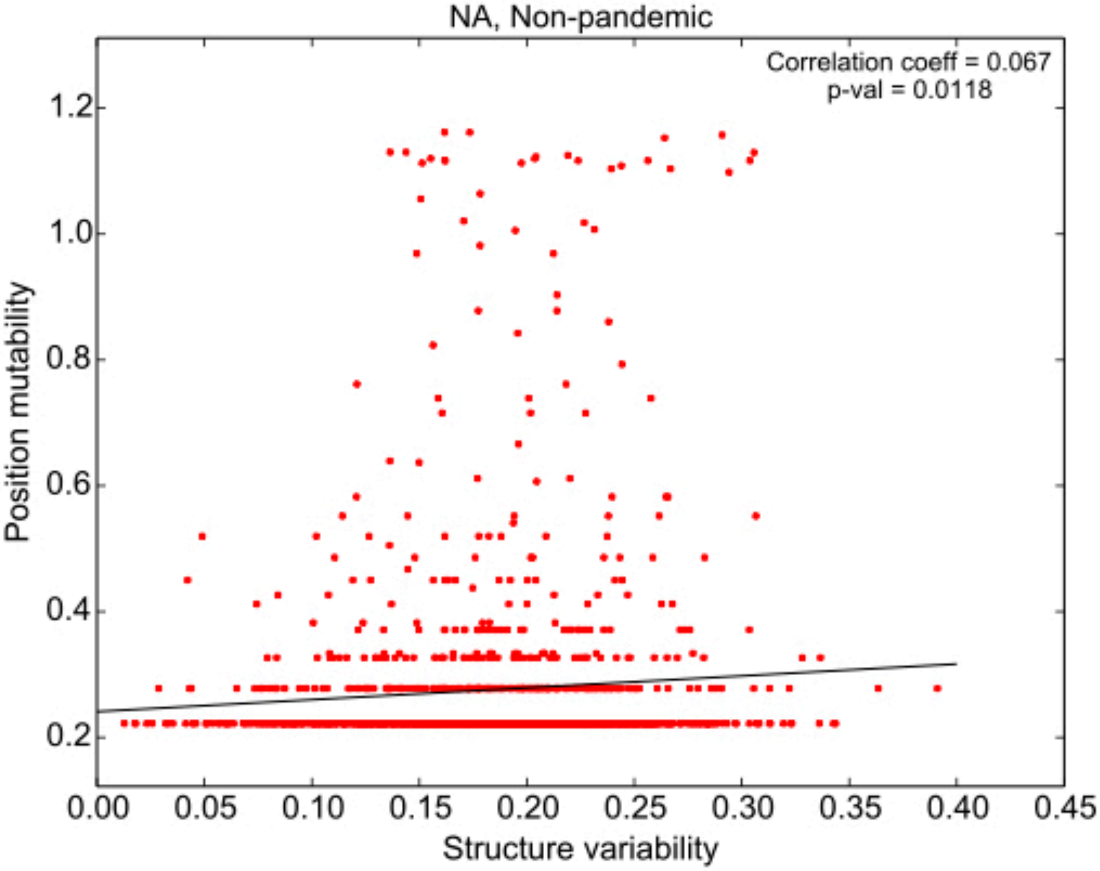

**Supplementary Figure 45**

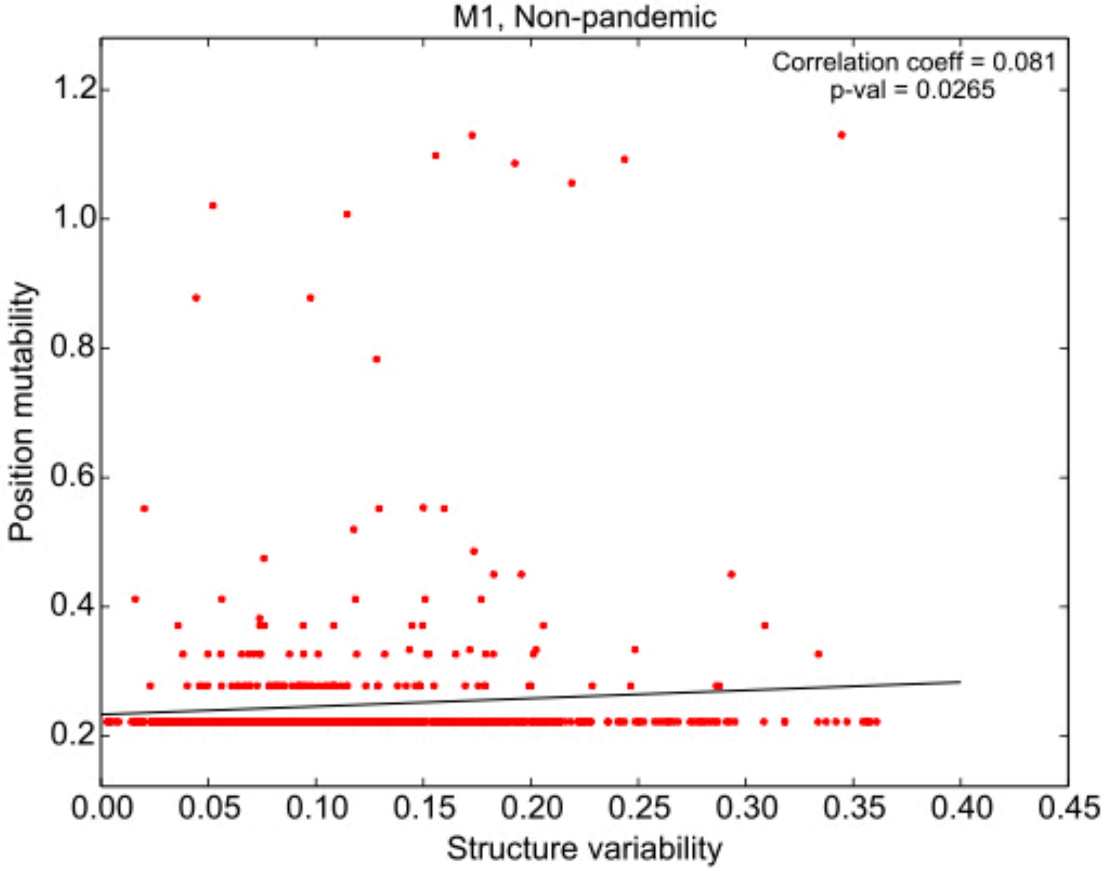

**Supplementary Figure 46**

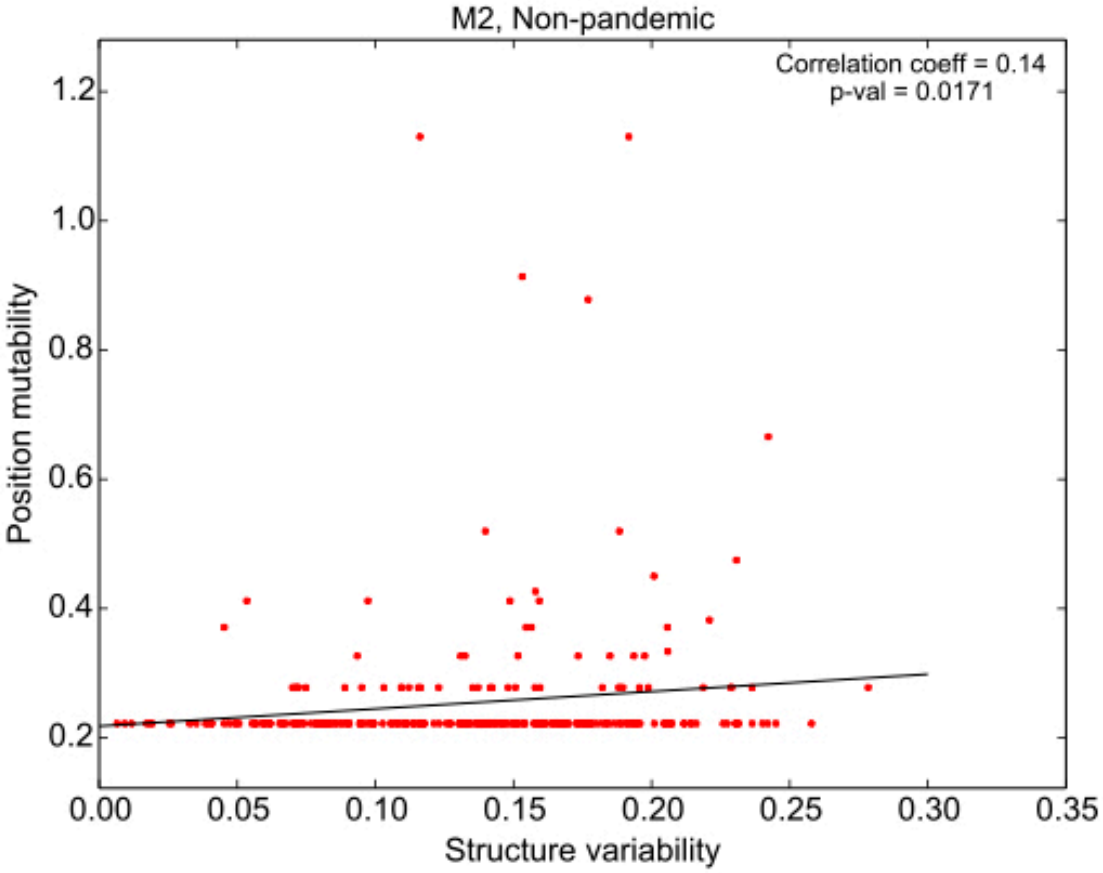

**Supplementary Figure 47**

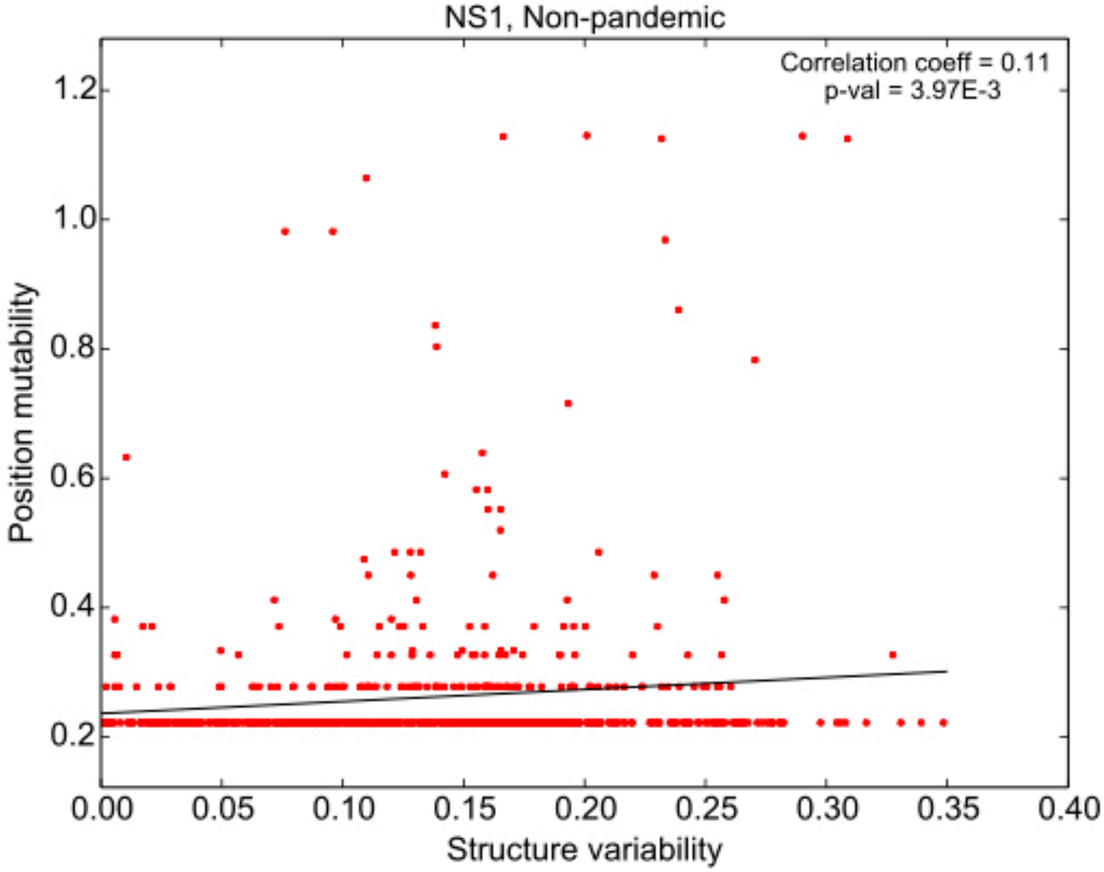

**Supplementary Figure 48**

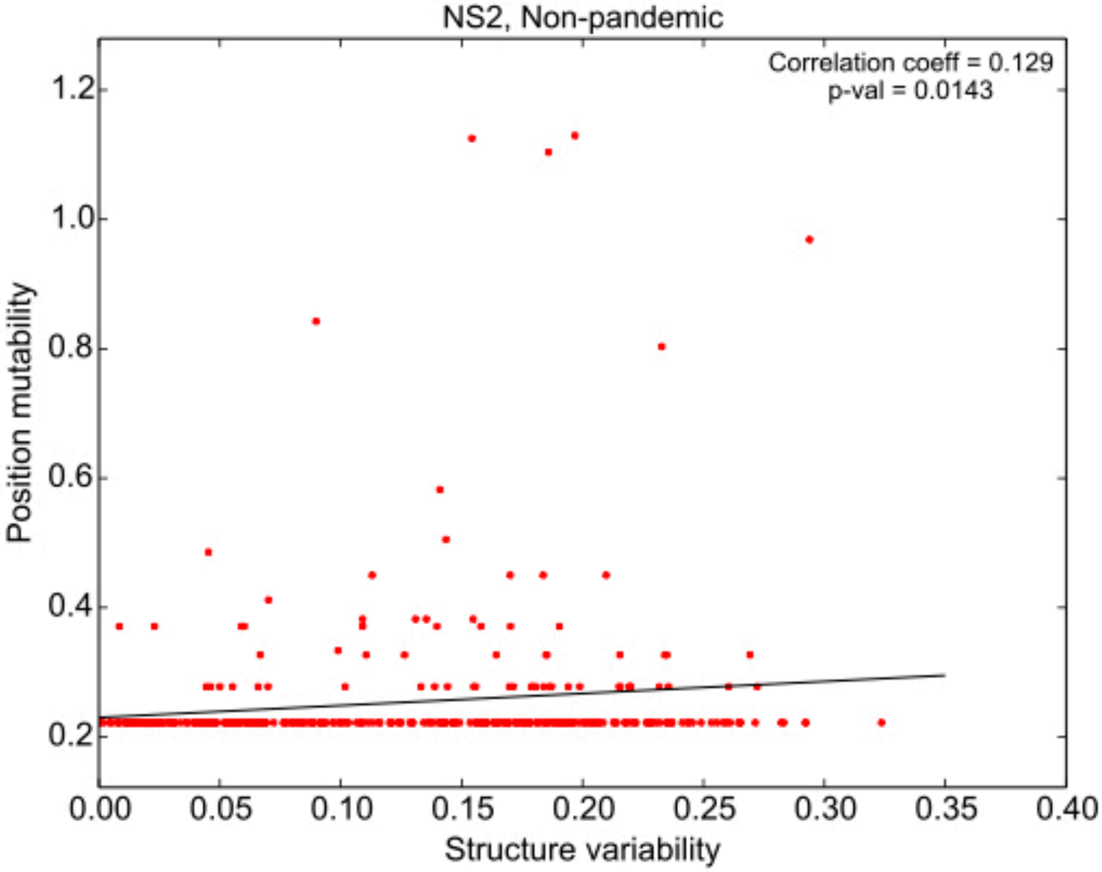

**Supplementary Figure 49**

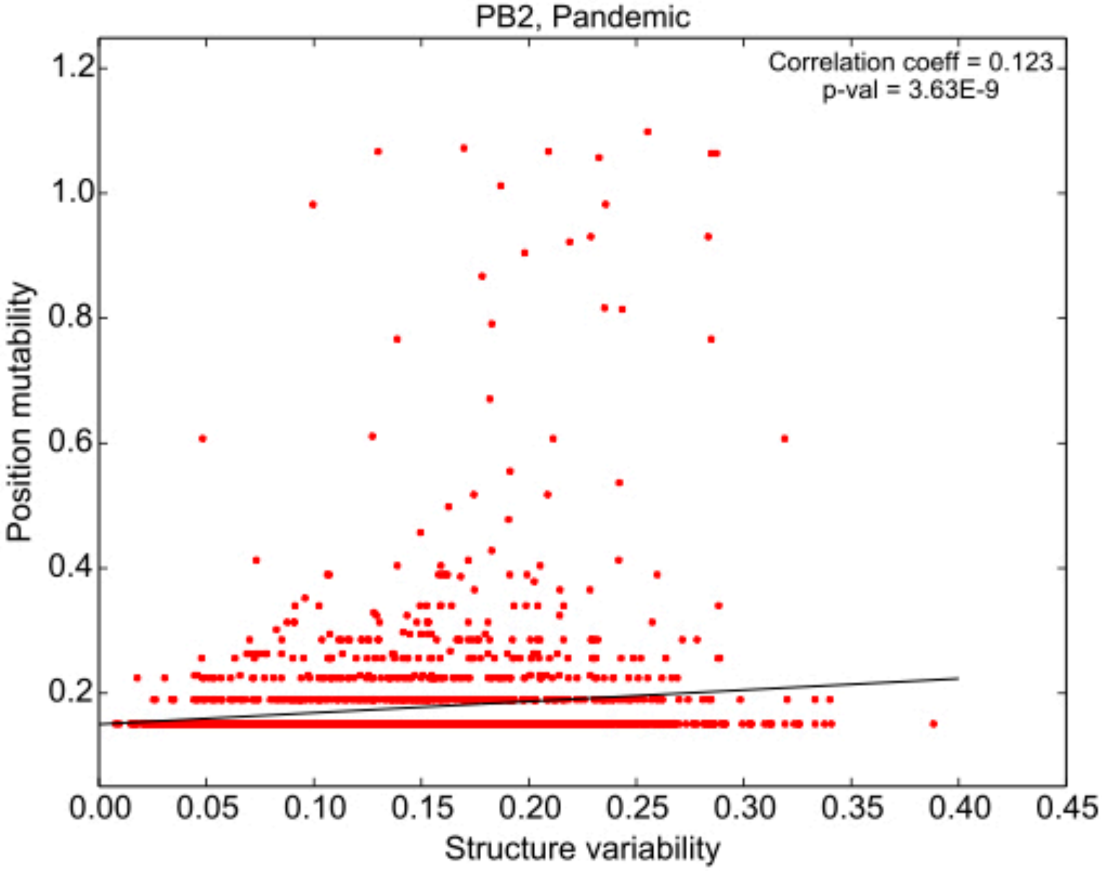

**Supplementary Figure 50**

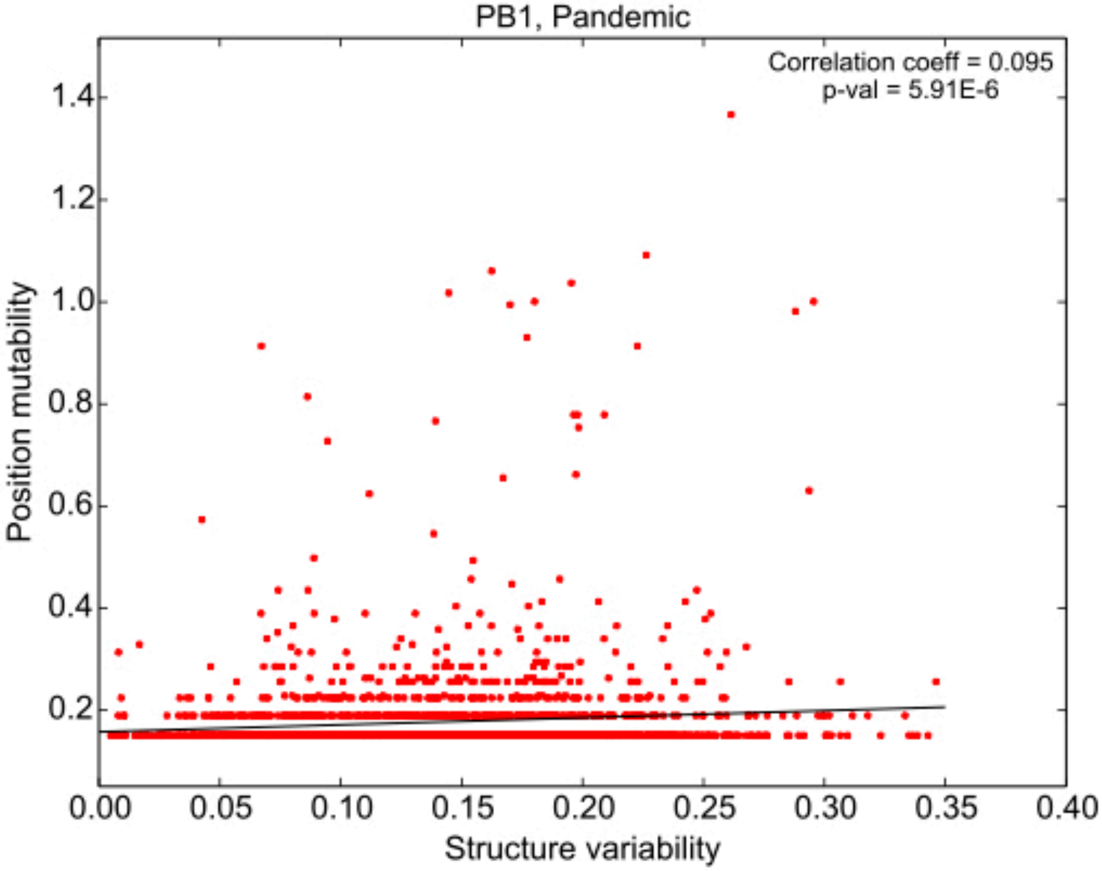

**Supplementary Figure 51**

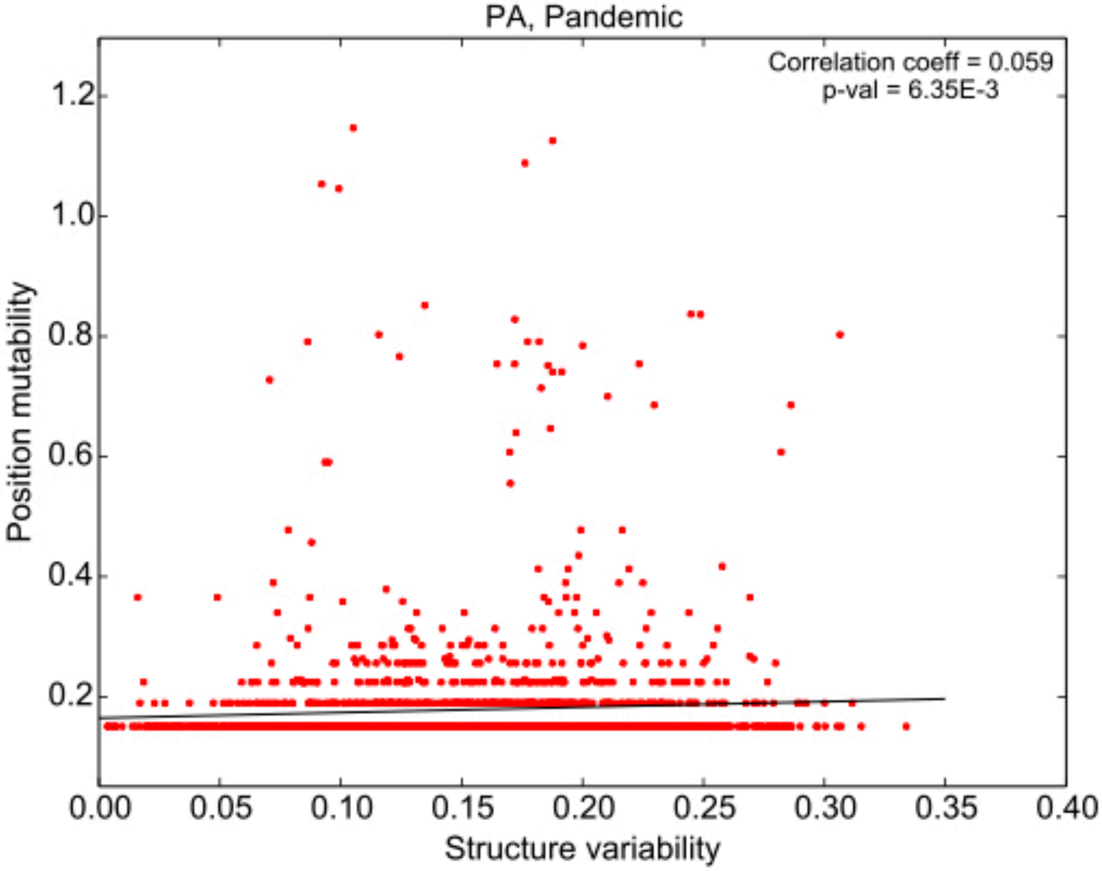

**Supplementary Figure 52**

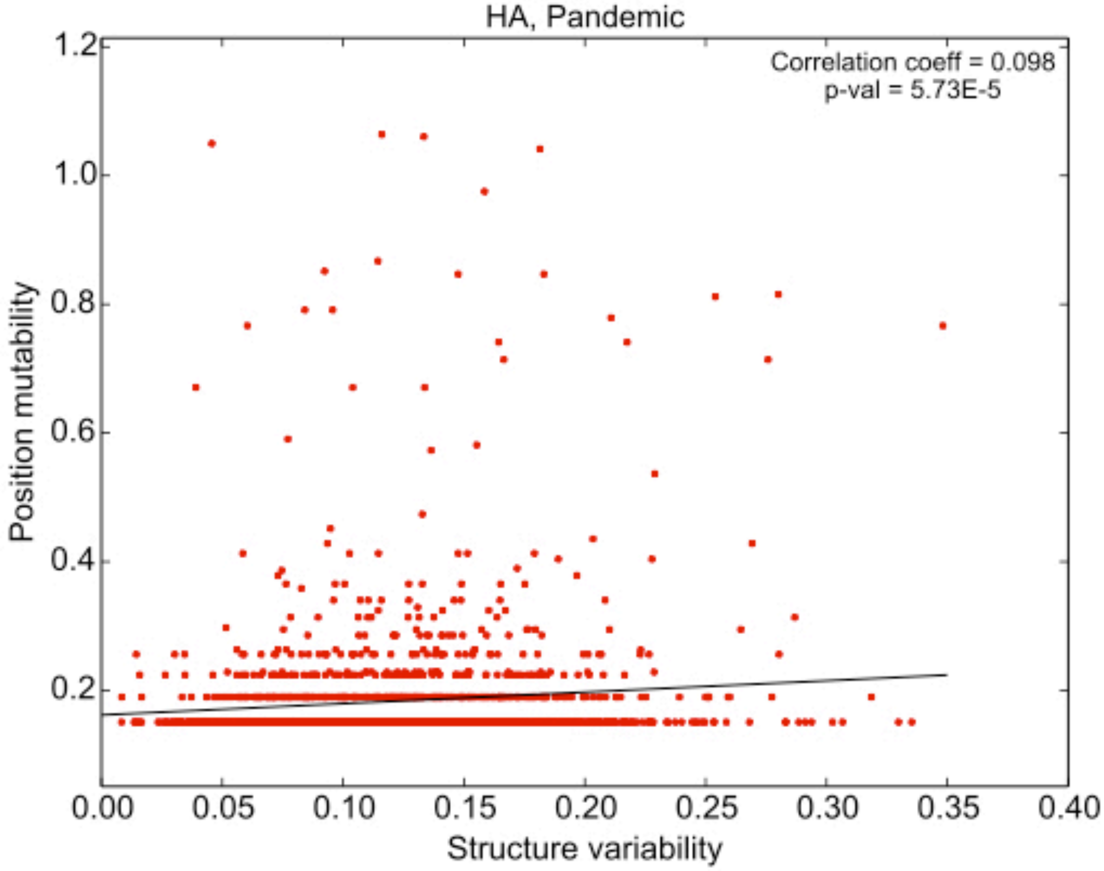

**Supplementary Figure 53**

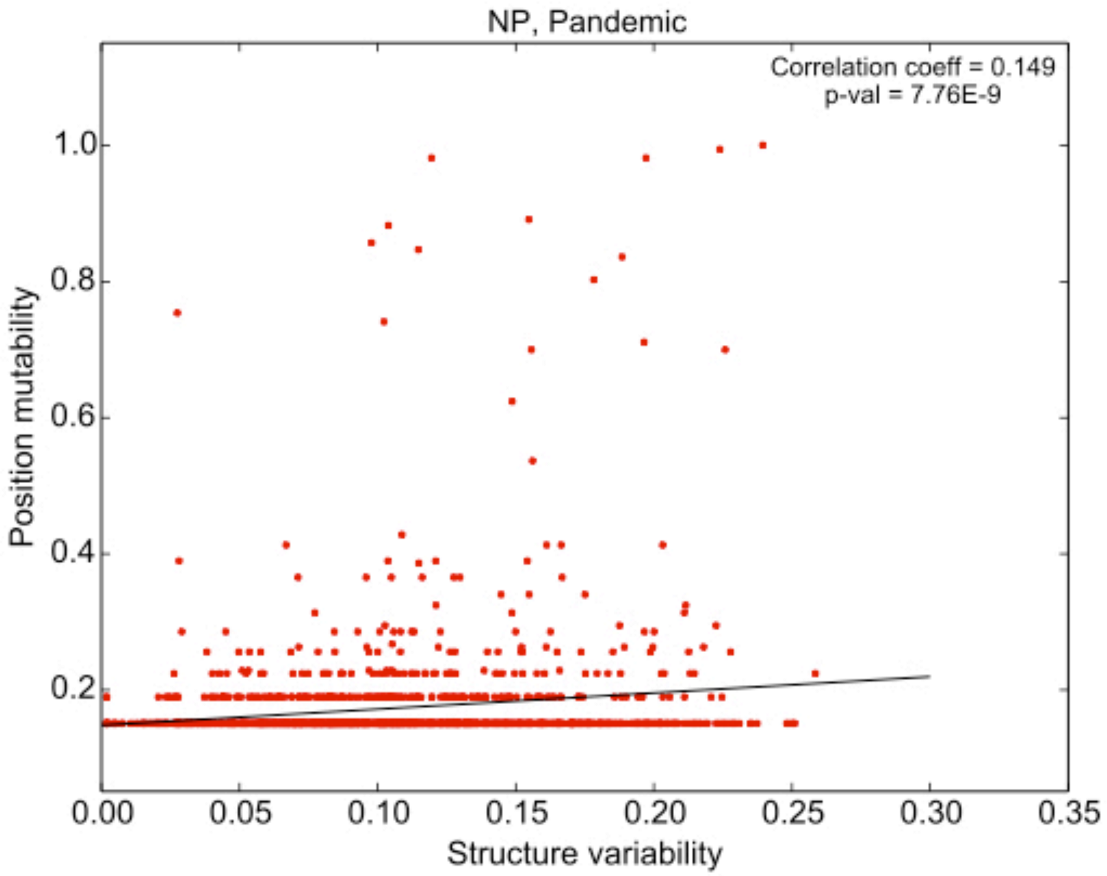

**Supplementary Figure 54**

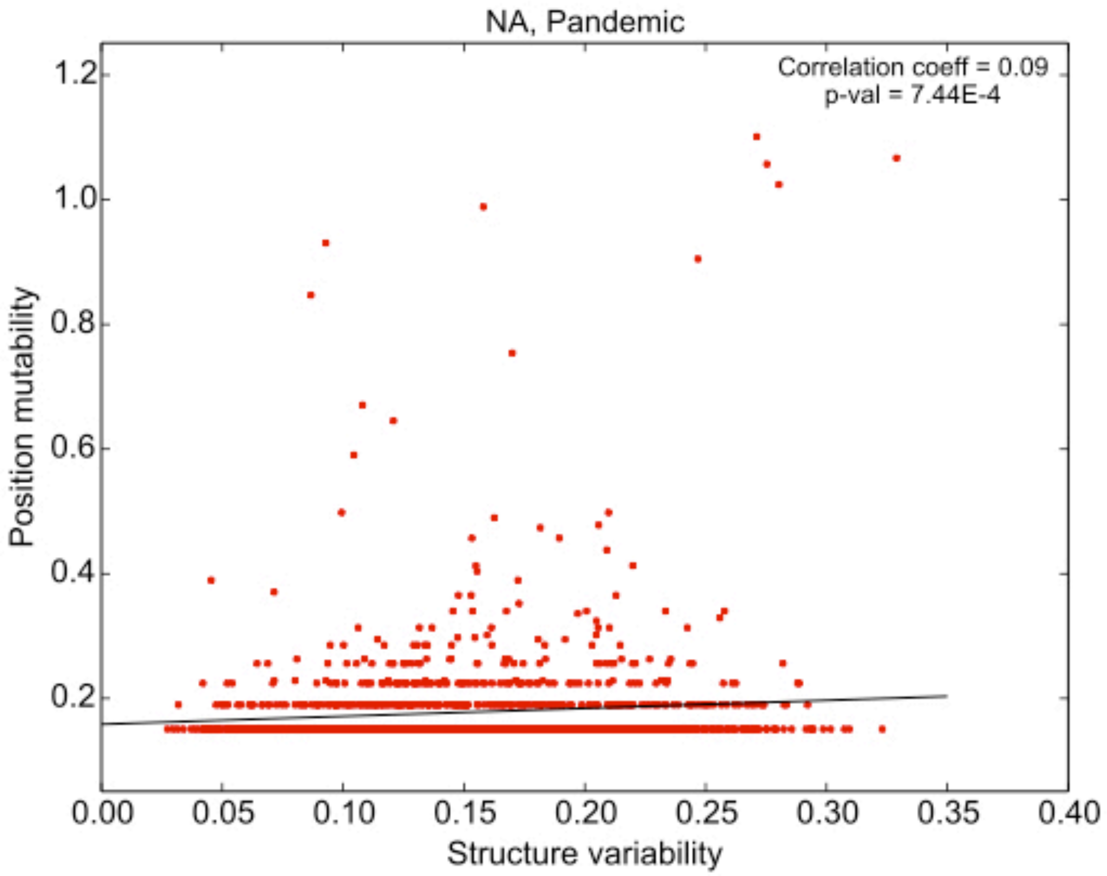

**Supplementary Figure 55**

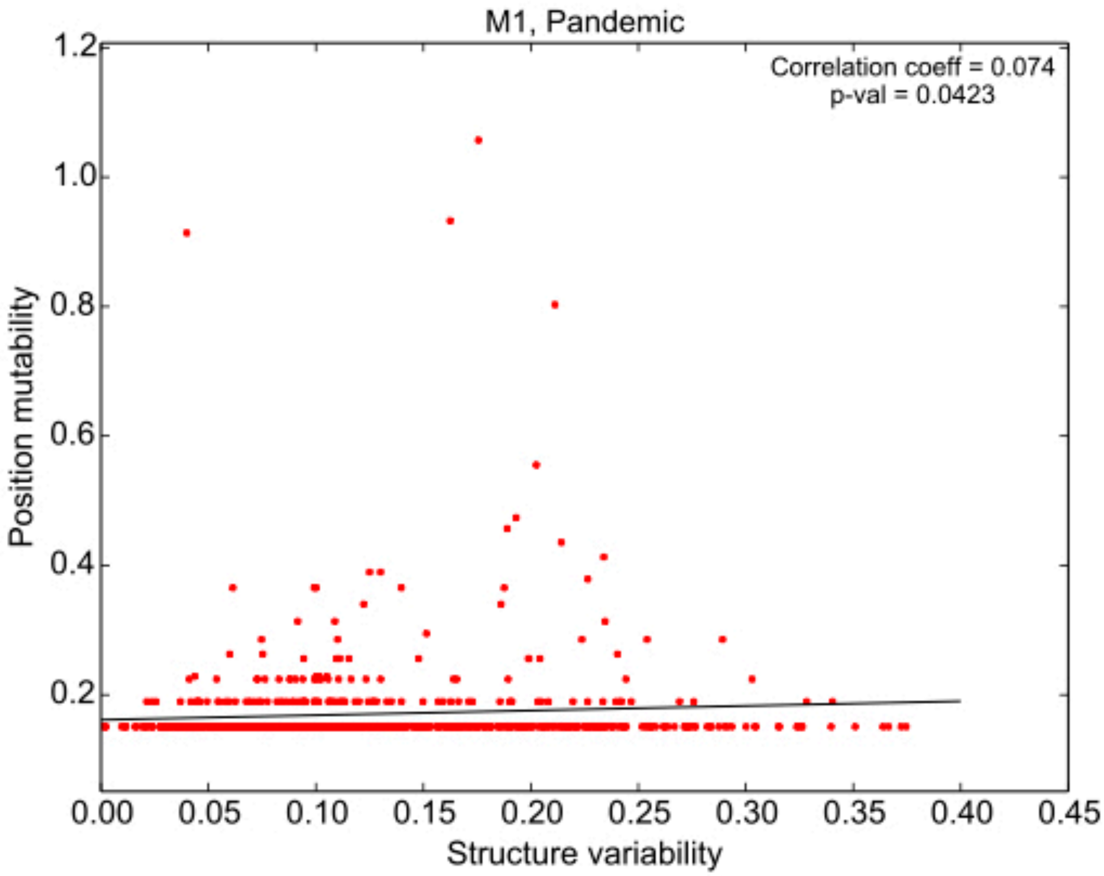

**Supplementary Figure 56**

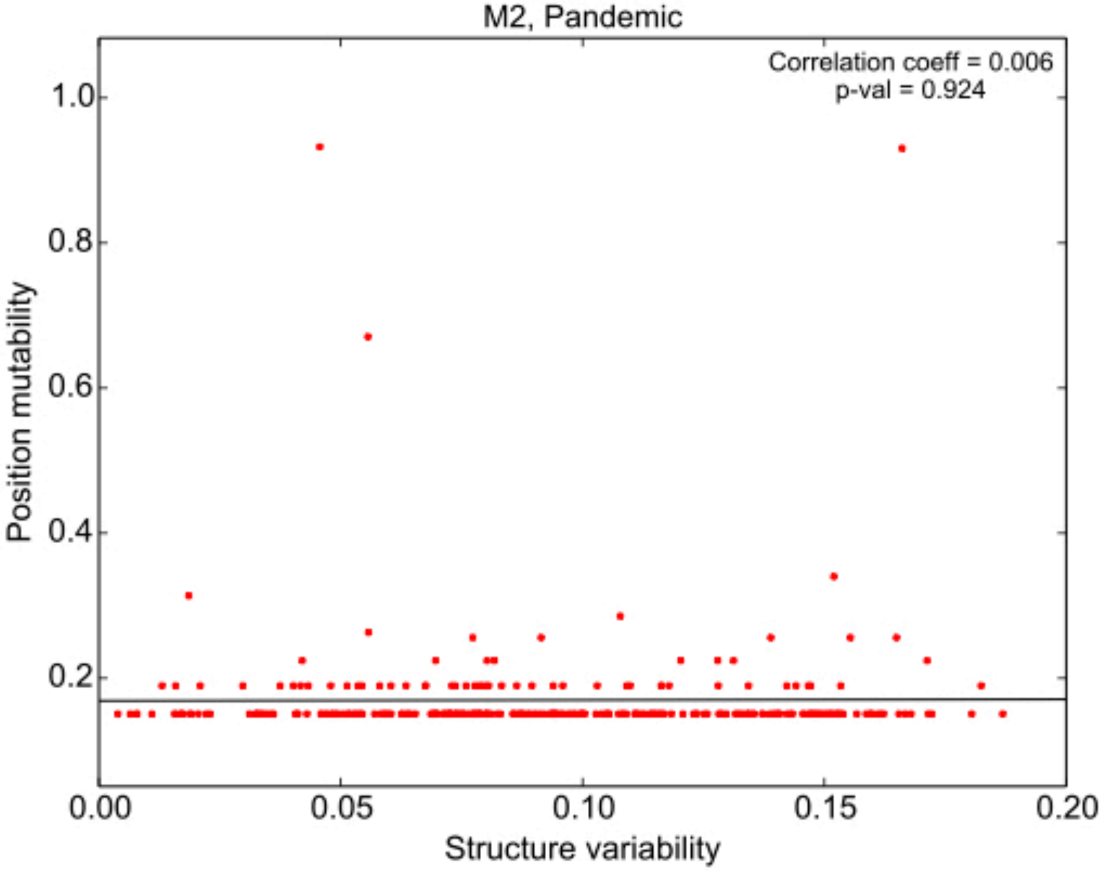

**Supplementary Figure 57**

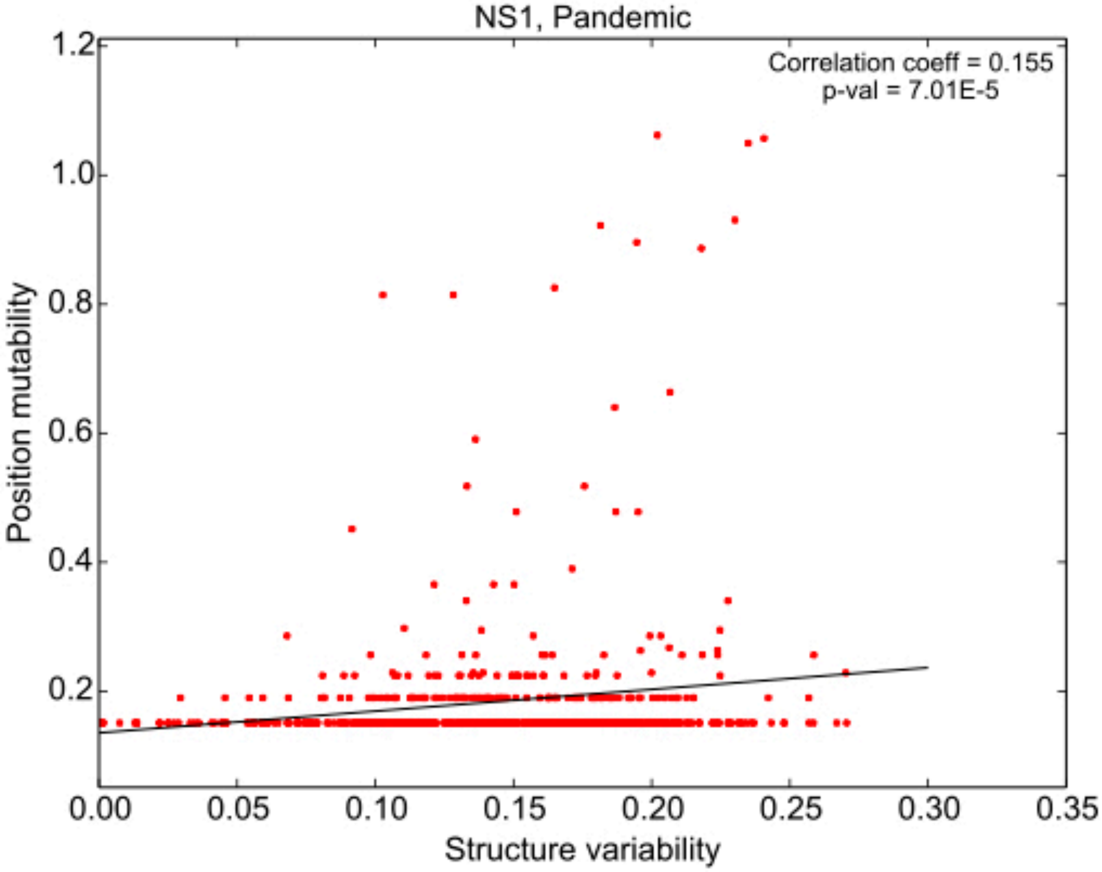

**Supplementary Figure 58**

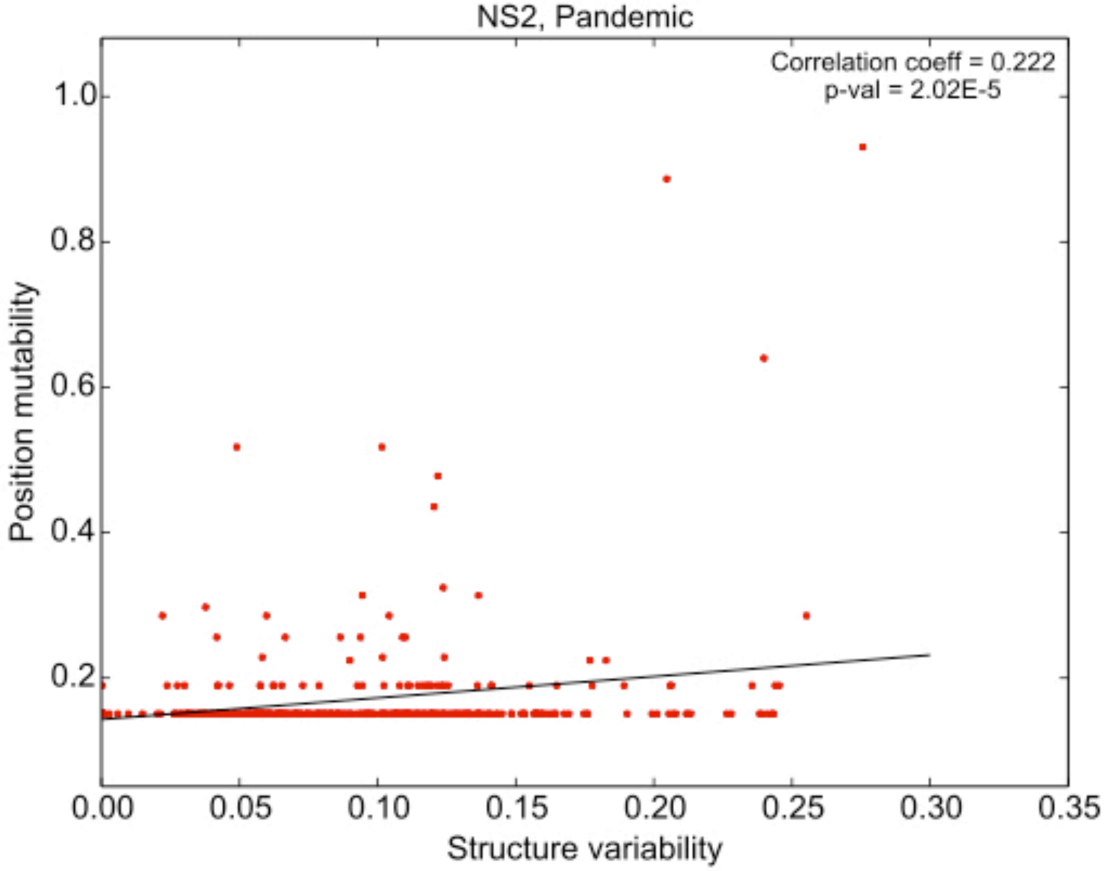
